# High-plex protein imaging across thick tissue by synchronized amplification

**DOI:** 10.64898/2026.10.12.757405

**Authors:** Kuanwei Sheng, Hanquan Su, Gokul Gowri, Maxwell Furman, Matthew Serrata, Lili Du, Charlotte George, Michel Nofal, Ryan McMillan, Kaibo Ma, Xiaokang Lun, Hiroshi Sasaki, Jiyoun Jeong, German Pihan, Peng Yin

## Abstract

High-plex protein imaging is essential for decoding complex tissue organization, yet it remains largely confined to thin sections. Current approaches in millimeter-thick samples suffer from poor signal penetration and depth-dependent, uneven signal amplification. Even with specialized instrumentation to facilitate probe diffusion, iterative millimeter-thick tissue workflows can take more than four days per staining round, and reported eleven-marker experiments extend beyond a month. We introduce Amplification by Cyclic Extension (ACE), a DNA-barcoded, synchronized *in situ* amplification strategy that uses thermally controlled cyclic extension to coordinate DNA concatemer growth across tissue depths. By synchronously activating and coordinating reaction progression throughout the sample, ACE enables controlled, uniform signal amplification across tissue depths. Using ACE, we demonstrate 63-plex protein and RNA imaging in thin sections, 22-plex protein imaging in 200 µm mouse brain sections, and 16-plex protein imaging in 1 mm-thick brain tissue encompassing over three million cells. For millimeter-thick tissues, ACE reduces per-round target-switching time from multiple days to less than eight hours, shifting volumetric high-plex protein imaging from months to days. Together, these results establish ACE as a synchronized amplification strategy that enables scalable high-plex protein imaging in thick tissues.

## INTRODUCTION

Understanding the spatial distribution of biomolecules, especially proteins, is fundamental for elucidating biological processes. Traditional approaches have been largely confined to <20 µm sections, which limits the study of multicellular systems where key biological features, such as neuronal connectivity, occur across millimeter to organ scales. Advances in tissue clearing now enable volumetric imaging from a few hundred µm up to whole organs or entire organisms ^1-6^. While indirect immunofluorescence remains a primary method for protein imaging, multiplexing is typically limited by the few host species of primary antibody (Ab) and the spectral overlap of the fluorescence channels. These challenges are further amplified in deep tissue, where diffusion slows and light scattering increases.

Approaches such as color unmixing and AI-based demixing have modestly expanded multiplexity ^7-9^ but depend on predefined models or extensive calibration and have limited validation in thick tissue. Non-fluorescent modalities, including imaging mass cytometry and Raman microscopy, offer higher plex capacity ^10-13^ but can be limited by slow acquisition (see calculation in **Footnote I**^1^), lower sensitivity, or destructive procedures, making them often less suitable for large-volume samples.

Iterative antibody staining can overcome the spectral and antibody-species limitations of fluorescence multiplexing ^14,15^, but it requires multiple rounds of slow primary antibody staining, which typically take hours to overnight even for thin tissue sections, and faces additional technical hurdles in thick specimens. For example, PathoPlex deployed 95 exchange cycles (with one cycle per day), which took over 3 months to image 150 proteins in thin sections ^15^. Commercial microfluidic systems such as Lunaphore COMET ^16^ accelerate Ab exchange, but remain limited by small imaging areas (∼12.5 × 12.5 mm²) and reliance on proprietary equipment, restricting scalability. Further challenges arise as the complexity of the specimen increases. For instance, the Multiplexed Analysis of Proteins approach demonstrated 12-plex imaging in 100 µm-thick hydrogel-cleared tissues, but each staining round necessitated extended Ab incubation and exchange cycles spanning approximately 2–3 days ^17^. mELAST images two markers per round and requires about 4-5 days per round, resulting in over one month to acquire 11 markers in 1 mm thick tissue, even with gel embedding and physically assisted reagent diffusion through tissue stretching (see calculation in **Footnote II^2^**) ^4,17^.

To overcome this iterative Ab staining bottleneck, DNA-Exchange based imaging methods stain all the protein targets in one step using DNA-barcoded primary Abs, followed by rapid cycles of DNA imager exchange to allow accelerated sequential imaging ^18-20^. Leveraging the programmability and small molecular size of DNA (∼6 kDa, ∼25-fold smaller than Abs), these methods enable order-of-magnitude increases in usable probe concentration and binding kinetics, and substantially faster diffusion (see calculation in **Footnote III**^3^). Although effective for imaging relatively abundant targets in cells and thin tissue sections, early implementations of DNA-Exchange–based imaging, including commercially available CODEX, lack signal amplification, limiting sensitivity and posing challenges in thick tissues where light scattering and signal attenuation are pronounced.

To achieve signal amplification, both *ex situ* and *in situ* amplifier synthesis strategies have been developed. *Ex situ* approaches, including SABER-FISH^21^, immuno-SABER ^22^ and CosMx SMI ^23^, use pre-synthesized DNA structures (e.g., long concatemers or branched DNA) as signal amplifiers. However, the non-specific binding of long DNA strands may increase background, and their substantial size hinders effective penetration in thick tissue. *In situ* amplification such as Rolling Circle Amplification (RCA) is widely used for RNA ^24-26^, but protein applications like Cassini ^27^ can suffer from unregulated DNA nanoball growth (>100 nm), leading to resource depletion and loss of subcellular detail. Hybridization Chain Reaction (HCR) ^28-33^ has enabled RNA and protein imaging in thin sections, and RNA detection in thick tissues, including RNA at the whole-brain scale ^34-36^. However, uniform amplification of protein in mm-scale tissue remains to be demonstrated ^37^. High-plex imaging with HCR, due to the limited number of orthogonal amplifiers, relies on repeated cycles of probe binding, amplification, and concatemer/fluorophore removal ^33,36^. In thick tissues, each cycle is slowed by limited diffusion and requires prolonged incubations and harsh washes, often extending over several days and compromising target retention. To improve throughput, cycleHCR introduced split-barcode encoding, which allows one-step staining of RNA targets using primary FISH probes, but still requires iterative rounds of HCR amplification and removal ^6^. This approach has enabled imaging of > 200 RNA targets, yet protein demonstrations remain limited (15 targets in < 60 µm thick sections). Despite acceleration relative to conventional HCR, cycleHCR still relies on iterative amplification, which can become progressively slower and less practical as tissue thickness increases. Moreover, compared with relatively sparse RNA targets, protein targets are orders of magnitude more abundant. Because HCR relies on locally initiated, self-propagating chain-growth amplification whose initiation and progression are not synchronized across targets, amplification can rapidly deplete local resources for dense protein targets in thick tissues, leading to uneven, depth-dependent signal.

To implement such an approach, an effective strategy would feature: (1) one-step high-plex staining followed by one-step high-plex amplification, (2) *in situ* implementation, (3) controlled and synchronized reaction initiation, (4) balanced molecule diffusion and consumption rates achieved with synchronized reaction progression, and (5) efficient exchange of small DNA imagers during imaging following amplifier synthesis. First, one-step high-plex staining and amplification improve workflow efficiency, as the diffusion of barcoded Ab and long DNA strands is time-consuming. Second, *in situ* amplification improves signal-to-noise and penetration in thick tissue. Third, keeping the reaction “off” during reagent delivery allows even diffusion through thick tissue before amplification starts, preventing surface-biased signals. Fourth, component consumption should be slower than diffusion to ensure uniform signal distribution, and synchronous control of amplification avoids local dominance by any single target. Fifth, efficient and mild exchange of short DNA imagers, rather than laborious and harsh stripping of primary antibodies or HCR amplifiers, simplifies workflows, accelerates target-switching cycles, and minimizes sample degradation across iterative imaging rounds, particularly in thick tissues.

Guided by these principles, we envisioned a workflow in which all targets in tissue are stained in a single step with primary probes (primary antibodies for protein targets, and primary FISH probes for RNA targets) modified with short barcode primers, followed by simultaneous high-plex *in situ* extension to generate long concatemer amplifiers in a synchronize and controlled manner, providing scaffolds for subsequent rapid exchange of short imagers. Inspired by the Primer Exchange Reaction ^38^ and the Polymerase Chain Reaction ^39^, we developed Amplification by Cyclic Extension (ACE), which uses stepwise, polymerase-driven, iterative DNA extension regulated by thermal cycling to enable precise on/off control of amplification and synchronized reaction progression for multiplexed in situ amplifier synthesis (**Fig. 1a** and **1c**). ACE allows concurrent amplification of multiple targets, enabling one-pot high-plex staining and one-pot high-plex *in situ* signal amplification (**Fig. 2a**). Short DNA barcodes minimize nonspecific binding, and the compact probes and imagers penetrate thick tissue efficiently. Synchronized initiation is achieved by first extension and then pausing the reaction at low temperature and resetting amplification for the next synchronized extension cycle by dissociation of ssEXT at high temperature. The cyclic control of on and off stages balances molecule diffusion and consumption, allowing even reagent distribution and uniform signal across tissue depths. Using rapid DNA imager exchange, ACE reduces the time for one three-target imaging round on 1-mm mouse brain sections from more than 4 days with prior method ^4^ to less than 8 hours, while eliminating the complex gel embedding and physical compression steps ^4^.

**Figure 1.**
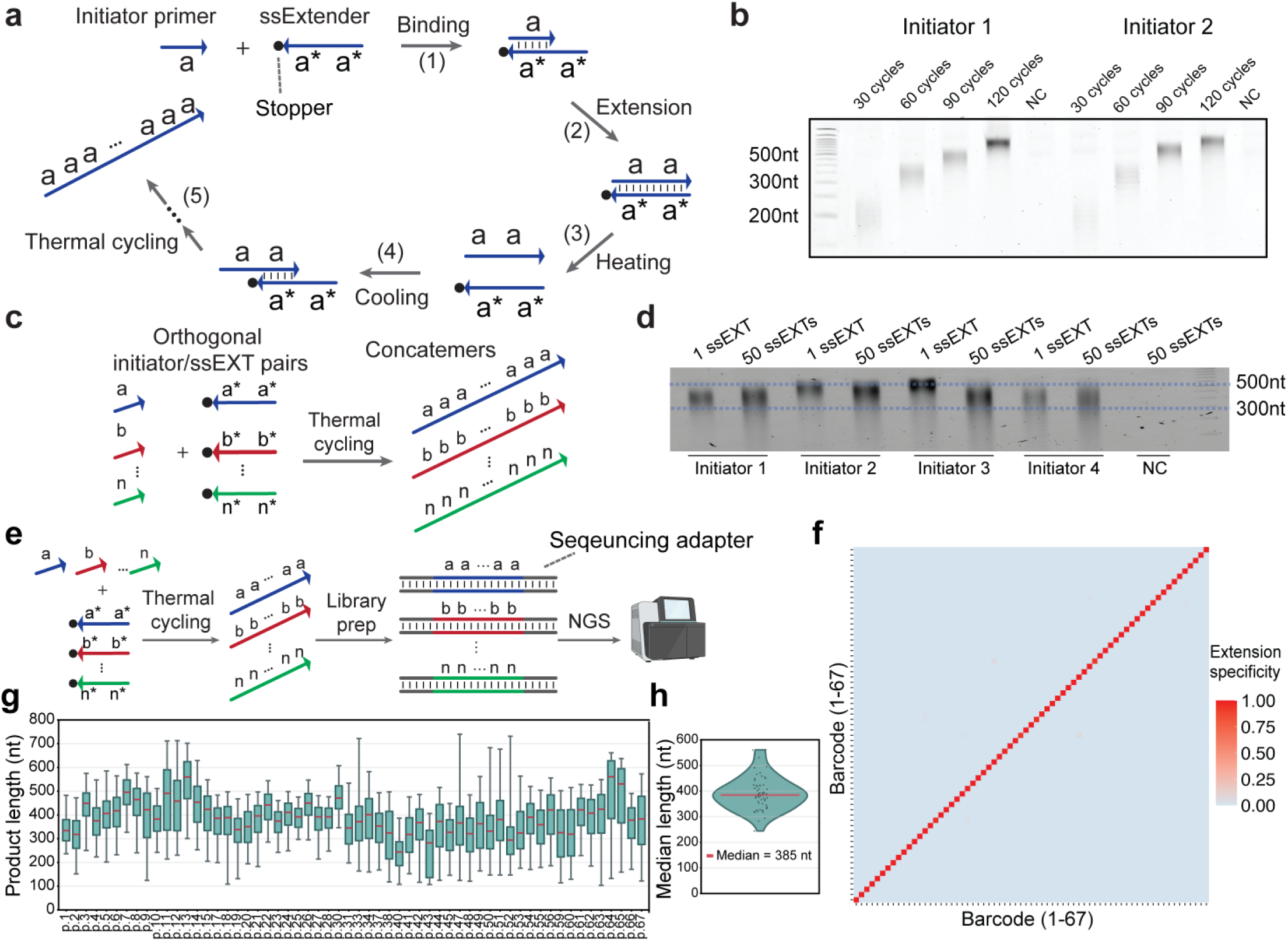
Characterization of the ACE chemistry. (**a**) Schematics of ACE amplification. Black dots indicate the 3’ stopper on ssEXTs, which prevents ssEXT extension and allows ssEXTs to act as reusable templates. A 3’ inverted dT was used for this purpose in this study. In step (1), ssEXTs bind to primer initiators at low temperature. In step (2), the primer initiators are extended by DNA polymerization at low temperature. In step (3), the elevated temperature dissociates ssEXTs from extended primer initiators. In step (4), extended primer initiators rebind to ssEXTs when the temperature cools down. In step (5), repeated thermal cycling results in the generation of long concatemers. (**b**) A denaturing PAGE gel showing the size distribution of amplified concatemers for different ACE cycles. Two randomly selected sequences were used for ACE reactions with different number of amplification cycles in test tubes. The size of the concatemers increased with the number of thermal cycles. (**c**) Schematics of multiplexed ACE amplification. (**d**) A denaturing PAGE gel showing the effect of multiplexed extension on single primer initiators. Four different initiators were ACE amplified for 80 cycles using either a single ssEXT or a mixture of 50 ssEXTs. (**e**) Schematics of orthogonality tests. (**f**) Sequencing results of a 67-plex ACE reaction. 120 cycles of ACE extension were performed in a test tube and sent for NGS. The crosstalk percentage was analyzed by calculating the pairwise probability of extension for all 67 sequences. p.16, p.29, p.32, p.35, p.36, p.39, p.46, p.57, and p.58 showed extension specificity of <99% (see **Table S1** for sequence of barcodes). (**g**) Fluorescence-weighted product-length distributions after 80 extension cycles for 58 initiator–extender pairs with >99% specificity. Boxes indicate the 25th–75th percentiles, red lines denote medians and whiskers extend from the 5th to the 95th percentiles. (**h**) Violin plot showing the distribution of fluorescence-weighted median product lengths across the same 58 pairs. Points denote median product length of individual pairs, and the red line indicates the overall median (385 nt).

**Figure 2.**
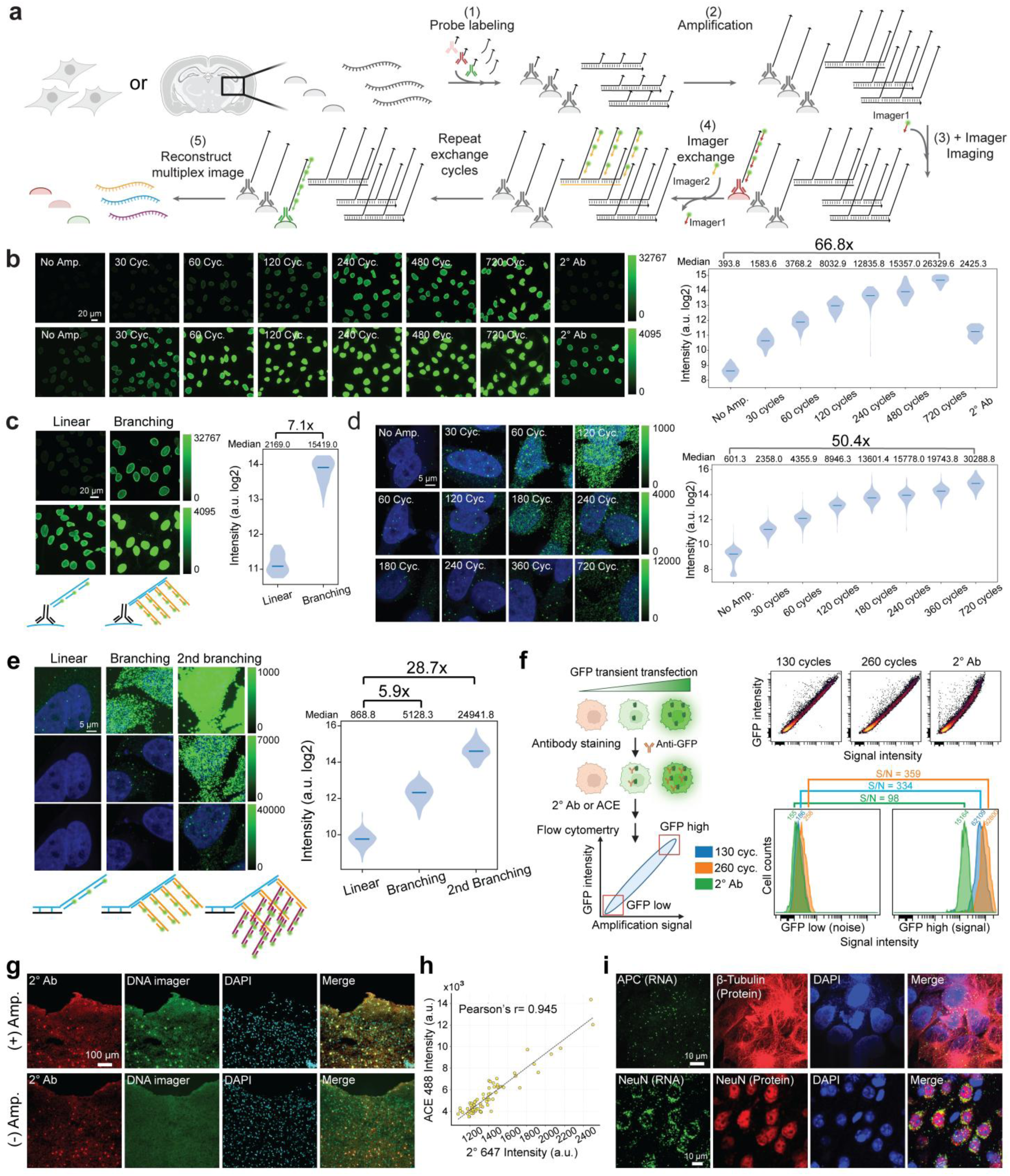
ACE allows high S/N signal amplification *in situ*. (**a**) Schematics of multiplexed ACE amplification for cell and tissue imaging. (**b**) Time-course analysis of ACE signal amplification for anti-LaminB in HeLa cells. For the “No Amp” sample, we performed a single extension step *in situ* to create a single imager binding site. Horizontal lines indicate medians. n = 123, 190, 158, 118, 196, 137, 106 and 56 nuclei for no amplification, 30, 60, 120, 240, 480 and 720 cycles, and secondary antibody-only control, respectively. (**c**) An additional 7.1× amplification can be achieved by concatemer branching. Horizontal lines indicate medians. n = 32 and 74 nuclei for linear and branching amplification, respectively. (**d**) Time-course analysis of ACE signal amplification for APC transcripts in HeLa cells. Horizontal lines indicate medians. n = 1,949, 1,817, 1,190, 1,613, 838, 1,635, 1,956 and 1,373 RNA puncta for no amplification, 1, 2, 4, 6, 8, 12 and 24 h, respectively. (**e**) Concatemer branching and iterative concatemer branching allow for further 5.9× (branching) and 28.7× (2nd branching) signal amplification for RNA targets. Horizontal lines indicate medians. n = 1,235, 1,499 and 1,340 RNA puncta for linear, branching and second-branching amplification, respectively. (**f**) GFP-transfected HEK293T cells were stained with anti-GFP primary antibody followed by ACE amplification or secondary antibody staining. ACE signal correlated strongly with GFP intensity after 130-cycle and 260-cycle amplification, with Pearson’s r = 0.9365, n = 13,593 events, and r = 0.9383, n = 13,690 events, respectively. Secondary antibody staining showed Pearson’s r = 0.8433, n = 13,053 events. ACE achieved a signal-to-noise ratio >300, compared with 97.8 for secondary antibody staining. (**g**) Signal amplification is needed to visualize SOX10 in thin mouse brain sections. (**h**) Scatter plot comparing raw mean nuclear intensities of SOX10-positive cells (N=71) between secondary antibody (x-axis) and ACE imager (y-axis) in the same individual cells. The dashed line represents the linear fit. Pearson’s r=0.945. (**i**) Co-imaging of RNA and protein in HeLa cells (RNA: APC, protein: β-tubulin) and mouse brain sections (RNA: Rbfox3, Protein: RBFOX3).

We validated 67 initiator/extender pairs and demonstrated that ACE can achieve 50-to over 400-fold signal amplification and higher signal-to-noise ratio than conventional immunofluorescence. Beyond protein imaging, ACE can also be applied to RNA targets, enabling the co-detection of RNA and protein within the same cells and tissue samples. Using ACE, we demonstrated 25-plex imaging of proteins and RNAs in Py2T cells at single-punctum resolution. We further achieved 63-plex protein and RNA mapping in a whole 20 µm mouse brain sagittal section, 22-plex volumetric protein imaging in a 200 µm thick mouse brain section (18.8 mm × 12.2 mm; > 2 million cells), and 16-plex volumetric protein imaging in a 1 mm thick coronal section (5.4 mm × 4.5 mm; > 3 million cells) with <8 hour target switch time. ACE enables depth-uniform, highly multiplexed signal amplification in thick tissue with accelerated target exchange cycles, and thus advances scalable multiplexed volumetric molecular imaging.

## RESULTS

### ACE enables multiplexed thermal cycle–controlled DNA extension *in vitro*

The ACE reaction relies on iterative, controlled DNA extension, paced by rapid temperature cycling. The kinetics of this reaction are predominantly governed by the programmed temperature cycles, which synchronize cycles of ssEXT binding, primer extension and ssEXT dissociation (**Fig. 1a**). Specifically, 3’-sealed, 19-nt, single-stranded extenders (ssEXTs), featuring dual repeats *a*a\** and the complementary initiator sequences *a*, are mixed with DNA polymerase and dNTPs in reaction buffer. The 3’ blocker prevents extension of the ssEXT itself, allowing ssEXTs to act as reusable templates during cyclic extension; a 3’ inverted dT was used for this purpose in this study. At a low temperature (22 or 40 °C were experimentally used in this work), ssEXTs bind to the designated initiators and enable their extension through DNA polymerization. A quick temperature rise (58 °C) triggers the dissociation of the ssEXTs, as the melting temperature (Tm) of ssEXTs was designed to remain below 56 °C in the given reaction buffer. This temperature cycling between a low temperature (22 to 40 °C) and 58 °C repeats, enabling the reannealing and elongation of initiators, resulting in the production of concatemers composed of repetitive sequence *a in situ*.

ACE enables cycle-dependent control of product extension. To demonstrate this, we randomly selected two DNA barcodes from our library and subjected them to varying temperature cycle numbers in parallel (**Fig. 1b**). Consistent extension patterns were observed at four distinct time points during the process.

ACE can generate concatemers in a multiplex fashion (**Fig. 1c**) if the initiator/ssEXT pairs are orthogonal to each other at reaction temperatures. To test this hypothesis, we computationally designed 67 orthogonal initiator/ssEXT pairs (**Table S1**) using seqwalk^40^ and NUPACK^41^ with the goal of minimizing crosstalk and maximizing extension efficiency. It is worth noting that the multiplexity is scalable with increasing the length of initiator/ssEXT pairs in computational simulation (**Fig. S1a;** please refer to **Table S1** for 92 available 9-nt sequences based on seqwalk).

To test whether applying multiple ssEXTs disrupts the ACE amplification process or alters extension uniformity, we examined the impact of applying multiple ssEXTs to individual initiator extensions. Four initiators were randomly selected, and ACE thermal cycling was conducted using either a single ssEXT or 50 ssEXTs. No major changes were detected in the size distribution of the extension on a denaturing PAGE gel (**Fig. 1d**).

To quantitatively assess the orthogonality of extension via ACE, we combined 67 initiators and extended them with 67 ssEXTs in a single PCR test tube (**Fig. 1e**). We then performed library prep for the extended products and sequenced the library with next-generation sequencing. The specificity of each initiator/ssEXT pair was evaluated. Our results revealed that 66 out of the 67 pairs exhibited an extension specificity >95% (**Fig. 1f**) and 58 pairs had a specificity >99%. Notably, among the pairs with specificity below 99%, each had a distinct off-target initiator/ssEXT pair, rather than exhibiting broad, nonspecific extension.

We next characterized the extension performance of ACE. Following 80 extension cycles, 52 of the 58 initiator–ssEXT pairs with greater than 99% specificity had median product lengths between 300 and 500 nt (**Fig. 1g** and **Fig. S1b**). Across the 58 pairs, the mean per-pair median product length was 391 nt, with a coefficient of variation of 16.4%, and the overall median was 385 nt (**Fig. 1h**).

These results suggest ACE allows for *in vitro* multiplexed linear amplification in a controllable manner.

### ACE allows for high signal-to-noise amplification *in situ* for protein and RNA imaging

ACE enables multiplexed signal amplification for *in situ* biomolecular imaging by labeling biomolecules with orthogonal initiators. To achieve this, we conjugated Ab and designed RNA probes with orthogonal short initiators. We then stained the sample with the initiator-barcoded Abs and/or RNA probes to label targets of interest. We performed ACE to amplify the signal and used fluorophore-labeled DNA imager exchange to switch targets rapidly (**Fig. 2a**). ACE applies sequential target-specific imaging, in which each target is identified by its specific imager during a predefined imaging round.

To evaluate whether DNA barcode conjugation preserves antibody staining behavior, we first validated the Ab-initiator conjugation by gel band-shift assays (**Fig. S2a**). We compared DNA-conjugated antibodies detected by DNA imager readout with the corresponding unconjugated antibodies detected by conventional fluorescent secondary antibodies. Across representative brain markers, including ALDH1L1, IBA1, NeuN and NPY, the conjugated and unconjugated antibodies produced closely matched anatomical and cellular staining patterns, supporting that the ACE DNA barcode conjugation and DNA-imager readout did not detectably alter the expected staining pattern of these antibodies (**Fig. S2b**). To further test whether conjugation introduced target-independent nonspecific binding, we performed a genetic loss-of-function validation using parental A549 cells and a matched EGFR-knockout A549 line. In wild-type cells, both unconjugated anti-EGFR detected by secondary antibody and DNA-conjugated anti-EGFR detected by ACE amplification produced membrane-localized EGFR signal. In EGFR-knockout cells, neither readout produced EGFR signal above background (**Fig. S2c**). Single-cell intensity quantification confirmed strong EGFR signal in wild-type cells and near-background signal in knockout cells for both ACE and conventional secondary-antibody detection (**Fig. S2d**).

To validate ACE *in situ*, we first confirmed that 22-58 °C thermal cycling does not alter the spatial distribution of protein and RNA (**Fig. S3a(i-ii)**). Subsequently, we assessed the specificity and preservation of morphology using ACE *in situ*. Using β-tubulin as a model target, we fixed HeLa cells with paraformaldehyde and stained them with initiator-barcoded anti-β-tubulin primary Abs. Distinct tubular structures were clearly visualized following ACE amplification (**Fig. S3a(i)),** and the observed patterns closely matched those obtained with conventional fluorescent secondary Ab staining. RNA puncta also showed high spatial overlap before and after ACE thermal cycling (**Fig. S3a(ii)**). We then further refined the ACE protocol *in situ* by optimizing the concentration of each component (e.g., polymerase concentration shown in **Fig. S3b**).

Next, we validated ACE for RNA imaging *in situ*. To test the specificity of the observed RNA signal, we labeled a housekeeping gene (APC) with two sets of ACE RNA probes. As expected, the two probe sets produced highly colocalized signals after 120 cycles of ACE amplification, with 94.5% puncta overlap, comparable to the overlap observed in the matched smFISH-equivalent sample. The ACE signal intensity was also markedly higher than noise from nonspecific probe binding (**Fig. S3c**; see **Table S2** for probe sequences).

We then characterized the ACE amplification *in situ*. We first evaluated ACE amplification for proteins using initiator-barcoded anti-Lamin primary Ab. In the case of linear amplification, ACE achieved a >20-fold signal amplification in comparison to cells with no amplification over 120 cycles of thermocycling (∼4 hours) and reached ∼67-fold amplification after 720 cycles (∼24 hours) (**Fig. 2b**). In contrast, secondary Ab, targeting primary Ab, exhibited a 6.2-fold amplification in the same batch of the sample, equivalent to 30-60 cycles (1-2 hours) of ACE extension.

The ACE signal can be further augmented using a branching amplification strategy (**Fig. 2c**). After the initial linear extension of concatemers, *ex situ* ACE-generated branching concatemers can be introduced to bind to the *in situ* generated concatemers, producing an increased number of imager binding sites on Abs (**Fig. 2c**). In HeLa cells, we observed an additional 7.1-fold amplification of the Lamin signal through a second ACE reaction on branching probes in ACE pre-amplified samples, culminating in a total amplification exceeding 400-fold. This amplification can be further enhanced with additional thermocycling rounds.

We then characterized ACE signal amplification for smFISH targeting the APC gene (**Fig. 2d**), where pronounced, punctate signals were observed at ∼50-fold amplification compared to smFISH without amplification over 720 cycles of ACE (∼24 hours). Branching offers 5.9-fold amplification of signal intensity, whereas second branching shows > 28-fold amplification, compared with linear ACE (**Fig. 2e**), making the total amplification > 1,000-fold.

A high S/N is critical for precise spatial profiling of targets, particularly those present in low abundance. Many existing amplification techniques depend on the incorporation of *ex situ* synthesized preamplifiers into these targets^21,22^. However, any non-specific binding of these preamplifiers can induce unintended signal amplification, thereby compromising the S/N. Additionally, it has been shown that conjugation of long DNA strands to the Ab, a strategy commonly used by previous methods, can also result in nonspecific signals^33^. In contrast, the ACE protocol does not introduce *ex situ* synthesized long amplifiers, and DNA barcodes are extended *in situ*. We also employ a short 11-nt barcode (a 9-nt ACE initiator and a 2-nt spacer) to reduce potential nonspecific Ab binding.

We first characterized the S/N of ACE in cell imaging. To evaluate non-specific extensions, we used FAM-labeled dCTP to fluorescently label concatemers during ACE DNA polymerization in HeLa cells with or without initiator-barcoded anti-β-tubulin Ab (**Fig. S4a**). We observed strong staining of tubulin with Ab addition while the no-Ab sample showed background comparable to the no-extension negative control.

To further evaluate S/N with DNA imager addition, HeLa cells stained with a series of increasingly diluted, initiator-barcoded anti-β-tubulin Ab, or no primary Ab, were ACE amplified and stained with either fluorescent DNA imagers or secondary Ab. Cells stained with secondary Abs exhibited markedly higher background than those imaged with ACE (**Fig. S4b**). We then transfected HeLa cells with GFP-β-tubulin fusion protein and stained the cells with diluted anti-GFP Ab (100 pg/mL) (**Fig. S4c**). We observed discrete punctate signals in the ACE-amplified condition that colocalized with tubules in the GFP+ cells. Quantitative analysis of the 100 pg/mL anti-GFP antibody experiment revealed that ACE amplification achieved a 14.9-fold increase in mean puncta intensity in transfected (GFP-high) cells over primary-omitted negative controls. In contrast, conventional secondary-antibody detection yielded only a 3.34-fold increase over background under identical low-input conditions, representing an approximately 4.4-fold improvement in signal-to-background separation for ACE over traditional secondary antibody detection (**Fig. S4d**). This indicates the high specificity of the ACE signal and is consistent with the capability of resolving the signal at single target level. This capability is helpful in profiling many targets that serve as pivotal regulators of cellular functions that are expressed at a low level (e.g., transcription factors).

To quantify the S/N of ACE compared with the conventional secondary Ab staining, we transfected HEK293T cells with GFP and quantified the amplified signal intensity across GFP-high and GFP-low cell populations via flow cytometry (**Fig. 2f**). ACE exhibited an S/N over 300 and strong correlation of raw signal intensity with Pearson’s r values of 0.94, in contrast to the sub-100 ratio observed with secondary Ab staining.

To test whether amplification is needed for tissue imaging of low-abundance targets, we imaged SOX10 on mouse brain sections with or without ACE amplification (**Fig. 2g**). While we can observe clear positive signals with ACE amplification and secondary Ab, the no-amplification condition suffers from a low S/N ratio, which makes positive signals difficult to distinguish from tissue background. To evaluate signal fidelity, we quantified the intensity correlation within the same SOX10-positive cells (**Fig. 2h**). The ACE imager and secondary antibody signals showed high correlation (Pearson’s r=0.945), indicating that ACE amplification preserves the spatial and quantitative distribution of the target protein at the single-cell level.

To assess run-to-run reproducibility of ACE signal measurements, we performed two independent ACE reactions on adjacent serial 12 µm mouse brain sections using the same three-channel panel: DAPI, NeuN, and OLIG2. The two runs produced matching anatomical staining patterns across the tissue section (**Fig. S5a**). We next compared cell-level intensity distributions for all DAPI-positive cells, NeuN-positive neurons, and OLIG2-positive oligodendrocyte-lineage cells. Median intensities were closely matched between the two independent runs, with Run 1/Run 2 ratios of 1.029 for DAPI, 1.008 for NeuN, and 0.991 for OLIG2 (**Fig. S5b**).

We next tested co-imaging of protein and RNA in one sample using ACE. We first stained HeLa cells with initiator-conjugated Ab, followed by RNA probe hybridization. ACE amplification preserved both protein and RNA signals (**Fig. 2i**). Similarly, we successfully co-imaged protein and RNA in mouse brain sections (**Fig. 2i**).

These results demonstrate the reduced noise and flexible signal amplification using DNA imagers with ACE, resulting in high S/N for imaging biomolecules in cells and tissues.

### ACE enables highly multiplexed multi-omics imaging in cells at single-punctum resolution

To demonstrate the utility of ACE for multiplexed imaging, we first evaluated whether introducing multiple orthogonal ssEXTs impacts the signal intensity of a specified barcode *in situ*. For this, we stained HeLa cells with initiator-conjugated anti-β-tubulin Ab and amplified the signal with ACE. Compared with an ACE reaction using a specific ssEXT, ACE with the addition of 50 orthogonal ssEXTs altered neither the signal’s intensity nor its morphology (**Fig. S6a**).

Following ACE amplification, we performed consecutive exchange cycles of imager addition, imaging, and imager removal. Both the morphology and intensity of the signal were maintained, even after >15 cycles of imager exchanges with cultured cells and mouse brain tissue sections (**Fig. S6b**).

To evaluate the specificity of the ACE chemistry *in situ*, we performed an imager leave-one-out control experiment for the set of 58 optimized sequences that have >99% specificity in the sequencing results (**Fig. 1f**). From this library, we selected nine representative sequences spanning the range of measured specificities (three close to 99%, three with intermediate specificity, and three close to 100% specificity). Each barcode sequence was conjugated separately to a primary antibody targeting Lamin A/C. HeLa cells were stained with the individual antibody conjugates, subjected to 58-plex ACE amplification for 120 cycles, and first imaged with the cognate imager to establish the on-target specific signal. Following signal removal by formamide washing, the same fields of view were re-imaged in the presence of a leave-one-imager-out pool containing all 57 non-cognate imagers (**Fig. S6c**). Quantitative single-cell analysis of the integrated background-subtracted signal intensities demonstrated that the cognate on-target signal was more than 35-fold higher than the nonspecific crosstalk signal across all nine sequences tested, with a mean signal-to-crosstalk ratio of 97.6-fold and a median of 79.7-fold (**Fig. S6d**). These results demonstrate that the filtered ACE barcode/imager library exhibits low crosstalk in the *in situ* imaging context.

To evaluate whether repeated imager exchange cycles cause cumulative signal loss, epitope degradation or cell detachment, we subjected ACE-amplified Lamin A/C staining in HeLa cells to 20 consecutive imager-exchange and imaging rounds over a 6-week period, mimicking the timescale of large-scale sequential imaging experiments (**Fig. S6e**). Lamin A/C showed preserved nuclear-envelope staining patterns across all 20 rounds, with no apparent loss of cellular morphology or cell attachment. Quantification of mean per-cell Lamin A/C intensity showed stable signal distributions across rounds, indicating that repeated imager hybridization, washing and imaging do not substantially degrade ACE-amplified targets under these conditions (**Fig. S6f**).

In high-plex imaging, sequence similarity between DNA imagers can lead to off-target hybridization to noncognate ACE amplifiers, particularly when the unintended target is highly abundant. To suppress this crosstalk, we introduced nonfluorescent “mock imagers” corresponding to all amplified targets, added at one-tenth the concentration of the fluorescent imager. These mock imagers preferentially occupy their fully complementary binding sites, reducing off-target binding of fluorescent imagers (**Fig. 3a**). In a representative comparison, mock imagers suppressed the crosstalk detected in the imager 2 channel while retaining the intended signal in the imager 1 channel, improving labeling specificity (**Fig. 3b**).

**Figure 3.**
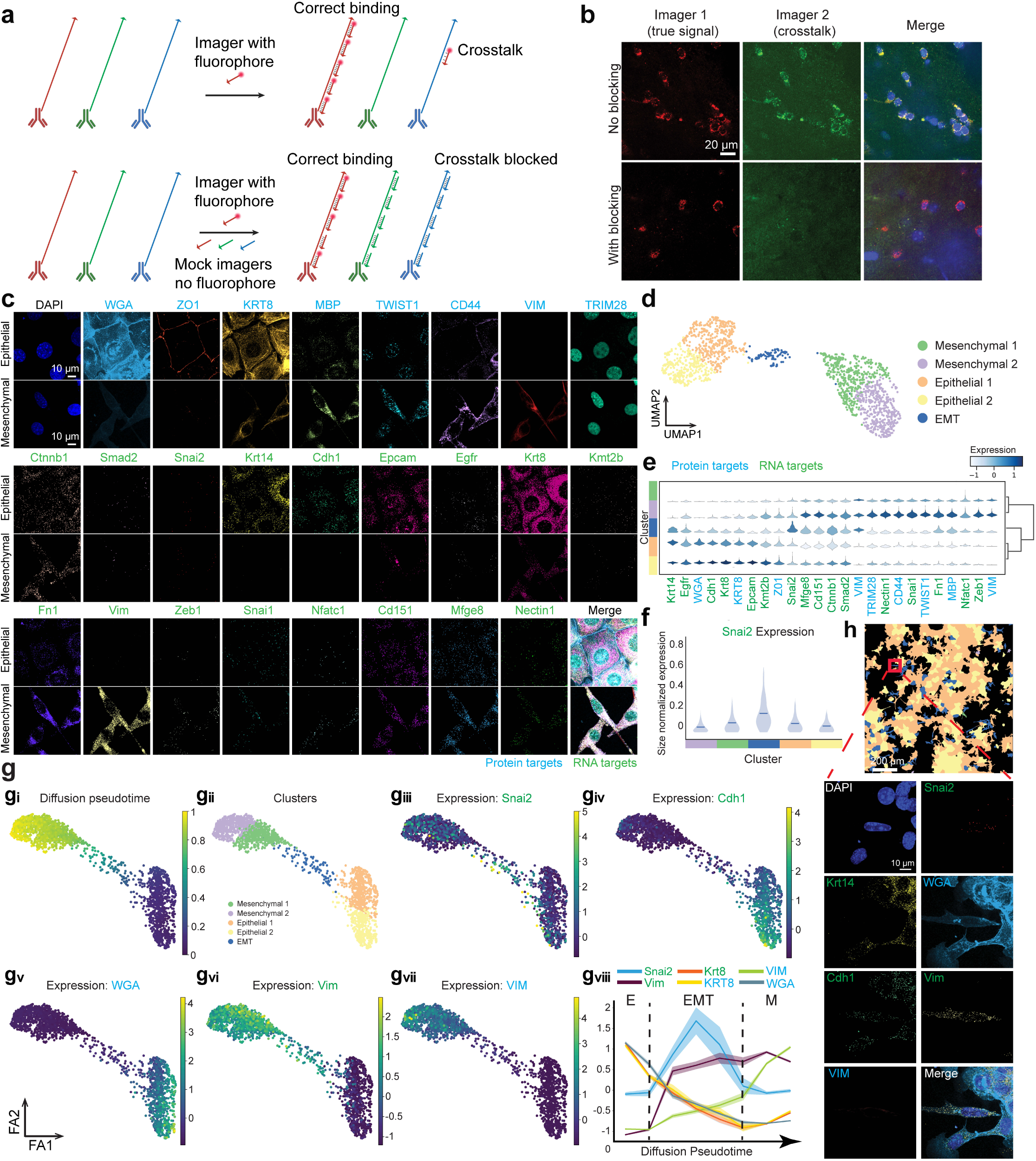
The mock imager strategy and multiplexed RNA and protein imaging of Py2T cells. (**a**) Schematic of mock imager–mediated suppression of sequence-dependent crosstalk in ACE imaging. Top, without mock imagers, fluorescent imagers bind to their intended targets but may also hybridize to partially complementary sequences on other ACE amplifiers, causing off-target labeling. Bottom, fluorescent imagers are added together with nonfluorescent mock imagers complementary to the amplifiers of all ACE-amplified targets, each at one-tenth the concentration of the fluorescent imager in this demonstration. By preferentially binding to their fully complementary sequences, mock imagers compete with off-target binding of fluorescent imagers, suppressing crosstalk while preserving labeling of the intended target. (**b**) The effect of mock imagers on imaging in tissue samples. Without mock imagers, the imager 2 channel showed nonspecific signals as it can bind to DNA for imager 1 binding. With mock imagers bound to DNA for imager 1, the imager 2 channel showed no detectable crosstalk signal. The mock imagers also did not substantially reduce the true signal intensity. (**c**) Spatial distribution of the 25 targets with or without TGFβ treatment in Py2T cells in the same field of view (See Figure S7a for the location of FOVs). Blue: Protein; Green: RNA. (**d**) UMAP plot of five molecularly defined Py2T clusters under different transition states (n = 1,699 cells). (**e**) The expression distribution and hierarchical clustering of the 25 targets of the 5 different clusters. (**f**) Snai2 RNA expression distribution among the 5 clusters. (**g**) Pseudotime analysis of the Py2T cells. **g(i)** The diffusion pseudotime of the cells. **g(ii)** Clusters of cells on the pseudotime map. **g(iii)** Expression of Snai2 RNA on the pseudotime. **g(iv)** Expression of Cdh1 RNA on the pseudotime. **g(v)** Signal intensity of WGA on the pseudotime. **g(vi)** Expression of Vim RNA on the pseudotime. **g(vii)** Expression of VIM protein on the pseudotime. **g(viii)** Selected markers’ expression changes at different pseudotime points. Thick solid lines indicate the mean normalized expression levels within each bin, and the surrounding shaded regions represent the standard error of the mean (SEM). (**h**) A representative image showing expression of selected targets in an EMT cell and its surrounding epithelial cells.

We next sought to demonstrate ACE’s multiplexity by studying a relatively well-defined biological process, the epithelial-mesenchymal transition (EMT)^42,43^. During EMT, an epithelial cell undergoes drastic RNA and protein expression changes to assume a mesenchymal cell phenotype. Transition to a mesenchymal phenotype involves changes in cytoarchitecture, elevated invasiveness, and increased migration capabilities. EMT plays a critical role in development, wound healing, cancer, etc. The Py2T cell line^44^ has served as a well-characterized model for studying the EMT process. Py2T cells possess an epithelial phenotype in standard cell culture conditions. Upon treatment with TGFβ, Py2T cells will gradually transition into mesenchymal-like cells, evident via morphological changes and elevated expression of mesenchymal RNA and protein markers.

To demonstrate the capabilities of ACE, we selected 17 RNA targets and 8 protein targets to study the process of EMT in Py2T cells, subjecting one condition to TGFβ treatment and the other to vehicle control (**Fig. S7a-b,** and **Table S3**). For each tile, a 15-μm z-stack comprising 31 planes was acquired. The total imaged regions measured 2 mm × 2 mm for cultured mesenchymal cells and 1.24 mm × 1.24 mm for cultured epithelial cells. After cell segmentation and filtering (**Fig. S7a,** and **Methods section**), we identified 826 cells in the control sample and 873 cells in the TGFβ-treated sample.

We were able to compare the expression profile of RNA targets, ranging from fewer than 10 puncta per cell to protein targets of high abundance between the two conditions (**Fig. 3c** and **S7c**). These results are corroborated by previous reports^44^, evident from similar enrichment of protein and RNA targets in a particular condition as well as distinctive morphologies of each. Notably, glycoprotein staining (WGA) was substantially reduced during EMT, indicating a global downregulation of glycosylation levels during the process.

Using the 25 RNA and protein targets, we clustered the cells (**Fig. 3d**) and identified 5 different clusters with distinct expression profiles (**Fig. 3e** and **S7d**). Of the 5 clusters, two were epithelial-like, two were mesenchymal-like, and we also identified an intermediate cluster, as shown in the Uniform Manifold Approximation and Projection (UMAP) plot (**Fig. 3d**), likely intermediates in the EMT process. Notably, ACE imaging suggested that whereas the cells treated with TGFβ all clustered as mesenchymal, the control sample had a small percentage of cells either undergoing EMT or having completed EMT. Notably, while Snai1 RNA showed higher expression in the mesenchymal clusters, we observed that its homolog Snai2 RNA was highly expressed only in the intermediary EMT cluster (**Fig. 3f** and **Fig. S8a**), suggesting the SNAIL2 transcription factor potentially plays an important role in the transition. Consistent with our findings, a recent computational analysis study indicates that SNAIL2 inhibits cells from undergoing a complete EMT and thus stabilizes them in a hybrid E/M phenotype^45^.

We then performed pseudotime analysis^46,47^ on the cells (**Fig. S8b**). Consistent with the clustering observation, we observed that Snai2 was highly expressed in the transition state between epithelial and mesenchymal phenotypes (**Fig. 3g**). Notably, while an epithelial marker Krt8 showed a similar expression pattern of protein and RNA across pseudotime, the Vim RNA, whose protein product vimentin (VIM) is a mesenchymal marker, showed earlier expression during EMT than its protein product (**Fig. 3g**). To validate the analysis, we selected a field of view containing a representative intermediary EMT cell and its surrounding epithelial cells (**Fig. 3h**). Indeed, in ACE imaging, the intermediary cell expressed a high level of Snai2 RNA, epithelial markers (Cdh1, WGA), and Vim RNA, but only a low level of VIM protein. This pattern is consistent with stepwise molecular changes during EMT reported in prior time-course and transition-state studies^48,49^, although differences in RNA-probe and antibody detection sensitivity may also contribute to the apparent offset.

These findings illustrate ACE’s ability for highly multiplexed mapping of RNA and protein targets across a high dynamic range, spanning single-molecule RNA puncta to highly abundant protein targets.

### ACE enables molecular mapping in brain tissue sections

We next applied ACE to tissue imaging. Understanding the function and biological processes of complex organs hinges on the spatial mapping of a plethora of molecular targets across various cell types. For instance, the brain consists of numerous distinct and interconnected functional regions inhabited by various cell types, including neurons, glial cells, and immune cells. High-throughput capabilities are important for efficiently imaging brain samples and comprehending their cellular and molecular landscape. The ACE technique, with its high S/N signal amplification and high multiplexity, shows potential in attaining the requisite throughput and clarity for effectively imaging tissues marked by high complexity and significant autofluorescence.

To demonstrate the throughput and multiplexing potential of ACE in thin mouse brain sections, we carried out a pilot 32-plex (29 protein and 3 RNA targets) imaging study on a 10 µm mouse coronal section, as shown in **Fig. S9a-c**. The results showed consistent staining with reference single-plex imaging from previous atlas studies ^50-53^.

To further demonstrate the capabilities of ACE, we expanded our target list to include 30 RNA and 33 protein markers, detailed in **Table S4**. These targets were subsequently imaged on a mouse sagittal section (15.3 mm × 7.1 mm × 20 µm), effectively utilizing the comprehensive range of capabilities offered by ACE technology. Initially, we imaged the RNA targets after the ACE amplification, followed by the complete removal of the RNA signal through RNase treatment. Subsequently, the same sample was labeled using DNA-barcoded Ab, subjected to another round of ACE amplification, and imaged with DNA imagers (**Fig. 4a**). We confirmed the thoroughness of RNA removal after RNase digestion (**Fig. S9d)**. To validate the observed ACE signals, each protein staining pattern was initially screened with DNA imagers and fluorescent secondary Abs to ensure colocalization with the primary Abs. RNA signals were compared to ISH results from the Allen Brain Institute ^50^.

**Figure 4.**
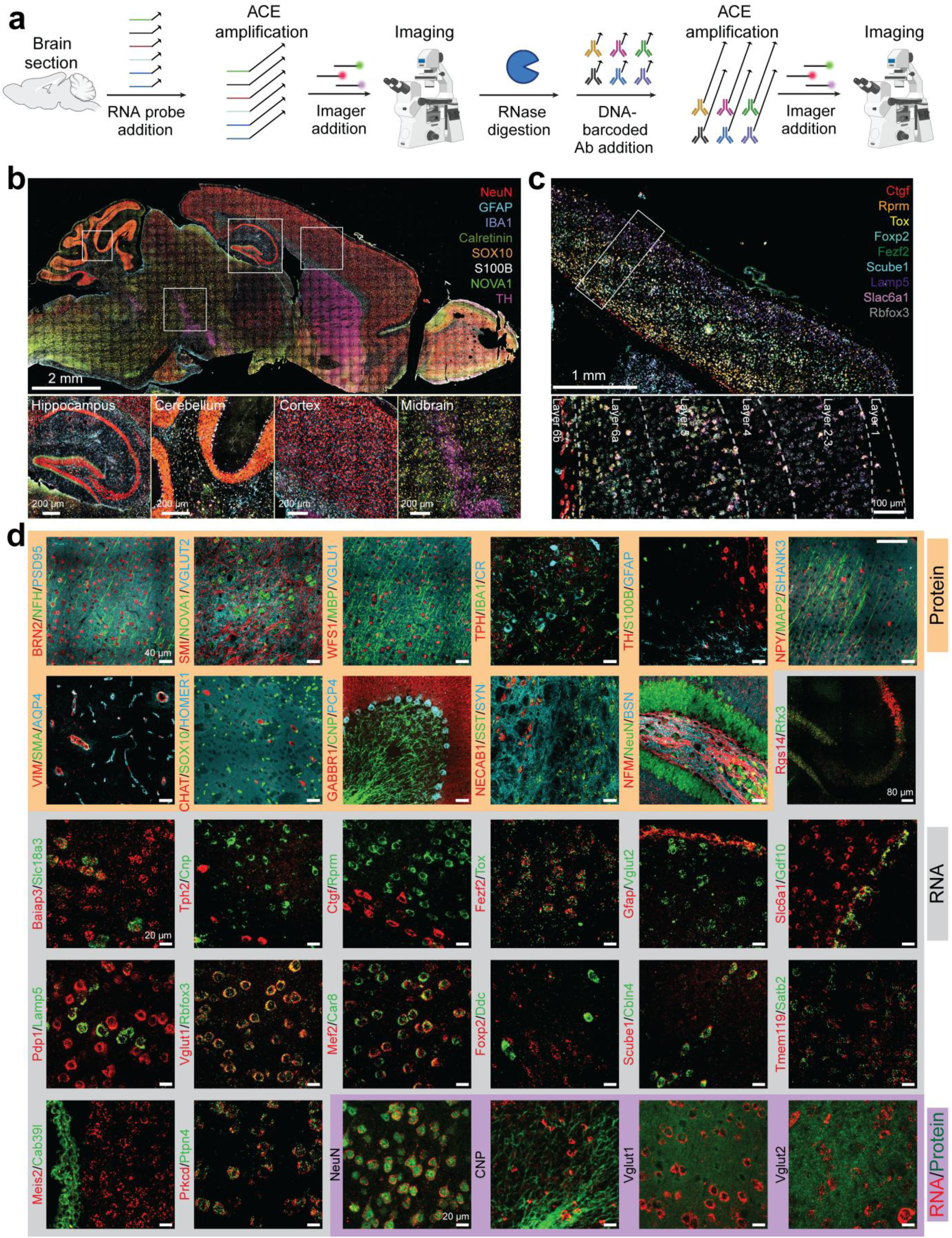
ACE multiplexed imaging of a mouse sagittal section. (**a**) Schematic for thin mouse brain imaging studies using ACE. (**b**) Top: a zoom-out image of selected protein targets on the entire mouse brain sagittal section. Bottom: zoom-in views of different regions. (**c**) Top: a zoom-out view of selected cortex layer RNA markers. Bottom: a zoom-in view of the cortex layers. (**d**) Representative zoom-in view of the 33 protein and 30 RNA targets at the cellular level. Protein and RNA co-staining of selected markers is also shown.

As demonstrated in **Fig. 4b**, ACE allows us to accurately map the spatial distribution of various cell types across multiple brain regions with protein targets. Further, ACE precisely delineated different cortical layers with a multiplexed RNA dataset using a 40× objective, highlighting the detailed and accurate mapping capabilities of the method (**Fig. 4c** and **Fig. S10**). Additionally, we showcased the ability of ACE to visualize multiple modalities at the same time, revealing information about structure and effector molecules via protein imaging, as well as deeper probing of cell states via RNA imaging (**Fig. 4d**). This dual functionality emphasizes the robustness of ACE in providing comprehensive insights into cellular composition and structure. This approach enables straightforward visualization of distinct expression profiles for RNA and protein originating from the same gene.

We next sought to use the ACE imaging to distinguish different cell types. After cell segmentation and marker quantification, we identified 113,547 cells (**Fig. 5a**). We then quantified the expression level of RNA and protein targets for cells identified with the tissue mask (**Fig. S11-S12**) and on UMAPs (**Fig. S18a, Fig. S14-S15,** and **Table S5**). Next, we performed clustering of cells using either 30 RNA targets, 14 protein targets, or 30 RNA + 14 protein targets. We identified 27, 14 and 31 clusters respectively (**Fig. 5a-b, Fig. S13a-d, Fig. S16-S17,** and **Fig. S18b**). We were able to annotate the clusters based on gene expression and spatial distribution (**Fig. 5b(iii)**). We identified major cell types (different neurons, astrocytes, microglia, oligodendrocytes, etc.) and were able to accurately map them onto a regional organization previously described ^50-53^. The different clusters show distinct expression profiles (**Fig. 5c** and **Fig. S13e-f**). These results highlight the accuracy and throughput of ACE.

**Figure 5.**
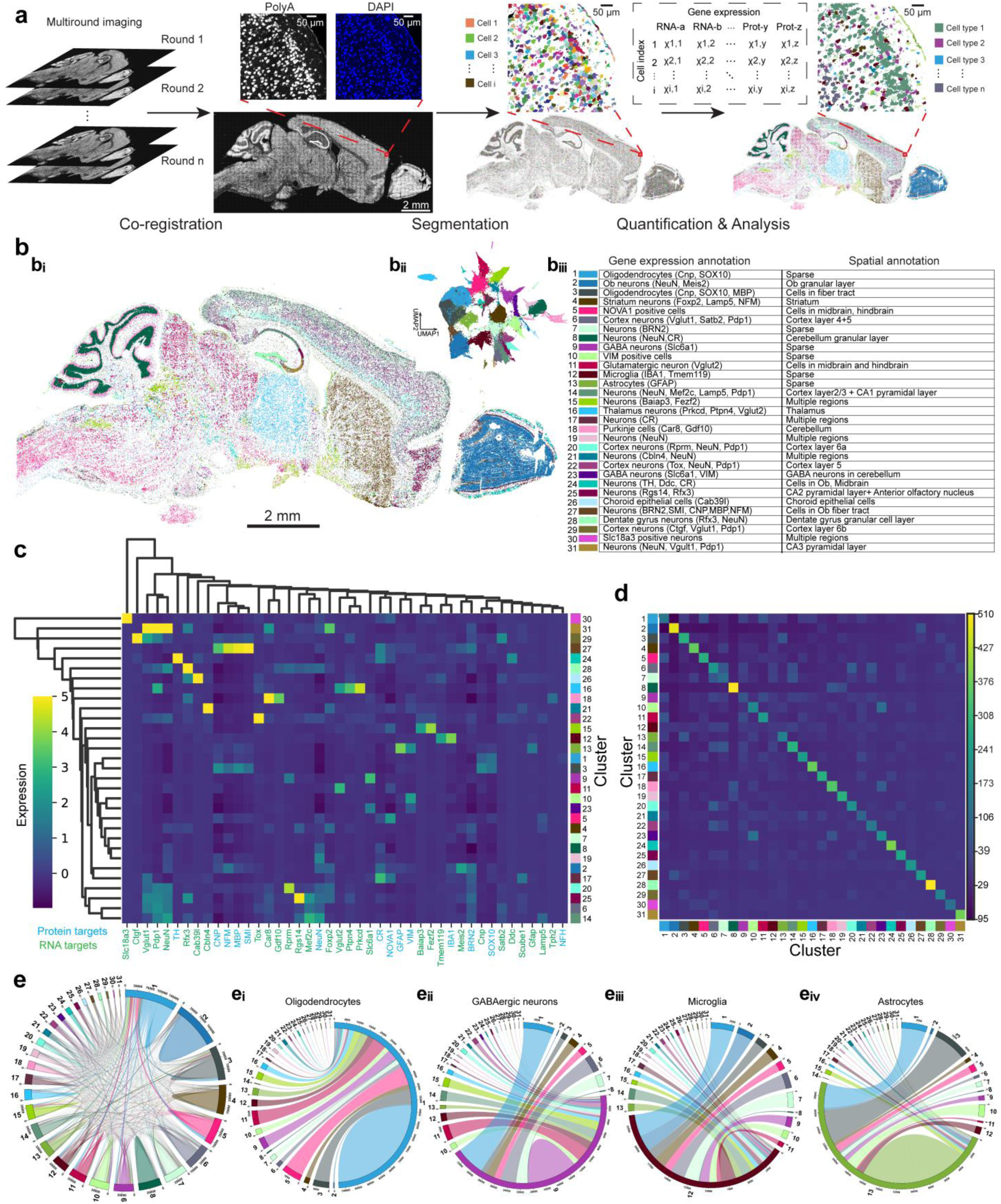
Data analysis of the mouse brain ACE imaging. (**a**) Schematics and representative cells for the segmentation and analysis pipeline. (**b**) **b(i)** Molecular cell-type maps and annotation of mouse sagittal section colored by clusters generated using RNA+protein targets. n = 113,547 cells. Each spot represents one cell. **b(ii)** UMAP plot of 31 clusters with the same color code as in Fig. 5b(i). Clusters were generated using selected RNA+ protein targets. **b(iii)** Annotation of the clusters using gene expression and the spatial distribution. (**c**) The expression distribution and hierarchical clustering of the selected RNA and protein targets for the 31 different clusters with the same color code as in Fig. 5b(i). n = 113,547 cells. (**d**) A heatmap showing neighborhood enrichment analysis of the 31 clusters with the same color code as in Fig. 5b(i). High values indicate higher physical proximity of cells in clusters. n = 113,547 cells. (**e**) Chord diagrams showing the abundance and proximity among clusters. Each pie corresponds to a specific cluster, with its size proportional to the number of cells passing proximity threshold of the cluster relates to. n = 113,547 cells. **e(i-iv)** Chord diagrams highlight the proximity networks of oligodendrocytes, GABAergic neurons, microglia, and astrocytes, respectively. The color code is the same as that in Fig. 5b(i).

We noticed that in addition to producing more clusters with protein and RNA, the degree of separation between the clusters was markedly improved, as evidenced by a comparison of the silhouette scores for each (**Fig. S13g**). Indeed, when looking at individual clusters, the protein + RNA clusters have better separation of selected cell types (**Fig. S13h**).

We also performed neighborhood analysis^54^ with the 31 clusters to assess spatial proximity among cell types. The resulting proximity patterns were broadly consistent with known brain organization (**Fig. 5d-e**). Oligodendrocytes, GABAergic neurons, astrocytes, and microglia exhibit relatively low intra-cluster proximity, consistent with their dispersed spatial distributions. Notably, we observed enriched proximity between oligodendrocytes, NOVA1+ neurons, and Vglut2+ neurons in the midbrain and hindbrain. GABAergic neurons also showed higher proximity with neurons in cortex layers and Vglut2+ neurons. Microglia were more frequently proximal to cortex neurons. Astrocytes, on the other hand, showed enriched proximity with cells in fiber tracts and VIM+ cells.

We next applied ACE imaging to expansion microscopy (ExM)^55^, which physically enlarges samples to improve effective spatial resolution. We observed well-retained signal integrity of classic cell-type markers (NeuN, GFAP, and SMI-312) and resolved the pre-synapses and post-synapses with synaptic markers in the same sample after a 3× expansion. (**Fig. S9e**).

These results demonstrate that ACE enables high-throughput, high-multiplex, and high S/N protein and RNA imaging in the tested cell and tissue samples, including cleared brain sections.

### ACE enables highly multiplexed volumetric protein imaging in 200 µm-thick brain tissues

Because the ACE protocol employs small reagents for multiplex amplification, we next tested whether ACE allows for multiplexed imaging at the whole tissue scale. To validate the approach, we first compared *in situ* ACE amplification with *ex situ* synthesized SABER concatemers in 1 mm mouse brain sections stained with NeuN. The ACE-amplified samples displayed uniform signal distribution across the 1 mm section. In contrast, the SABER concatemer-amplified samples exhibited signal depletion at depths exceeding 200 µm (**Fig. S19a**). This observation underscores the challenges of diffusing long concatemers deep into samples.

Subsequently, we demonstrated 22-plex ACE protein imaging across an entire 200 µm thick sagittal mouse brain section (18.8 mm × 12.2 mm). We employed a one-pot staining approach utilizing DNA-barcoded Abs targeting diverse cellular components, including cell-type-specific markers, axons, and blood vessels. Following ACE amplification, the tissue was embedded in hydrogel and imaged iteratively using a light-sheet microscope equipped with a 3.6× objective (**Fig. S19b**). Multi-round images were co-registered using POPO-1 nuclear staining. Specifically, the cells were detected through a blob detection algorithm from Bigstream ^36,56^ followed by point-cloud alignment using phathom ^57^. We used Imaris software (Andor Technology, Belfast, Northern Ireland) to further analyze the aligned images (**Fig. S19c-d**). This pipeline identified ∼2.5 million cells, enabling spatial profiling of marker distributions across the brain section (**Fig. 6a, Fig. S20– S21,** and **Video S1**).

**Figure 6.**
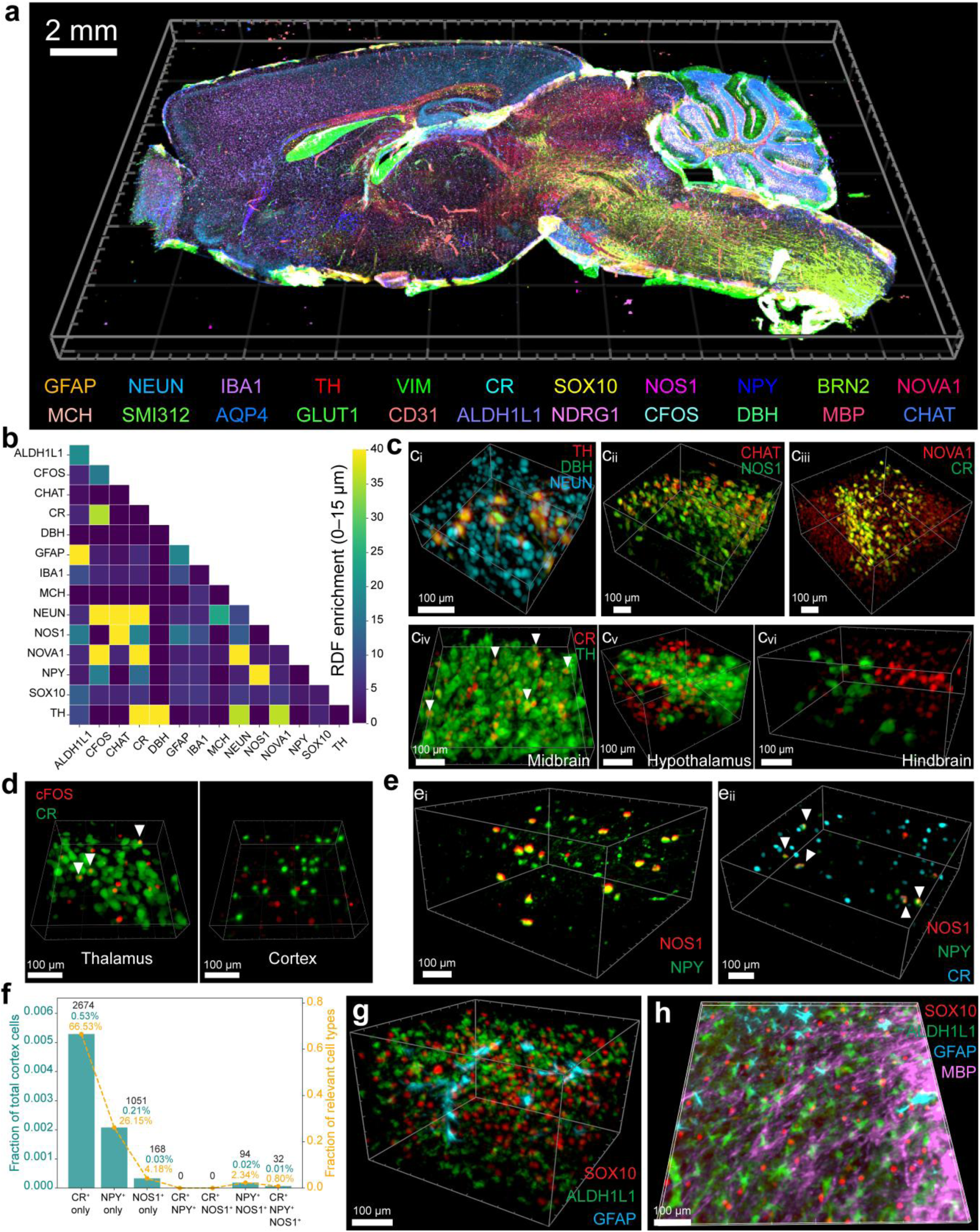
ACE multiplexed imaging of a 200 µm thick mouse sagittal section. (**a**) Multiplexed imaging of 22 markers in a 200 µm thick mouse sagittal brain section after co-registration. n = 2,542,660 cells. (**b**) Neighborhood analysis of spatial marker relationships across the tissue. n = 2,542,660 cells. (**c**) Imaging validation of selected spatial proximity relationships. **c(i)** Zoom-in view showing the spatial distribution of NeuN, DBH, and TH in the hindbrain. **c(ii)** Zoom-in view showing CHAT and NOS1 distribution in the hindbrain. **c(iii)** Spatial distribution of CR and NOVA1 in the hypothalamus. **c(iv-vi)** Spatial co-localization of CR and TH in the midbrain, hypothalamus, and hindbrain, respectively; arrows highlight double-positive cells. (**d**) Zoom-in views of CR and cFOS co-expression in the thalamus and cortex; arrows indicate cells positive for both markers. (**e**) Zoom-in view of the spatial distribution of **e(i)** NPY and NOS1, as well as **e(ii)** triple-marker expression of NPY, NOS1, and CR in the cortex. (**f**) Bar and line plot showing the distribution of CR⁺, NPY⁺, and NOS1⁺ interneuron subtypes in the mouse cortex. Bars (left y-axis, teal) represent the fraction of each subtype relative to the total number of cortical cells (n=505,510 cells), while the overlaid line plot (right y-axis, orange) shows the proportion of each group among the relevant marker-positive cells. Black numbers are the total population detected for each cell type. (**g**) Zoom-in view showing the spatial distribution of SOX10, ALDH1L1, and GFAP in striatum. (**h**) 3D rendering of SOX10, ALDH1L1, GFAP, and MBP in striatum. ALDH1L1⁺ astrocytes are spatially proximal to oligodendrocytes in the region.

Using 14 cell type markers, we extracted 3D coordinates for each labeled cell and performed a cell neighborhood analysis (**Fig. 6b**). As expected, NeuN⁺ cells exhibited strong spatial association with other neuronal markers such as cFOS, CHAT, CR, and TH. Subregional annotation and enrichment analysis further revealed region-specific associations (**Fig. S22a**). For example, DBH+ cells show high enrichment with TH+ cells in the hindbrain region but not with pan-neuronal marker NeuN+ cells (**Fig. S22d**). Consistently, DBH staining marked a TH⁺ subpopulation with low NeuN expression (**Fig. 6c(i)**). Similarly, CHAT⁺ cells in the hindbrain partially overlapped with NOS1⁺ cells (**Fig. 6c(ii)** and **Fig. S22d**). In most regions, NOVA1 co-localized with CR⁺ cells (**Fig. 6b**, **Fig. 6c(iii),** and **Fig. S22b-k**); however, CR⁺ cells were enriched near TH⁺ cells exclusively in the midbrain, not in the hindbrain or hypothalamus, despite expression in all three regions (**Fig. 6b**, **Fig. 6c(iv-vi), Fig. S22c,d,** and **Fig. S22g)**. cFOS⁺ cells showed specific enrichment with CR⁺ cells in the thalamus, but not in other regions (**Fig. 6d** and **Fig. S22b-k**). In the cortex, we quantified CR⁺, NPY⁺, and NOS1⁺ interneuron subtypes. CR⁺ cells represented the majority of relevant populations and were rarely co-expressed with either NPY or NOS1, except in the rare CR⁺NPY⁺NOS1⁺ subset (∼0.01% of total cells in cortex). In contrast, partial co-expression was observed between NPY⁺ and NOS1⁺ cells, indicating limited overlap among these interneuron subtypes (**Fig. 6e, f**). These results suggest substantial neuronal heterogeneity in different regions of the 200 µm mouse brain.

We also observed enrichment for two astrocyte markers, GFAP and ALDH1L1 (**Fig. 6b**). Notably, in striatum, ACE imaging showed that ALDH1L1⁺ cells displayed high proximity to SOX10⁺ oligodendrocytes whereas GFAP⁺ astrocytes did not (**Fig. 6g** and **Fig. S22j**). Notably, we also observed that ALDH1L1⁺ astrocytes were often found in close proximity to SOX10⁺ oligodendrocyte cell bodies and myelin sheaths (labeled by MBP), whereas GFAP⁺ astrocytes exhibited only limited spatial proximity (**Fig. 6h**).

### ACE enables highly multiplexed volumetric protein imaging in mm-thick brain tissues

We further extended ACE to enable high-plex imaging in millimeter-thick tissues (**Fig. 7a**). At this scale, maintaining physical integrity during iterative buffer exchanges is critical. Hydrogel-based protocols requiring protease digestion within soft gel matrices are less favorable due to their tendency to compromise tissue structural integrity during reagent exchange. Instead, advanced tissue preservation techniques such as iDISCO, SHIELD, and others^1-4^ have been developed to enhance molecular retention while maintaining structural integrity.

**Figure 7.**
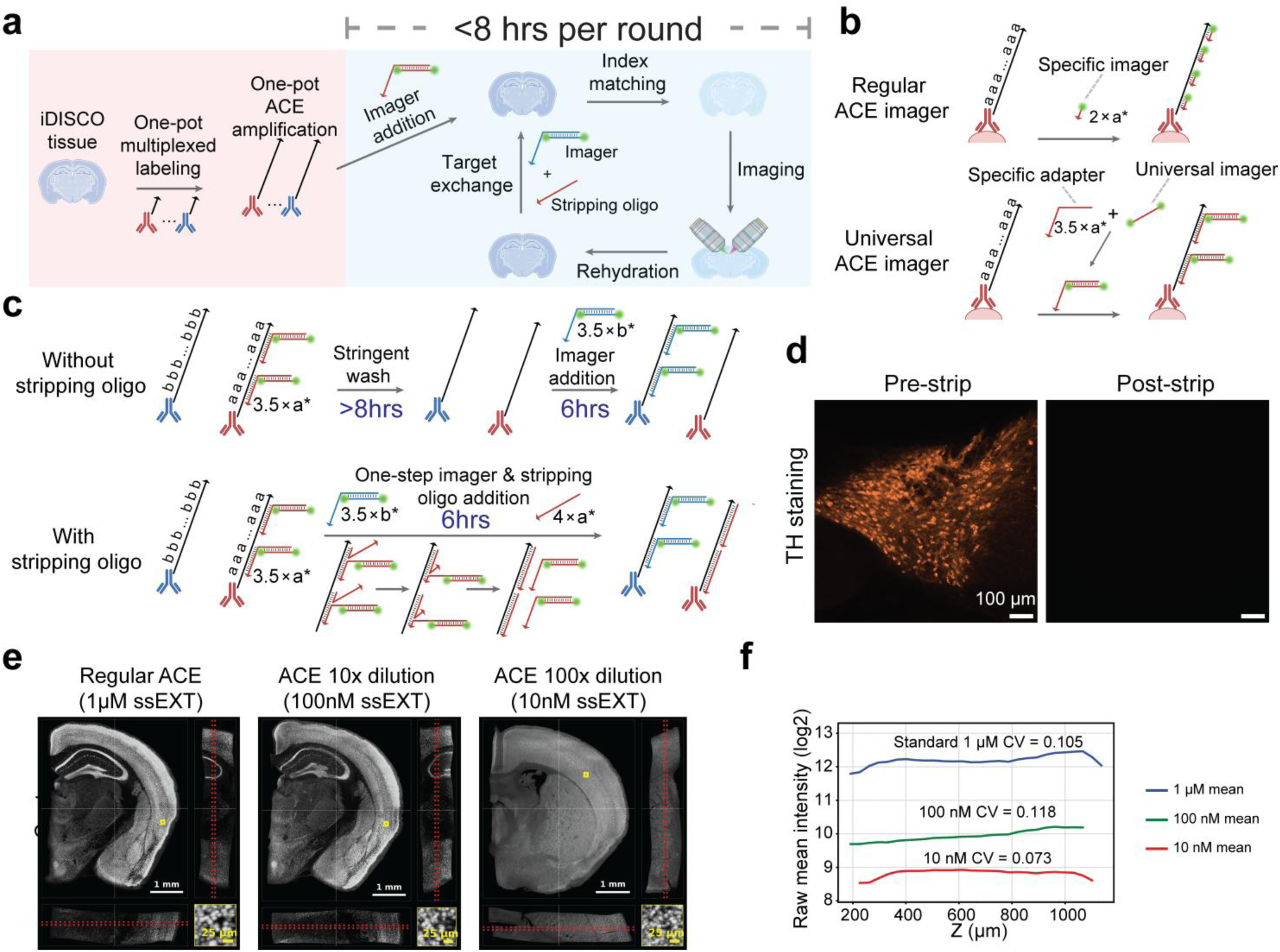
ACE reaction characterization in millimeter-thick tissue sections. (**a**) Schematic of whole-hemisphere coronal imaging of a 1 mm-thick cleared mouse brain section using the iDISCO protocol combined with ACE. (**b**) Schematic comparison of the regular ACE imager design and the universal ACE imager design. (**c**) Schematic comparison of the regular imager exchange protocol and the stripping-oligo–based exchange protocol. (**d**) Signal of TH in 1-mm iDISCO+ tissue before and after applying stripping oligos. (**e**) ACE NeuN imaging performed with preincubation using ssEXT at 1 μM (undiluted), 100 nM (10× dilution), or 10 nM (100× dilution). Tissue center orthogonal views and magnified insets are shown for each condition. Red dashed lines in the x–y and y–z views indicate the regions used to generate the corresponding maximum-intensity projections. The ssEXT dilution samples showed high depth uniformity over Z. Surface staining can be found in Fig. S30c. Display contrast was adjusted independently for each method or condition to visualize staining patterns. (**f**) Raw mean intensity profiles across z-depth for the same ssEXT dilution series, plotted on a log2 scale. Dilution of ssEXT reduced raw signal intensity over Z. However, depth profiles remained uniform with depth CVs of 0.105, 0.118 and 0.073 for 1 µM (regular ACE), 100 nM (10× dilution), and 10 nM ssEXT (100× dilution), respectively.

These methods are often coupled with index-matching solutions designed to minimize light scattering by aligning the tissue and protein refractive indices. Organic solvents such as ethyl cinnamate (ECi) are frequently used for this purpose. However, the index-matching workflow involves low-salt and organic-solvent conditions that can destabilize DNA duplexes. Stable DNA-imager binding is therefore required to maintain robust signal retention and uniform amplified readout throughout thick tissue.

To address this challenge, we adopted a universal imager design inspired by Stoichiometry-Invariant Multiplexable DNA Libraries (unpublished, Yin lab), incorporating a modular structure with a longer adapter sequence and a high-GC imager binding site (**Fig. 7b** and **Table S6**). The adapters hybridize to 3.5 repeats on the ACE concatemers, while the imager-binding site allows stable interaction with fluorophore-labeled DNA imagers, thereby improving hybridization stability in index-matching solvents. We validated dual-dye conjugation and purified the labeled oligos by HPLC (**Fig. S23a**).

We then compared regular ACE imagers with universal imagers in 1-mm iDISCO-cleared mouse brain tissue. Regular ACE imagers performed well in 1× PBS but were not robust after ECi refractive-index matching (**Fig. S23b**). For SOX10, the regular ACE imager showed increased nuclear background after ECi processing, whereas the universal imager maintained cleaner SOX10 staining. For GFAP, the regular ACE imager lost the expected astrocytic morphology after ECi processing, including fine protruding endfeet, whereas the universal imager preserved the expected GFAP-positive astrocyte processes.

We also found that passive probe wash-off was slow (>8 h, data not shown). To accelerate exchange, we designed stripping oligos that hybridize to four repeats of the ACE barcode sequence (**Fig. 7c** and **Table S6**). For each round, we added stripping oligos specific to the previous round’s targets together with the universal imagers for the current round. The stripping oligos efficiently depleted signal from the previous round (**Fig. 7d**), and when co-applied with universal imagers, we did not observe measurable reduction in signal intensity or alteration of staining patterns for the intended targets (**Fig. S23c**).

To assess the quantitative accuracy and spatial preservation of target protein intensities following signal amplification steps, we evaluated the voxel-level signal correlation between ACE and secondary Ab using anti-NeuN Abs on the same 1mm tissue sample. After alignment, ACE signal and secondary Ab signal showed spatial overlap of fine cellular morphology architectures with few apparent structural distortions (**Fig. S24a**). To extend this validation across the entire tissue volume, we performed a localized voxel-level intensity correlation analysis across five sequential depth zones (Z = 200 to 600 slices). Depth-dependent scatter plots revealed stable and high signal fidelity throughout the tissue, with Pearson correlation coefficients exceeding 0.87 (**Fig. S24b**). We next compared NeuN-positive cell counts obtained by ACE and conventional secondary-antibody detection across the full z-depth of the same 1-mm tissue sample. The two methods produced closely matched depth-dependent cell-count profiles, with a Pearson correlation coefficient over 0.99 (**Fig. S24c**). Together with the voxel-level correlations, these results indicate that ACE preserves both relative protein-intensity patterns and cell-positive classification across tissue depth.

To assess whether synchronized *in situ* amplification allows for depth-uniform protein imaging in thick tissue, we compared ACE with HCR and RCA, which amplify targets *in situ* without synchronized reaction initiation or progression, and immuno-SABER, which uses pre-synthesized concatemer amplifiers generated *ex situ* (**Fig. S25-S29**). We tested markers spanning different subcellular compartments and expression levels, including NeuN, an abundant nuclear neuronal marker (**Fig. S25**); CD31, a vascular membrane marker (**Fig. S26**); GFAP, a cytoplasmic astrocyte marker (**Fig. S27**); and OLIG2, a relatively lower-expression nuclear oligodendrocyte-lineage marker (**Fig. S28**). ACE produced the expected marker-specific staining patterns with more uniform signal across depth for all four targets whereas all other amplification methods tested exhibited pronounced surface-bright, center-dim profiles. HCR produced recognizable target-specific staining at the tissue surface, but signal intensity was more than 20-fold lower in the tissue center. Fluorophore-labeled tertiary-antibody staining of the HCR samples showed relatively uniform antibody labeling across depth, suggesting that the observed depth bias arose primarily during HCR amplification rather than from antibody penetration (**Fig. S25-S28**). Immuno-SABER produced recognizable NeuN signal, but the center signal was more than fourfold dimmer than the surface signal; for CD31, GFAP, and OLIG2, immuno-SABER showed substantial nonspecific nuclear background. RCA-based protein detection was tested for GFAP and NeuN, but also produced nonspecific nuclear background. Notably, even where high nonspecific background was observed with immuno-SABER and RCA, the overall signal remained strongly surface-bright and center-dim (**Fig. S29**).

To further test the mechanism underlying depth-uniform ACE amplification, we benchmarked ACE with secondary Ab staining and performed ssEXT titration and pre-incubation controls in 1-mm mouse brain tissue stained with anti-NeuN Abs. The ACE-amplified sample showed signal uniformity over Z comparable to that of secondary Ab staining (**Fig. 7e** and **Fig. S30a, c, d, f**). When ACE thermal cycling was initiated without pre-incubating the reaction mixture in the tissue, the amplified signal became surface-bright and center-dim (**Fig. S30b** and **Fig. S30e**). This result indicates that reagent equilibration before synchronized initiation is required for uniform amplification. We next reduced the ssEXT concentration from 1 µM to 100 nM and 10 nM (**Fig. 7e** and **Fig. S30c**). As expected, ssEXT dilution reduced the overall ACE signal intensity (**Fig. 7f**). However, the normalized depth profiles remained uniform and comparable to secondary Ab staining (**Fig. 7f** and **Fig. S30f**). These results show that ACE signal amplitude depends on ssEXT concentration, while depth uniformity is maintained when reagent equilibration is followed by synchronized cyclic amplification.

To demonstrate multiplexed volumetric imaging with ACE, we performed 16-plex protein imaging in 1-mm-thick coronal sections (5.4 mm × 4.5 mm) from iDISCO+-processed mouse brain hemispheres. Abs labeled with DNA barcodes were applied in a one-step staining procedure followed by a single-step ACE amplification. Histone H3 was included in each imaging round as a fiducial marker to facilitate 3D co-registration and spatial analysis, following previously published approaches (**Fig. S19c** and **S19d**)^56,57^. We subsequently analyzed the 3D distribution of molecules, cells, and overall tissue organization. This strategy enabled successful co-registration and analysis of ∼3 million cells across the tissue volume (**Fig. 8a, d, Fig. S31-S32**, and **Video S2**).

**Figure 8.**
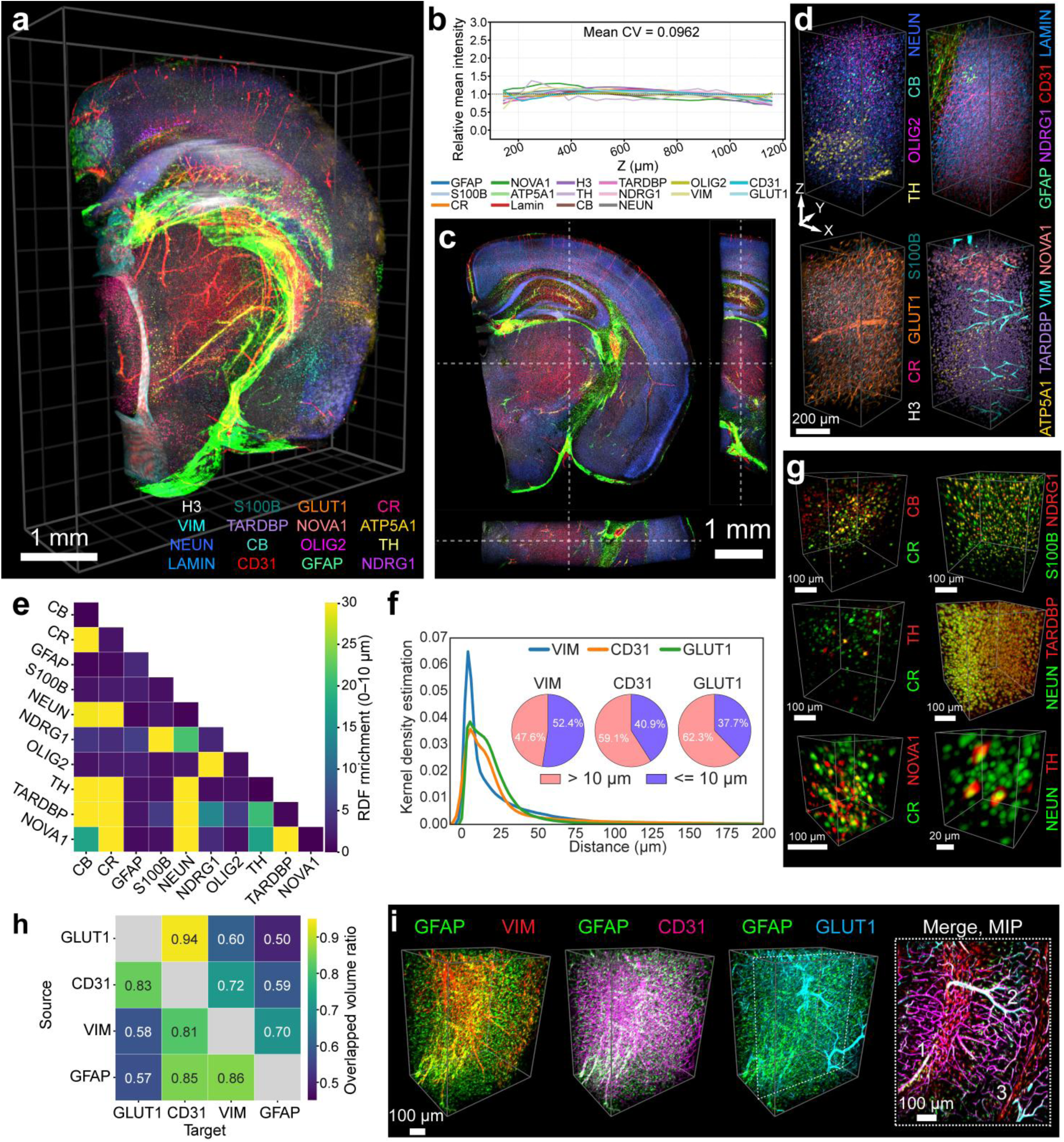
ACE multiplexed imaging of a 1 mm thick mouse coronal section. (**a**) Co-registered image of all 16 markers of a 1-mm-thick mouse coronal block. n = 3,464,350 cells. (**b**) Staining intensities across 1 mm z-depth for all 16 targets. The analyzed tissue window extended from slice 73 (127.75 μm) to slice 671 (1174.25 μm), with both boundaries defined as the positions at which the tissue cross-sectional area reached 10% of its maximum. Solid lines show the relative mean intensity within consecutive valid 20-μm z-bins. All individual profiles were normalized by their respective mean intensity across valid bins (dashed line at y = 1). The overall mean coefficient of variation (Mean CV = 0.0962) indicates highly uniform volumetric signal across the depth profile. (**c**) The cross-section of tissue showing uniform signal amplification. In ECi, the tissue shrunk by ∼ 20%. (**d**) Zoom-in view of subregions for different targets. (**e**) Neighborhood analysis of the entire block using 10 cell-type markers. n = 3,464,350 cells. (**f**) Spatial proximity analysis between target cells and three vascular markers: VIM, CD31, and GLUT1. Kernel density estimation curves show the distribution of shortest distances from target cells to each marker. Inset pie charts summarize the proportion of cells located within (≤10 µm, blue) and beyond (>10 µm, pink) this threshold. n = 234,916 GFAP-positive cells. (**g**) Imaging validation of the neighborhood analysis. (**h**) Heatmap of pairwise overlapped volume ratios among vascular and glial markers in a cropped hippocampus region: GLUT1, CD31, VIM, and GFAP. Each cell represents the fraction of the volume of the row marker that overlaps with the column marker, with values color-coded and numerically labeled. n = 5,880 GLUT1, 8,752 CD31, 7,843 VIM and 5,014 GFAP segmented objects. (**i**) 3D spatial relationship between astrocytes (GFAP) and vascular markers in a representative brain region. Left to right: GFAP is shown with VIM, CD31, and GLUT1. The fourth panel (far right) shows a maximum intensity projection (MIP) over 100 µm of the dashed square region from the third panel, visualizing complex astrocyte-vasculature spatial proximity. White numbers (1-3) label representative vessel structures with distinct marker compositions.

To evaluate the volumetric uniformity of signal amplification across the tissue depth, we systematically profiled relative staining intensities across a 1-mm-thick window. Consistent signal preservation was observed throughout the Z-depth across all 16 evaluated targets, encompassing 14 discrete spot-detection channels (including the pan-neuronal marker NeuN) and 2 volumetric vessel masks (CD31 and GLUT1) (**Fig. 8b**, **Fig. S33a-c**). The target cohort maintained a stable distribution throughout the section, with an overall mean coefficient of variation of 0.096, which is comparable to secondary Ab staining (**Fig. S30a** and **S30d**). The staining pattern also showed uniform signal amplification across the section (**Fig. 8c-d**, and **Video S3**).

Next, we performed cell neighborhood analysis on ten cell-type markers (**Fig. 8e, Fig. S34a**). As expected, we observed strong enrichment between NeuN and other neuronal markers CR, CB, TARDBP, and TH. Subregional annotation and enrichment analysis further revealed region-specific associations (**Fig. 8g, Fig. S34b**). We observed that CR co-stained with CB, TH, and NOVA1 in distinct neuronal populations. S100B co-stained with NDRG1 as they both label myelinating oligodendrocytes. These data also highlight the subcellular resolution of our imaging, with cytoplasmic markers such as CR and TH wrapping around nuclei labeled by the nuclear markers NOVA1 and NeuN.

Astrocytes, key regulators of the blood-brain barrier, were analyzed relative to vasculature. Following 3D surface reconstruction using Imaris, we computed the spatial distances between GFAP⁺ astrocytes and vascular markers VIM, CD31, and GLUT1. Kernel density estimation (KDE) suggested a bimodal distribution of GFAP-to-vessel distances, with peaks near 0 µm and >15 µm (**Fig. 8f**). Quantitatively, we observed that ∼40-50% of GFAP⁺ astrocytes were in close proximity to blood vessels. Notably, the >15 µm KDE peak was attenuated for GFAP-to-VIM distances, suggesting closer association between GFAP and VIM signals.

To further investigate blood vessel heterogeneity, we analyzed the spatial overlap among GFAP, VIM, CD31, and GLUT1 (**Fig. 8h**, and **Video S4**) in the hippocampus region where astrocytes and blood vessels are abundant. We noticed that CD31 and GLUT1 stained more capillaries with fewer neighboring GFAP-positive cells, whereas VIM partially co-stained GFAP-positive astrocytes (**Fig. 8i**). We noticed ∼94% of GLUT1⁺ vessels overlapped with CD31⁺ vessels whereas only ∼83% of CD31⁺ vessels overlapped with GLUT1⁺ vessels. Specifically, we observed multiple vessels to be CD31⁺ while having low GLUT1 staining (**Fig. 8i**). These results indicate CD31 as a pan-blood vessel marker and GLUT1 as a blood-brain barrier marker and highlight the vessel heterogeneity in the region. Additionally, VIM exhibited greater overlap with GFAP than vice versa (**Fig. 8h**). Indeed, we noticed that in the region, many astrocytes were GFAP⁺/VIM⁻, indicating diverse astrocyte subpopulations (**Fig. 8i**).

Together, these results demonstrate the utility of ACE-enabled volumetric imaging for resolving complex cellular and vascular architectures across thick brain tissues with high spatial fidelity. The ability to multiplex across millimeter-scale samples reveals diverse neuronal subtypes, astrocyte-oligodendrocyte and astrocyte-vascular spatial proximity, and highlights region-specific heterogeneity in cell-cell and cell-vessel associations.

## DISCUSSION

In summary, ACE chemistry offers multiplexable, controllable, synchronized, and high S/N amplification under the tested conditions. When applied to imaging, ACE allows for spatial profiling of biomolecules in cells and thick tissue samples. Its versatility extends to applications in imaging and flow cytometry (**Fig. 2f**) and can be integrated with ExM for super-resolution imaging or tissue-clearing protocols for thick samples.

Amplification controllability and evenness can enhance the accuracy and morphological integrity of target spatial mapping. In RNA imaging, probe sets targeting a particular transcript usually consist of > 20 probes. Uneven signal amplification can, in principle, lead to overestimation of RNA puncta as amplification from nonspecifically bound probes can potentially confound quantification. RCA^58^, RNAscope^59^, HCR^30^, and ClampFISH^60^ used proximal probes for generating signals to reduce off-target noise. In ACE, with controllable linear amplification, we can distinguish true signal from off-target noise based on signal intensity and puncta size (**Fig. S3c**).

In protein imaging with signal amplification, uncontrollable amplification can lead to a loss of morphology and structural details. ACE showed even amplification for markers expressed in various cell compartments including the nucleus, cell membrane, and cytosol. We also demonstrated that the integrity of the cytoarchitecture remains intact by imaging tubulin, cytokeratin, and vimentin, all of which retained distinct filamentous morphology.

Several factors underpin the high S/N of ACE. The initiator/ssEXT pairs were engineered using a three-letter code (A, C, T for initiators and A, G, T for ssEXTs) to minimize secondary structure to optimize extension efficiency. The GC content, annealing temperature, and melting temperature of the orthogonal sequences were designed to maintain a < 5 °C temperature difference, facilitating uniform hybridization behavior across pairs and reducing performance variability. The ssEXTs feature a sealed 3’ end, and only dATP, dCTP, and dTTP are utilized for extension, minimizing crosstalk and unintended extension from the ssEXTs. Given that amplification necessitates sequence-matching 3’ single-stranded protruding DNA, nonspecific extension is minimal, as demonstrated by our experiments (**Fig. S4a**). Furthermore, we introduce non-fluorescent mock imagers that share the same sequence composition as the true imagers to suppress nonspecific imager binding (**Fig. 3a**). For Ab staining, the short conjugation initiators further reduce nonspecific binding induced by DNA conjugation^33^. For the antibody panels demonstrated in the manuscript, we selected antibodies with well-characterized, previously published staining patterns and confirmed that the DNA-conjugated antibodies reproduced the expected cell-type, subcellular, and anatomical staining patterns (**Table S7**).

ACE is also compatible with formalin-fixed paraffin-embedded (FFPE) tissue, supporting its potential utility for archived clinical and pathology specimens (**Fig. S35**). We demonstrated this by performing 9-plex ACE protein imaging on a 5-µm human tonsil FFPE section, detecting markers that label diverse immune, epithelial, vascular and proliferative compartments, including CD4, Ki67, CD11b, CK6, CD63, HLA-DR, CD31, CD19 and CD45. The preserved tissue morphology and marker-specific staining patterns suggest that ACE can be applied to standard FFPE samples, expanding its potential use in translational studies and retrospective analysis of clinically annotated tissue cohorts.

In the current studies, we showcased a 67-plex orthogonal extension *in vitro* and 63-plex imaging *in situ*. The ACE protocol’s capabilities are not confined to this multiplexity. Our design pipeline yields 92 orthogonal pairs for 9-nt initiators (**Table S1**). Increasing the initiator length could further expand multiplexing capacity. Our simulations predict more than 150 orthogonal pairs for 10-nt initiators, more than 300 pairs for 11-nt initiators, and more than 500 pairs for 12-nt initiators (**Fig. S1a**). This scalable multiplexity addresses limitations in other amplification techniques, where multiplexity is often constrained by design or reaction kinetics. On the other hand, ACE simplifies the probe design process, reducing the complexities often encountered in traditional methods, thus facilitating practical higher-plex design.

Beyond the applications presented here, ACE amplification holds potential for broader research areas. Future extensions could pair ACE with spectral unmixing and other multichannel strategies to further expand imaging throughput, sensitivity, and multiplexity. We have already shown that ACE can enhance mass cytometry into highly sensitive platforms^61^, and its integration with advanced microscopy opens opportunities for ultrahigh-throughput volumetric imaging at ultra-high scale (1 gigavoxel per second)^62^. Looking ahead, ACE could also be combined with combinatorial barcoding schemes to achieve higher-plex RNA imaging, where its high signal-to-noise amplification could help mitigate current decoding challenges.

The extension efficiency per thermocycle does not reach 100%. The ssEXTs contain two tandem repeats of the reverse complement of the initiator sequence. During each temperature cycle, extension can be blocked if an ssEXT binds to the final two repeats of a concatemer, forming a stable DNA duplex at low temperature. If a newly synthesized terminal repeat remains exposed, extension by an ssEXT becomes probable. Given the stochastic nature of this binding process, we estimate the extension efficiency per cycle to be approximately 50% (**Fig. 1g, h**). This results in a distribution of the concatemer lengths, rather than a fixed length.

A practical limitation of ACE-based RNA FISH is that the FISH probe-target duplex must remain stable during the 58° C cycling step in the ssEXT reaction buffer. This requirement constrains probe design for short or low-GC transcripts, for which it may be difficult to obtain enough probes with adequate melting temperature. Our analysis of Allen Mouse Whole Cortex and Hippocampus SMART-seq neuronal marker genes^63^ suggests that this limitation is modest for many biologically relevant targets: among 2,000 annotation-resolved neuronal marker genes, 1,886 genes (94.3%) had more than 10 ACE-compatible PaintSHOP^64^ probes. Moreover, published MERFISH^65^ probe designs use RNA-targeting regions with melting temperatures above 58° C, indicating that many multiplexed RNA-FISH probe sets are already compatible with this thermal regime. For transcripts that remain challenging, modified bases such as LNA could further increase duplex stability and expand ACE-RNA applicability to shorter or lower-GC targets.

## ACKNOWLEDGMENTS

This research was supported by National Institutes of Health grants UM1MH134812, RF1MH128861, R01MH134803, R01CA283202, RF1MH124606, RF1MH132599, and R01HG012926; the Wyss Institute Molecular Robotics Initiative; and a generous gift from an anonymous donor to the Wyss Institute to support Yin lab research. Portions of the data analyses were conducted on the O2 High Performance Compute Cluster, supported by the Research Computing Group at Harvard Medical School. We gratefully acknowledge the MicRoN (Microscopy Resources on the North Quad) Core for its support and assistance, and the Neurobiology Imaging Facility at Harvard Medical School for imaging support. The authors thank Thomas Ferrante at the Wyss Institute for his assistance in imaging and microscopy. The authors thank Dr. Jinyoung Kang for assistance in expansion microscopy. R.M. was supported by National Science Foundation Graduate Research Fellowship Grant 2140743. R.M. was supported by a National Institutes of Health Molecular Biophysics Training grant (NIGMS T32 GM008313). The cartoons in Fig. 2 and Fig.7 were created with BioRender.com.

## Footnotes

1 **Footnote I.** Raman imaging time estimation: for a 200 μm-thick sagittal mouse brain section measuring 17.8 mm × 12.2 mm (area: 217 mm²), the imaging speed of epr-SRS is 1 min/mm²/channel for every 5 μm Z-step. Therefore, imaging one Z-plane takes 217 min per channel. To cover the full 200 μm thickness (40 Z-steps), the total imaging time is 8,680 min/channel (∼6 d).

2 **Footnote II.** One multiplexing exchange round of mELAST 1mm tissue includes: primary antibody staining (24 h); post-primary wash (12-16 h); secondary antibody and YOYO-1 staining (16-18 h); post-secondary wash (3-4 h); optical clearing and refractive-index matching (1 day); SDS pre-incubation for antibody removal (overnight at 45 °C; ∼12-16 h); thermal elution (1 h at 80 °C); pumping-assisted destaining (3-5 h); SDS removal wash (3-5 h); and buffer exchange prior to the next round (2-3 h). These steps sum to ∼100-116 h (4.2-4.8 days), resulting in a total exchange time of 4-5 days per round (imaging time excluded).

3 **Footnote III.** DNA probes exhibit an effective binding rate ∼100-fold higher than IgG antibodies: 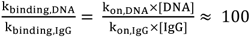. Working concentration: [DNA] ≈ 0.5 μM, [IgG] ≈ 33 nM. Rate constant: kon,DNA ≈ 10^6^ − 10^7^ M^−1^S^−1^, kon, IgG ≈ 10^5^ − 10^6^ M^−1^S^−1^. Separately, DNA diffusion is ∼10-fold higher than IgG diffusion: 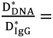 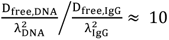. Diffusion coefficient: D ≈ 150 μm²/s, D ≈ 65 μm²/s. Tissue tortuosity: λ ≈ 1.6, λ ≈ 3.1.

