## Supplementary Information for "High-plex protein imaging across thick tissue by synchronized amplification"

### 1 Supplementary Figures

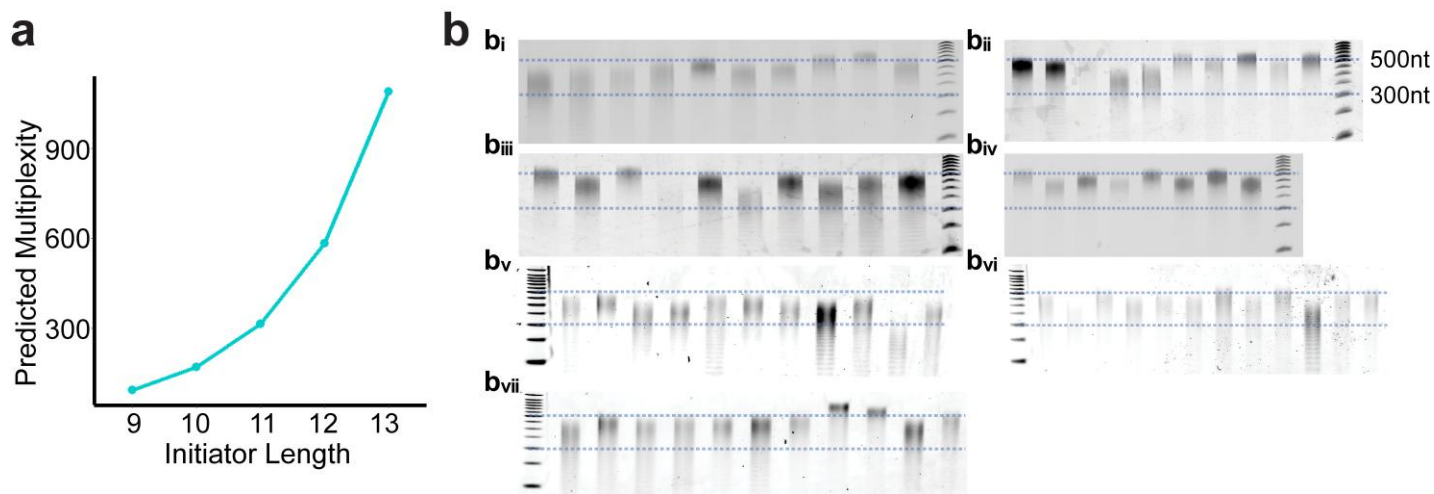

**Figure S1. Orthogonal sequence design and in-tube validation of initiator/ssEXT pairs.** (a) Computational analysis using Seqwalk for the maximum multiplexity for different primer lengths. 9 nt: 93; 10 nt: 170; 11 nt: 315; 12 nt: 585; 13 nt: 1092; 14 nt: 2048; 15 nt: 3,855. (b) Test-tube validation of initiator/ssEXT pairs by gel electrophoresis. Each reaction underwent 80 extension cycles. Samples were divided among seven gels, with lanes ordered from left to right as follows: **b(i)** p.1, p.2, p.68, p.69, p.3, p.4, p.5, p.70, p.71, and p.6; **b(ii)** p.7, p.8, p.9, p.72, p.10, p.11, p.12, p.13, p.14, and p.73; **b(iii)** p.15, p.16, p.74, p.75, p.17, p.76, p.18, p.19, p.20, and p.21; **b(iv)** p.22, p.23, p.24, p.25, p.26, p.27, p.77, and p.28; **b(v)** p.29, p.30, p.31, p.33, p.79, p.34, p.80, p.37, p.38, p.40, and p.41; **b(vi)** p.42, p.43, p.44, p.45, p.47, p.48, p.49, p.50, p.51, p.52, p.53, and p.54; and **b(vii)** p.55, p.56, p.59, p.60, p.61, p.62, p.63, p.64, p.65, p.66, and p.67. Blue dashed lines indicate the 300–500-nt size range. A total of 72 initiator/ssEXT pairs were tested. Of these, 58 exhibited greater than 99% specificity in the sequencing experiment shown in Fig. 1f; the remaining 14 pairs were derived from previously reported SABER sequences<sup>1,2</sup>. Barcode sequences are provided in Table S1.

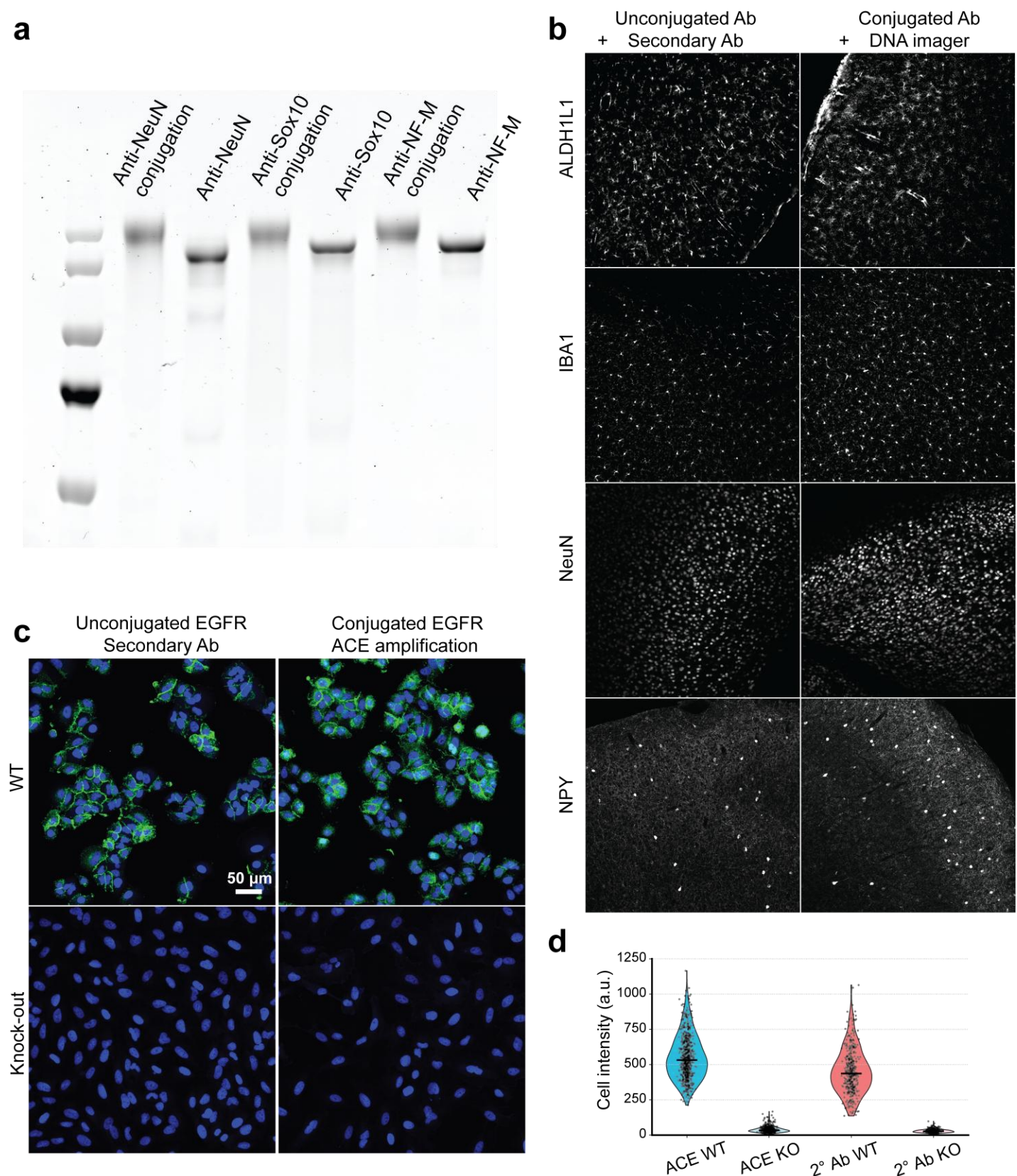

**Figure S2. Antibody conjugation and validation of ACE protein detection.** (a) Gel band-shift assay results showing effective Ab-initiator conjugation. (b) Comparison of unconjugated parent antibodies detected by fluorescent secondary antibody and DNA-conjugated antibodies detected by DNA imager readout for representative mouse brain markers. Matched anatomical regions are shown for ALDH1L1, IBA1, NeuN and NPY. The conjugated and unconjugated antibodies produced similar cellular and anatomical staining patterns. (c) EGFR staining in wild-type (WT, EGFR+) and EGFR-knockout (KO, EGFR-) cells using either the ACE amplification or a standard secondary antibody. (d) Violin plots displaying the single-cell background-subtracted

mean fluorescence intensities of EGFR staining in wild-type (WT, EGFR+) and EGFR-knockout (KO, EGFR-) cells using either the ACE amplification (blue/cyan) or a standard secondary antibody detection system (secondary Ab, red/peach). All measurements were obtained using a low-excitation laser power (640 nm, 10% laser power). Black horizontal bars represent the median value of the cell population. Individual data points (single cells) are overlaid as black jitter points (ACE WT: n=660 cells; ACE KO: n=517 cells; secondary Ab WT: n=400 cells; secondary Ab KO: n=379 cells; pooled from N=4 independent fields of view).

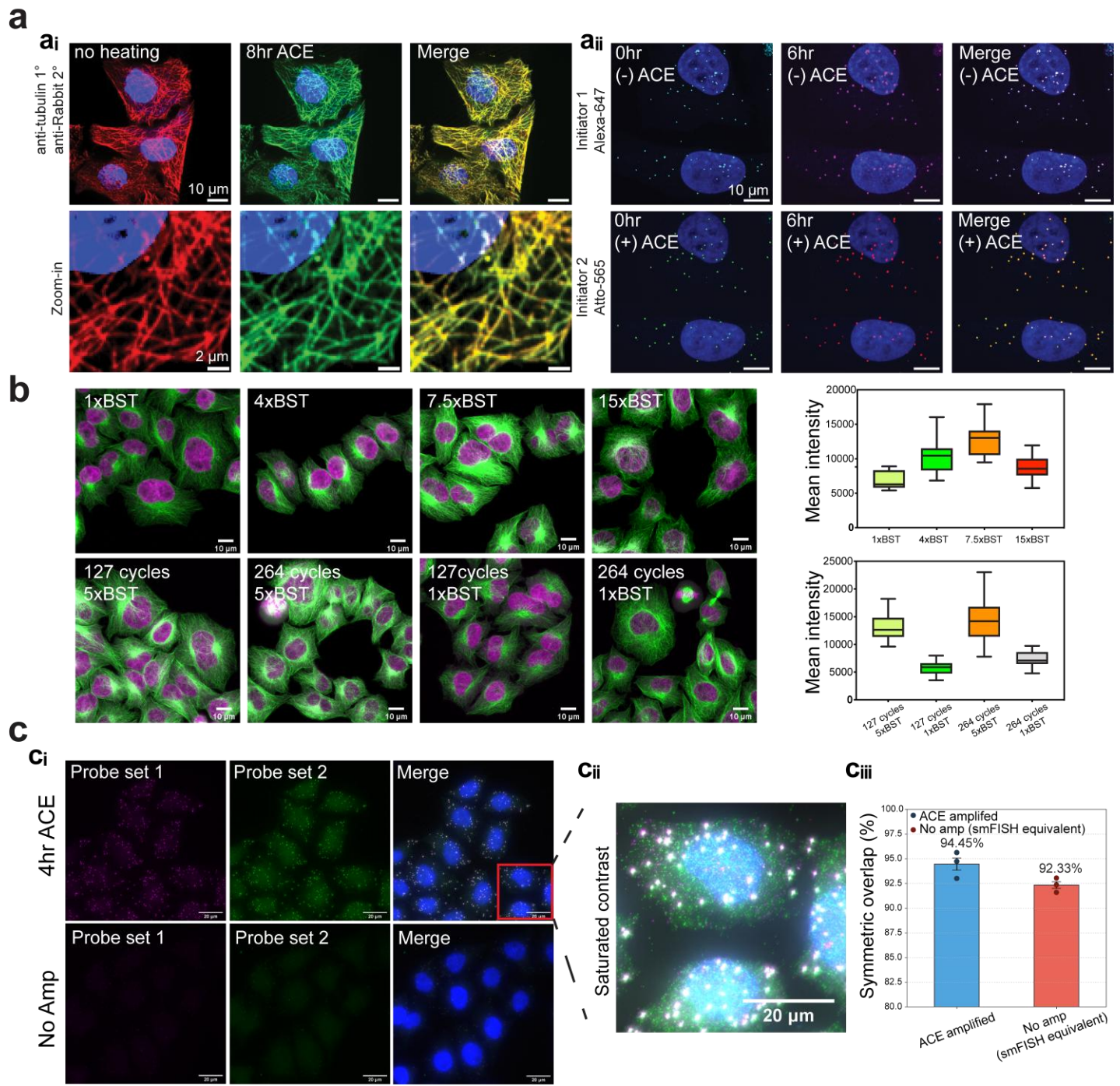

**Figure S3. Signal integrity, polymerase optimization, and RNA co-localization.** (a) The effect of ACE thermal cycling on signal integrity for **a(i)** protein and **a(ii)** RNA. For protein, we stained the cells with primary and secondary Abs and performed post-fixation followed by 8 h of ACE thermal cycling. For RNA, we used two sets of ACE-barcoded probes targeting APC transcripts and performed ACE. (b) Titrating Bst DNA polymerase concentration for optimal *in situ* ACE. scale bars, 10 μm. Top, cells amplified using 1×, 4×, 7.5× or 15× Bst. Bottom, cells amplified with 264 or 127 cycles using 5× or 1× Bst. Box plots show mean fluorescence intensity per image; center lines indicate medians, boxes indicate interquartile ranges and whiskers indicate the data range. $n = 15$ quantified images per condition. (c) ACE amplification of 2 probe sets targeting the same gene **c(i)** a zoom-out view and **c(ii)** a zoomed-in view with contrast adjusted to visualize nonspecific primary-probe signal. At this contrast, the specific (true) signal is saturated. We observed high signal overlaps. **c(iii)** Symmetric co-localization of detected RNA puncta between independent probe channels targeting the same transcript. Bars represent the mean co-localization rates for the ACE-amplified protocol (blue, $n=3$ independent fields of view) and no ACE amplification as the smFISH equivalent (red, $n=3$ independent fields of view). Individual data points (fields of view) are plotted as filled circles. Error bars represent mean ± s.e.m.

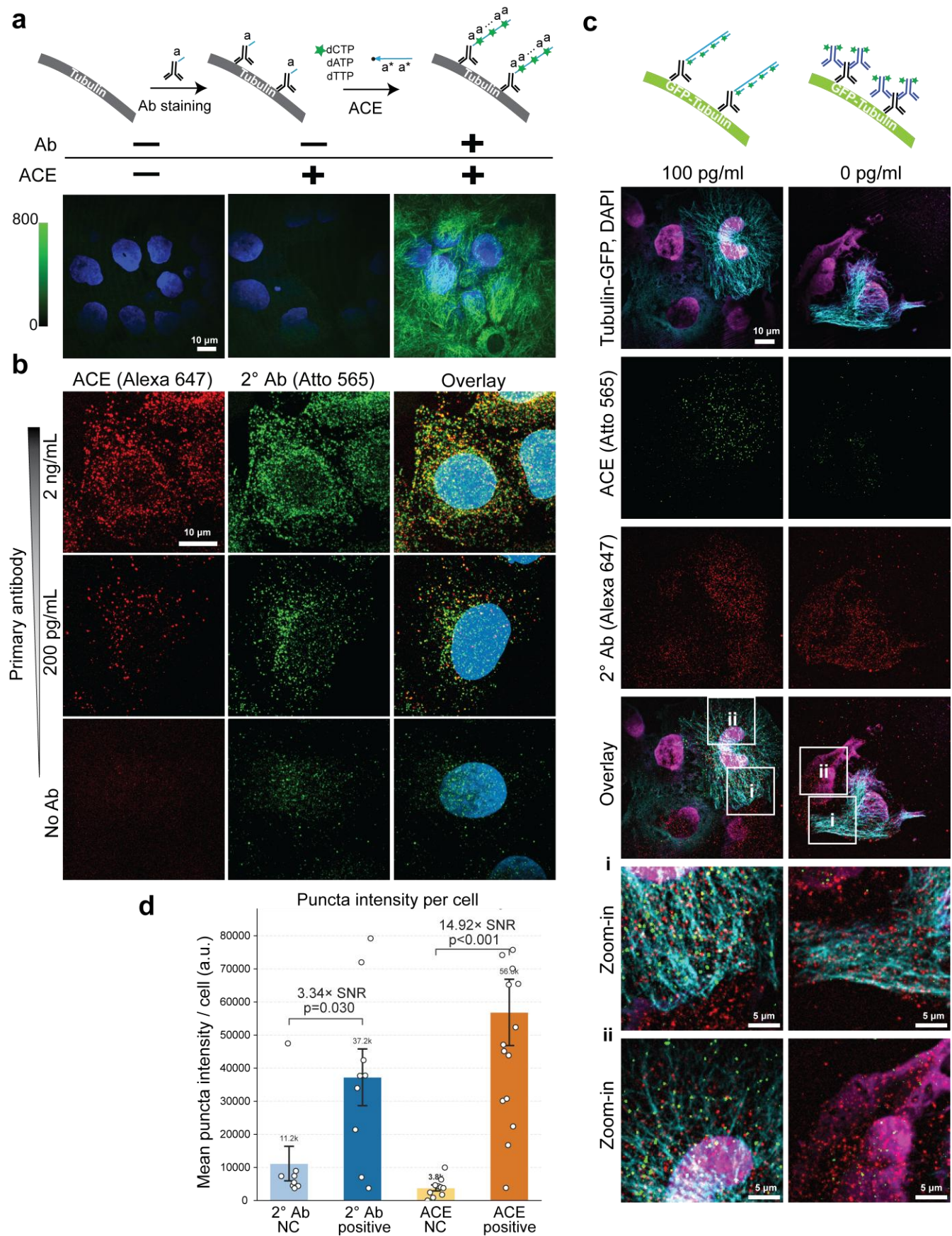

**Figure S4. FAM-dCTP labeling and comparison of ACE with secondary-antibody detection.** (a) Representative images showing FAM-dCTP-labeled DNA after ACE amplification of initiator-conjugated anti-β-tubulin Ab in HeLa cells. The ACE-amplified sample without primary antibody showed background comparable to the no-ACE, no-extension control. (b) Signal comparison of ACE and secondary Ab for a serial

dilution of primary Anti-β-tubulin Ab in HeLa cells. **(c)** Signal comparison of ACE and secondary Ab in GFP-β-tubulin transfected HeLa cells. Diluted primary anti-GFP Ab were used to stain the cells. **(d)** Quantification of mean puncta intensity per cell (GFP-high transfected cells) for conventional secondary antibody staining (blue) and ACE amplification (orange) in the presence (positive, dark bars) or absence (negative control [NC], light bars) of the anti-GFP primary antibody. Data are presented as mean ± s.e.m. across independent images (biological replicates; secondary antibody NC: n=8 images, secondary antibody positive: n=9 images, ACE NC: n=10 images, ACE positive: n=16 images). Circles represent individual image means. Signal-to-noise ratio (SNR) values indicate the fold-enrichment of positive over NC group means. Statistical significance was evaluated using a one-sided Mann-Whitney U test.

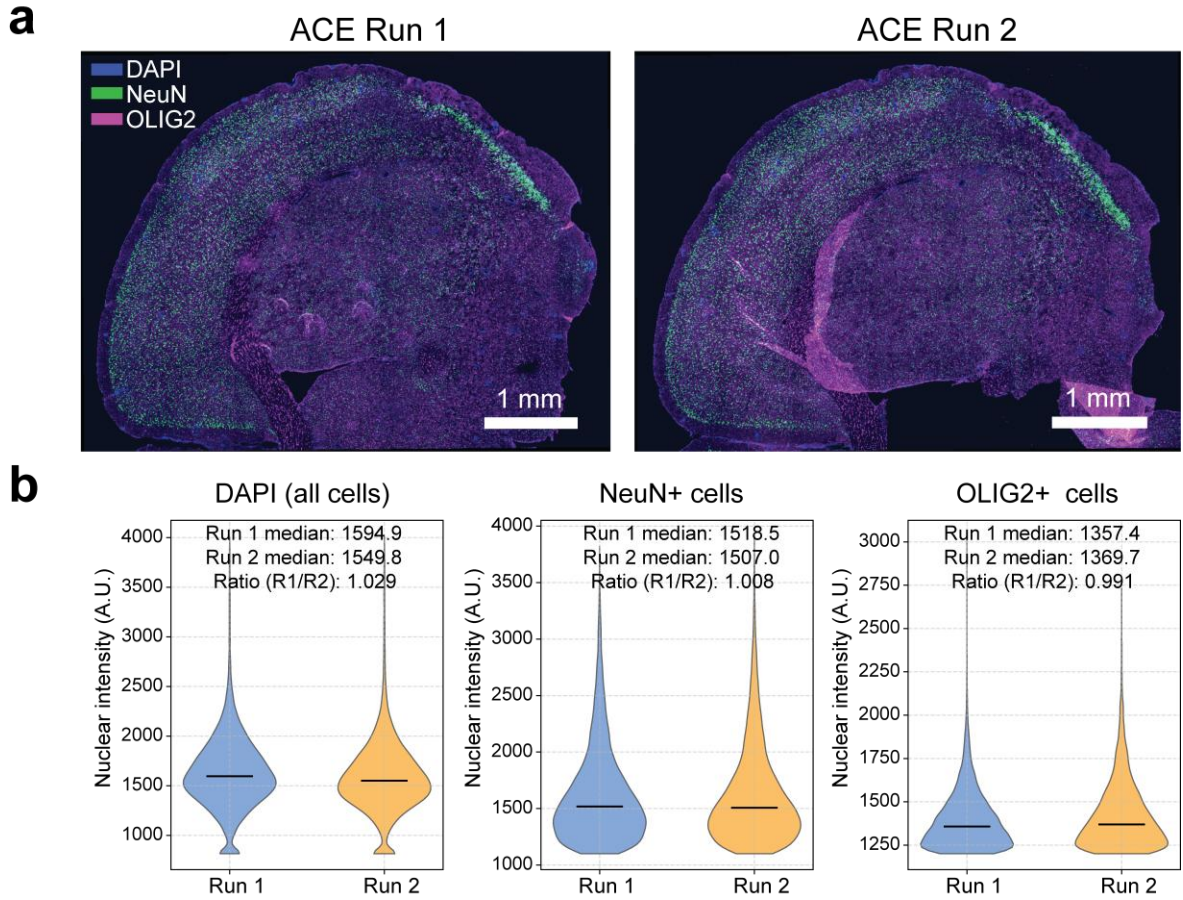

**Figure S5. Reproducibility of multiplexed staining across adjacent mouse brain sections. (a)** Three-color multiplexed fluorescence composite images of adjacent mouse brain coronal sections from run 1 (left) and run 2 (right). Blue, DAPI (nuclei); green, NeuN (neurons); magenta, OLIG2 (oligodendrocytes). Fixed contrast limits are applied across both runs. **(b)** Per-cell fluorescence intensity violin plots of positive cell populations for DAPI (Run 1: $n=43,603$ cells, Run 2: $n=41,974$ cells), NeuN ($n=23,013$ vs. $n=24,024$), and OLIG2 ($n=6,127$ vs. $n=7,696$). Medians are indicated by horizontal black bars. Individual panels show independent vertical axes to highlight distribution morphology. Insets display median values and the respective run 1 / run 2 intensity ratios.

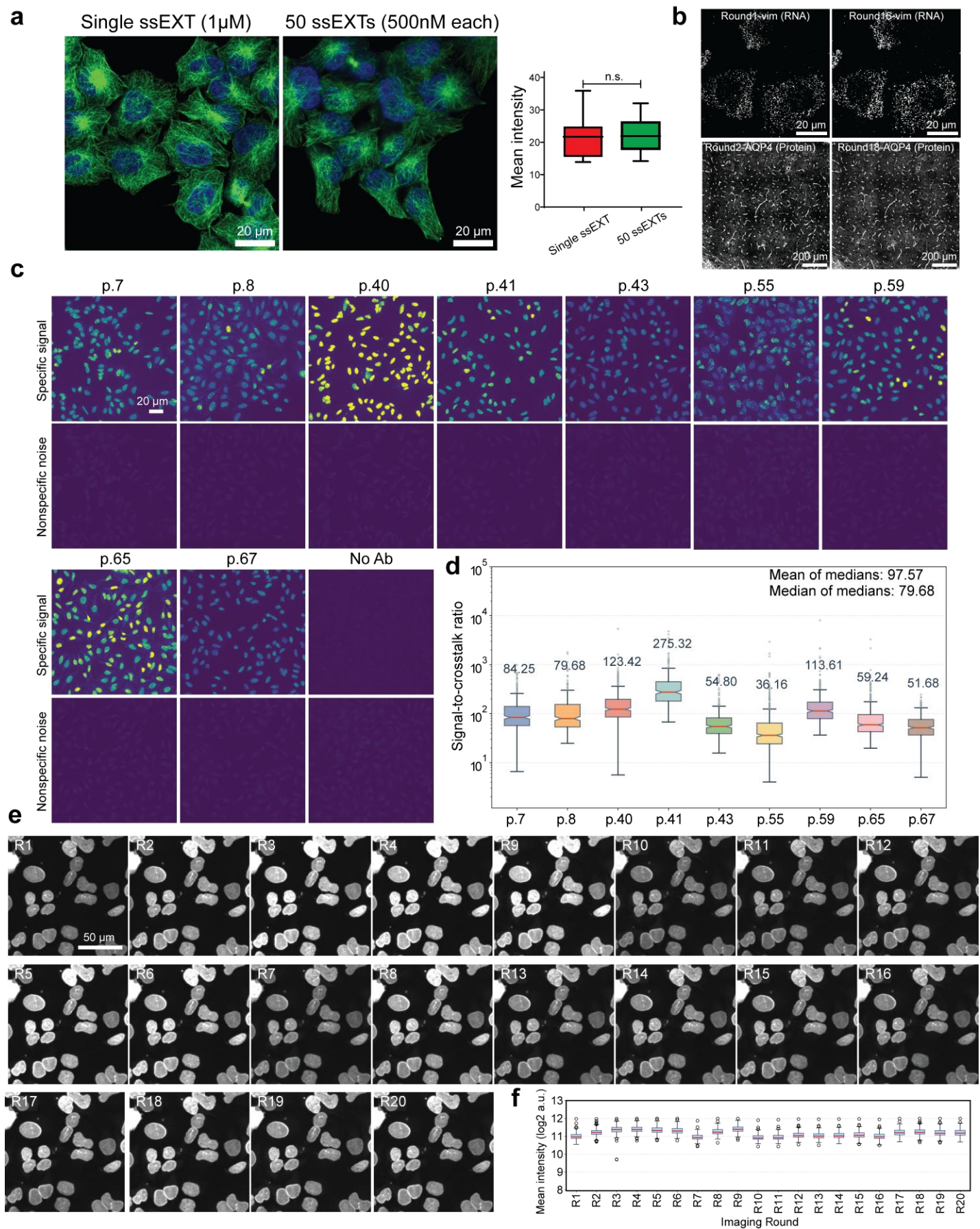

**Figure S6. Multiplex amplification, imager exchange, and in situ barcode specificity.** (a) Effect of multiplex ACE on a single target *in situ*. We ACE amplified HeLa cells stained with initiator-conjugated Anti-β-tubulin Ab with either a single ssEXT or 50 ssEXTs. Scale bars, 20 μm. Box plots show mean fluorescence intensity per image; center lines indicate medians, boxes indicate interquartile ranges and whiskers indicate the data range. $n = 15$ images per condition. n.s., not significant. (b) The effect of imager exchange rounds on signal morphology and integrity for (i) an RNA target in cultured Py2T cells and (ii) a protein target on a mouse brain section. (c) *In situ* validation of ACE barcode sequence specificity using an imager leave-one-out assay. Representative maximum-intensity projections (MIPs) showing specific Lamin A/C immunofluorescence (left panels) and residual nonspecific crosstalk signal (right panels) across HeLa cells stained with Lamin A/C conjugated to nine representative ACE barcodes: p.7, p.8, p.40, p.41, p.43, p.55, p.59, p.65 and p.67. The corresponding extension specificities were 100%, 99.11%, 99.99%, 99.99%, 99.98%, 99.05%, 99.84%, 99.32% and 99.99%, respectively. The negative control was performed with omission of the primary antibody (labeled as “No Ab”). Specific signal and nonspecific noise channels are displayed under the same contrast scale (intensity range, 800-3,500) to allow direct visual comparison. Scale bar, 20 μm. (d) Quantitative single-cell sum-based signal-to-crosstalk ratio distributions across the nine ACE barcodes. Box plots represent the interquartile range (IQR, 25th to 75th percentiles), with the center line denoting the median and notches indicating the 95% confidence interval of the median. Whiskers extend to $1.5 \times$ IQR, and individual data points are shown as gray circles. $n = 382, 414, 491, 324, 361, 602, 404, 664$ and $358$ cells for p.7, p.8, p.40, p.41, p.43, p.55, p.59, p.65 and p.67, respectively. Text values above each whisker denote the median signal-to-crosstalk ratio for each barcode. The legend indicates the mean of the medians (97.57) and median of the medians (79.68) across all nine tested barcodes. (e) Representative cropped and aligned fluorescence micrographs showing Lamin A/C signal (640 nm channel) in the same field of view across 20 consecutive imaging rounds (R1 to R20). Inter-cycle stage translation drift was computationally corrected using a hybrid DAPI-based phase cross-correlation algorithm. Scale bar in R1, 50 μm. All panels are shown under a fixed intensity display range to visually demonstrate the intensity. (f) Box plot showing the distribution of per-nucleus independent log<sub>2</sub> mean Lamin intensity across rounds ($n \approx 240$ nuclei segmented per round). Median intensities (red horizontal lines), interquartile ranges (blue boxes), and outliers (black dots).

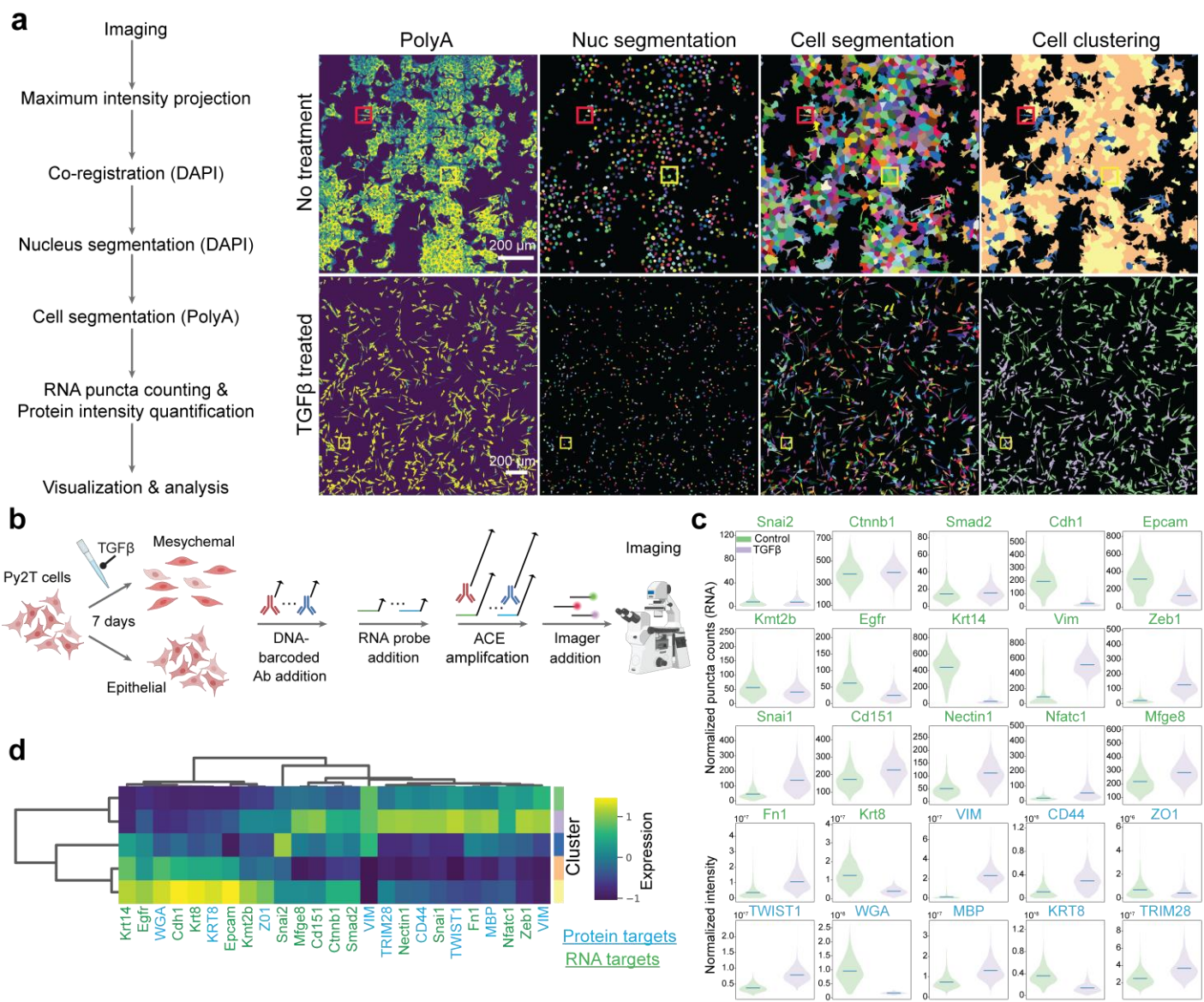

**Figure S7. Py2T cell segmentation, clustering, and expression analysis.** (a) The cell segmentation and cell clustering pipeline for Py2T cell analysis and the images show the entire imaging areas for the two conditions. We showed the PolyA and the color-coded nucleus segmentation images, the final color-coded cell segmentation, and the cell typing after clustering. The yellow squares outline the fields of view shown in Fig. 3c and the red squares outline the fields of view shown in Fig. 3h. (b) Schematic for Py2T EMT imaging studies using ACE. (c) The cell size normalized abundance of the selected targets in the two different groups. Normalized puncta counts were used to quantify the abundance of most RNA targets. Normalized intensity was used to quantify the expression of protein targets. For Fn1 RNA and Krt8 RNA, normalized intensity was used because expression is too high to resolve individual puncta. $n = 1,699$ cells. (d) The expression profile of the 25 targets for the 5 different clusters. Center lines indicate medians. $n = 1,699$ cells.

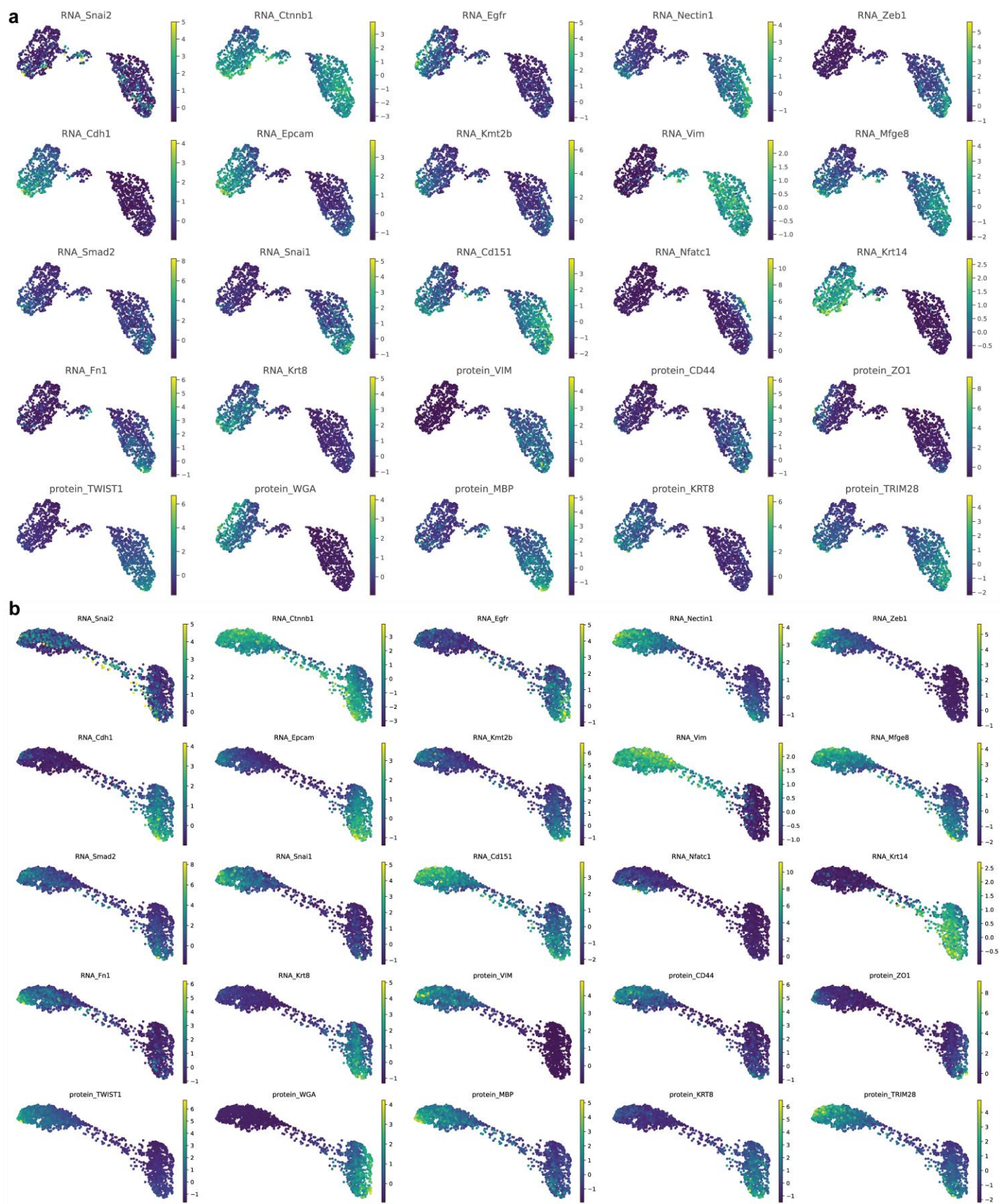

**Figure S8. Expression profiles of 25 Py2T targets on UMAP and pseudotime.** The expression profile of the 25 targets in the Py2T analysis on (a) UMAP in Fig. 3d and (b) pseudotime in Fig. 3g. n = 1,699 cells.

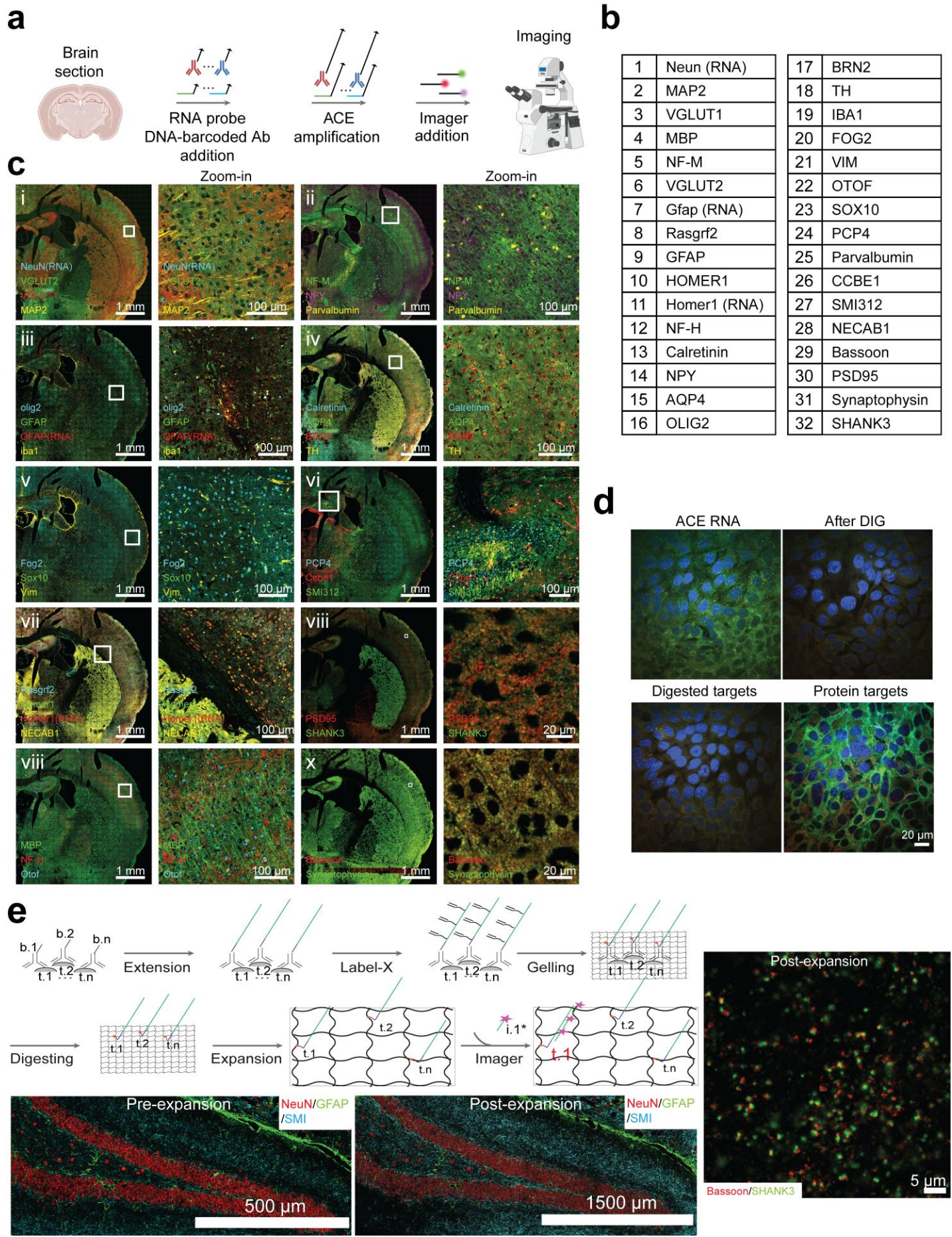

**Figure S9. Multiplex imaging, RNA removal, and expansion microscopy.** (a) Workflow for ACE imaging in mouse brain sections. RNA probes and DNA-barcoded antibodies are added, followed by ACE amplification and sequential fluorescent imager exchange. (b) List of the 32 targets imaged, comprising 29 proteins and 3 RNAs. (c) 32-plex ACE imaging (29 protein targets and 3 RNA targets with a single amplification step) of a 10-μm mouse coronal section. (d) Validation of successful removal of RNA signal after RNase H + RNase I<sub>f</sub> digestion. In Py2T cells, we first imaged the ACE-amplified RNA signal. We then used RNase H + RNase I<sub>f</sub> to digest the RNA and check the removal of signals. We then labeled the cells with Abs carrying the same initiators used by the RNA probes and imaged the ACE-amplified sample again. (e) Schematics and results for ACE-ExM. Melphalan-X was used to crosslink the concatemers generated by ACE to the hydrogel; the gel was then expanded and subjected to imager exchange.

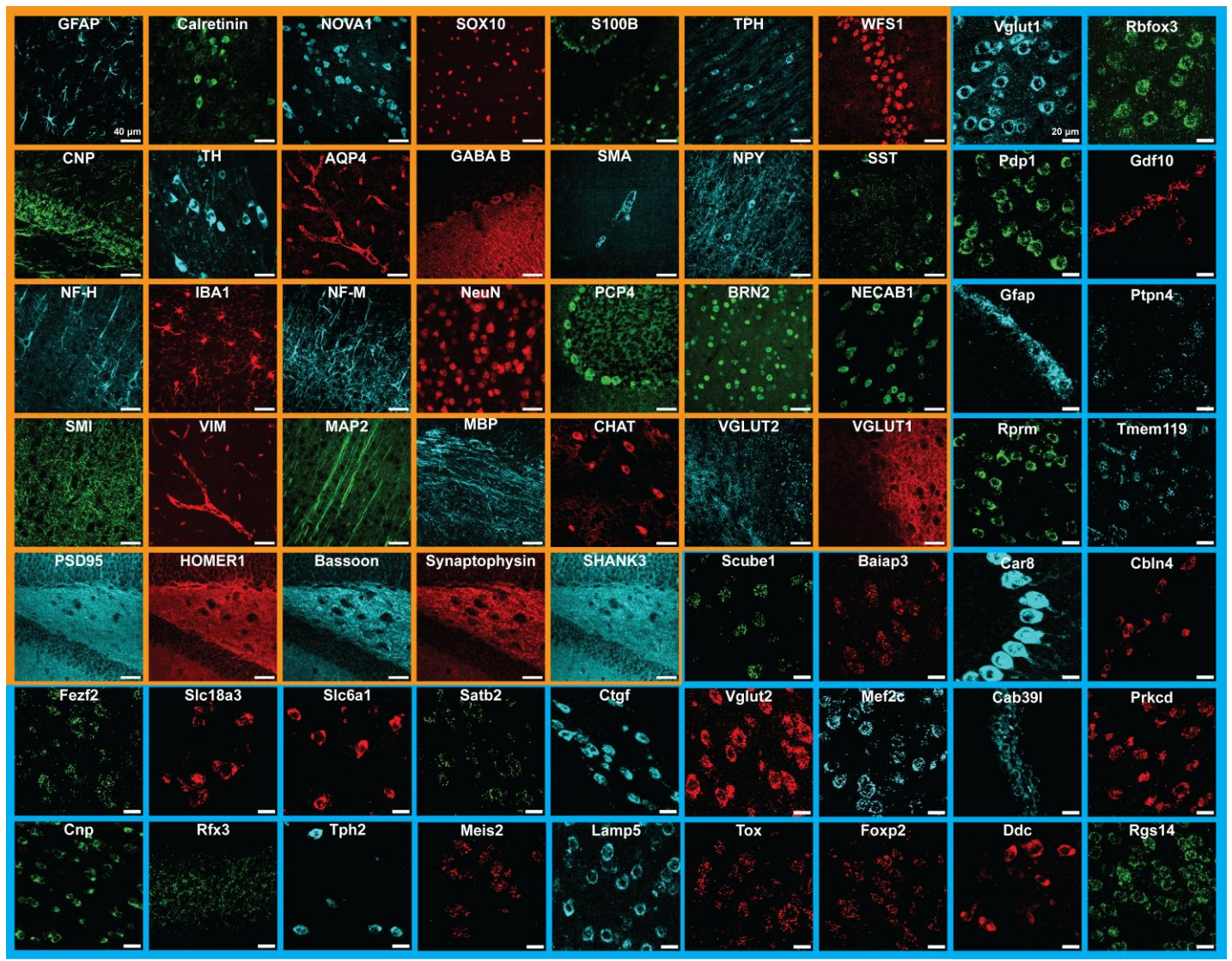

**Figure S10. Zoom-in views of 63 targets in the mouse sagittal section.** Zoom-in views of all 63 targets imaged in the mouse sagittal section. All protein targets are labeled orange and RNA targets labeled blue.

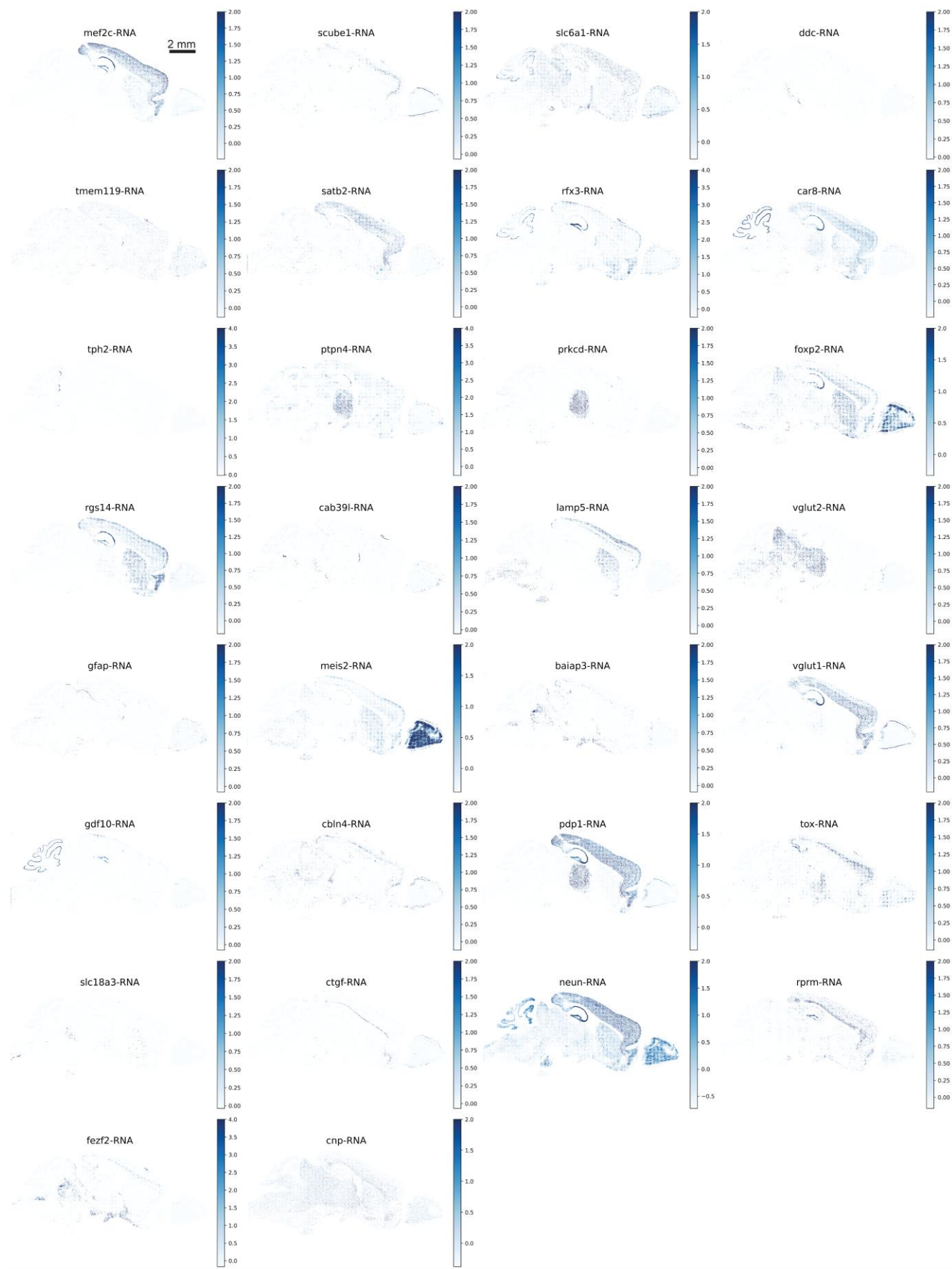

**Figure S11. Whole-section expression profiles of 30 RNA targets.** The expression profile of the 30 RNA targets at the whole-section level. Expression levels were normalized by cell size and subsequently scaled. $n = 113,547$ cells.

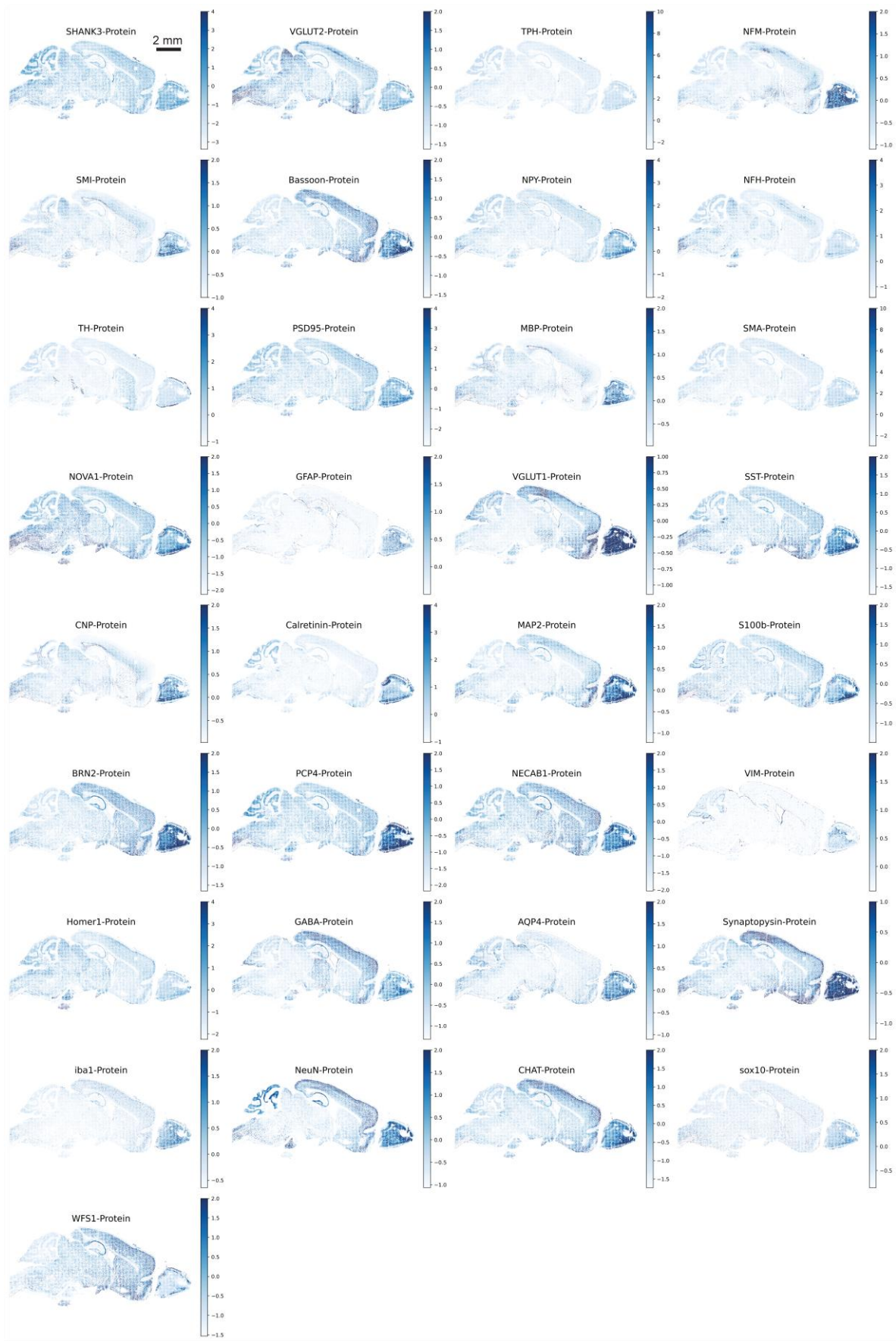

**Figure S12. Whole-section expression profiles of 33 protein targets.** The expression profile of the 33 protein targets at the whole-section level. Expression levels were normalized by cell size and subsequently scaled. $n = 113,547$ cells.

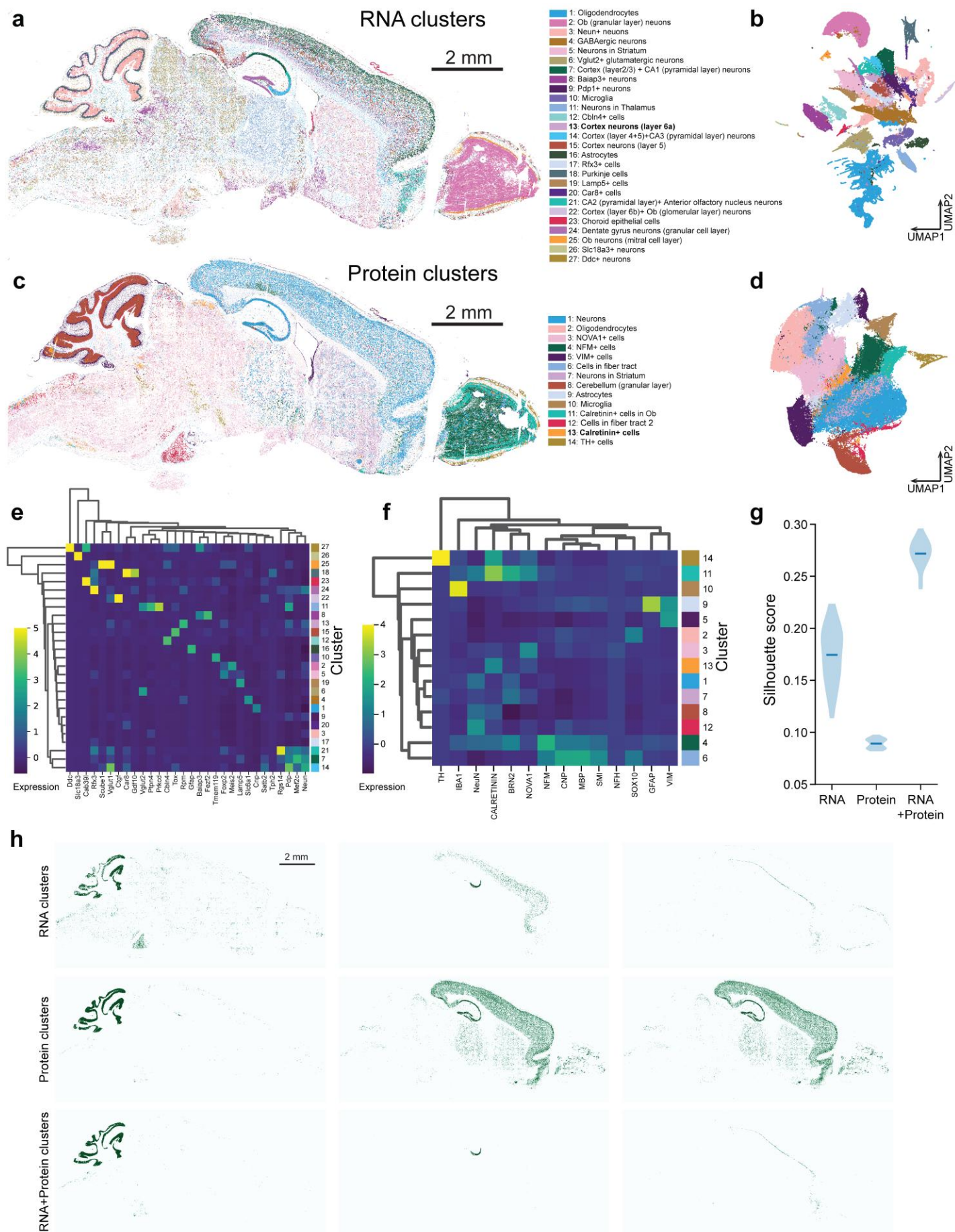

**Figure S13. Cell-type maps and comparison of RNA and protein clustering.** (a) Molecular cell-type maps and the annotations of a mouse sagittal section colored by clusters generated using RNA targets. Each spot represents one cell. n = 113,547 cells. (b) UMAP plot of 27 clusters with the same color code as in Fig. S13a. Clusters were generated using RNA targets. n = 113,547 cells. (c) Molecular cell-type maps and the annotations of a mouse sagittal section colored by clusters generated using selected protein targets. Each spot represents one cell. n = 113,547 cells. (d) UMAP plot of 14 clusters with the same color code as in Fig. S13c. n = 113,547 cells. (e) The expression distribution and hierarchical clustering of the RNA targets for the 27 different clusters with the same color code in Fig. S13a. n = 113,547 cells. (f) The expression distribution and hierarchical clustering of the selected protein targets for the 14 different clusters with the same color code as in Fig. S13c. n = 113,547 cells. (g) Cluster quality comparison using silhouette score between RNA-only, protein-only and RNA + protein clustering projected on UMAP 30 times. Horizontal lines indicate medians. (h) Representative clustering of RNA clusters, protein clusters, and RNA + protein clusters. n = 113,547 cells.

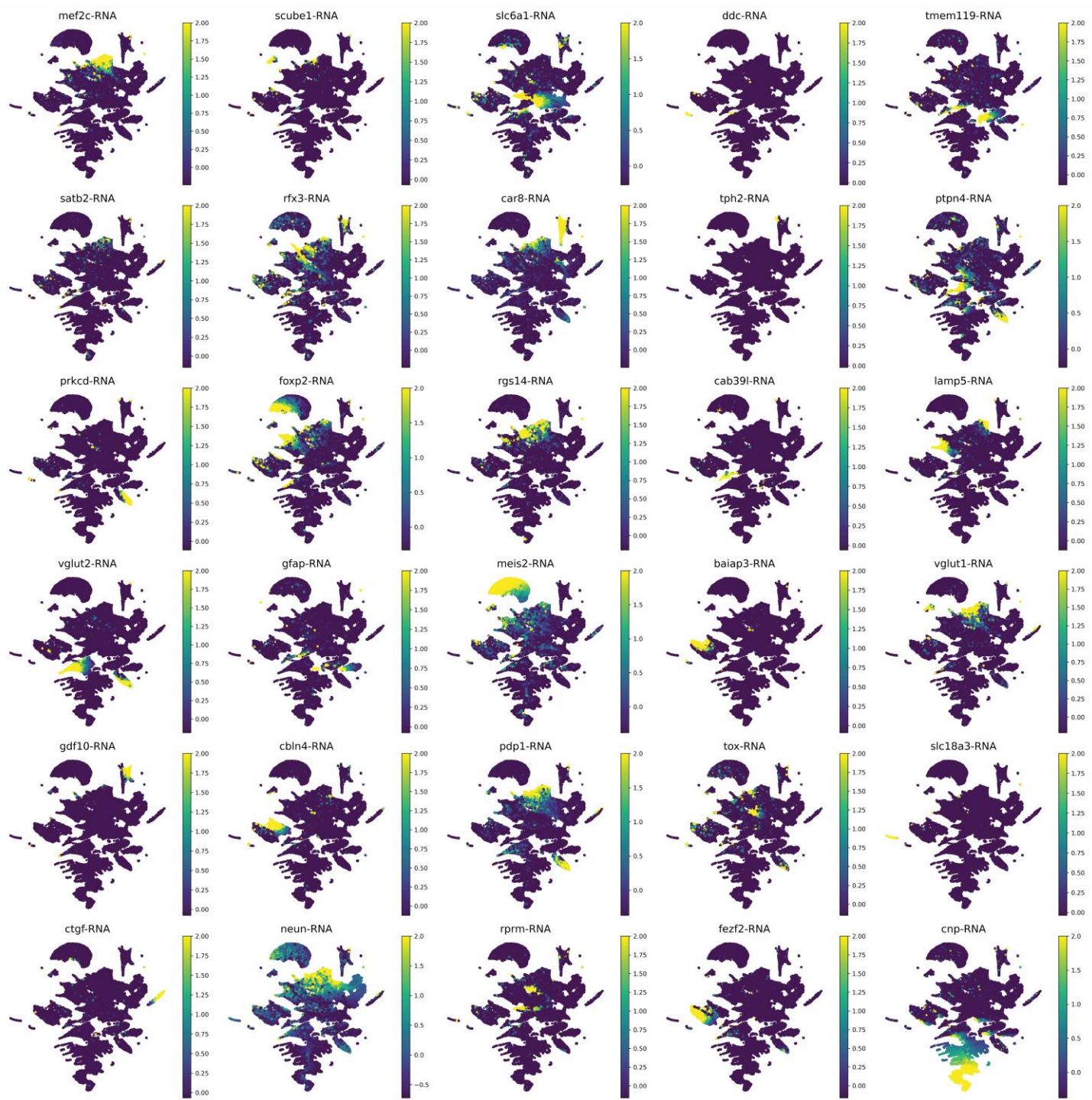

**Figure S14. Expression of 30 RNA targets on the RNA-based UMAP.** The expression of all 30 RNA targets on the same UMAP in Fig. S13b. $n = 113,547$ cells.

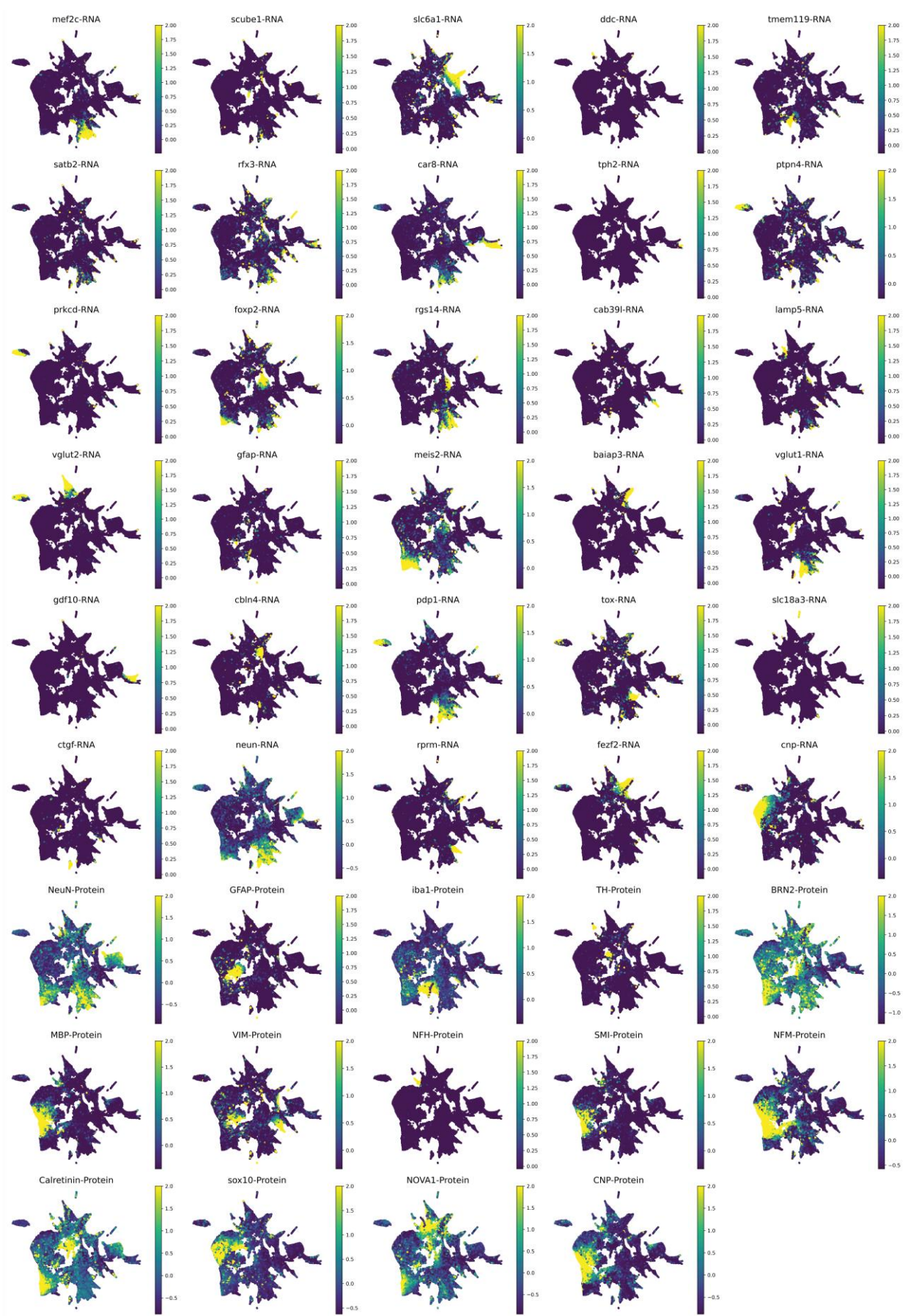

**Figure S15. Expression of 30 RNA and 14 protein targets on the combined UMAP.** The expression of the targets on the same UMAP in Fig. 5b(ii). n = 113,547 cells.

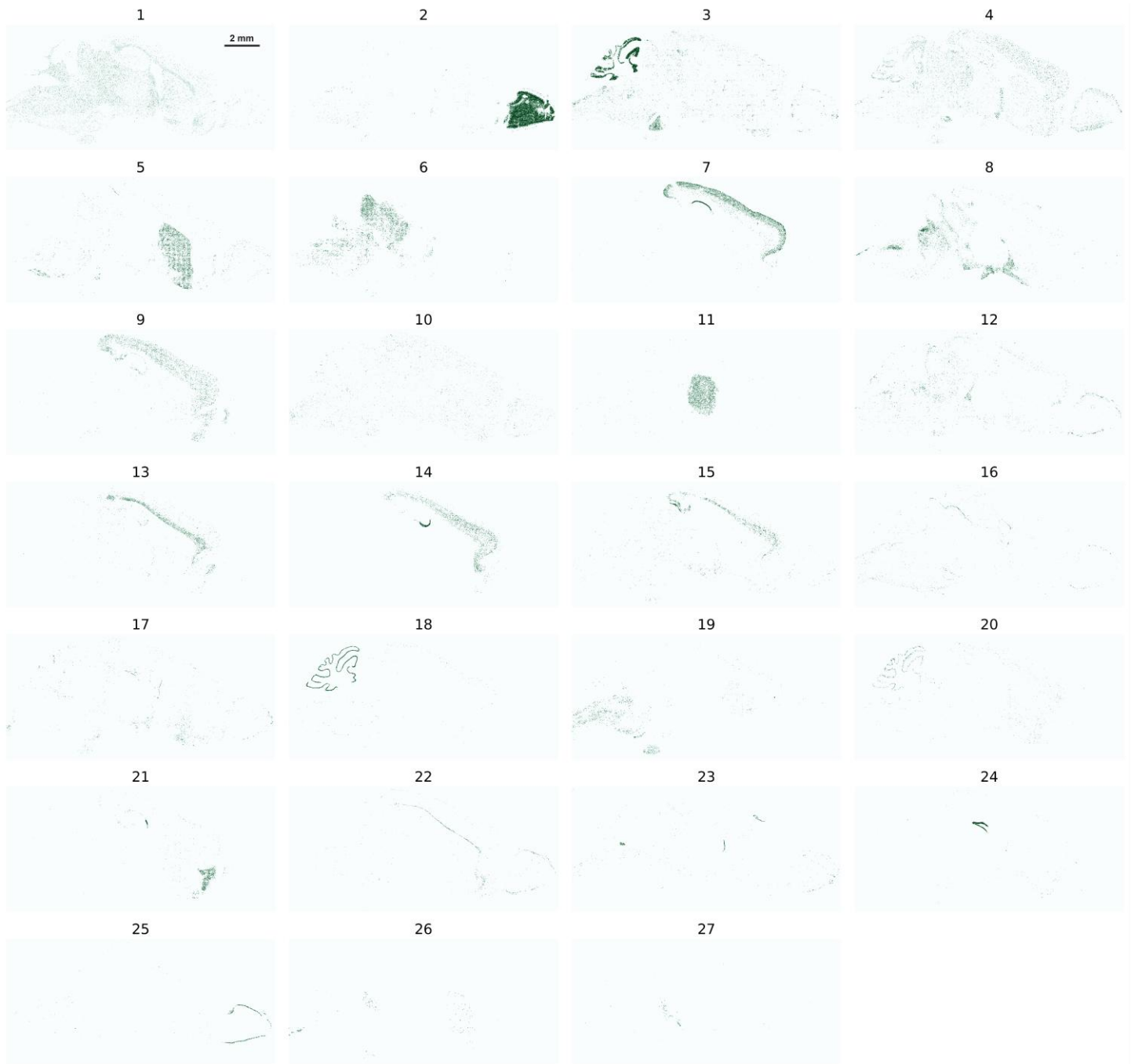

**Figure S16. Tissue projections of 27 RNA-based clusters.** The 27 clusters generated by RNA targets in Fig. S13a were projected on the tissue. $n = 113,547$ cells.

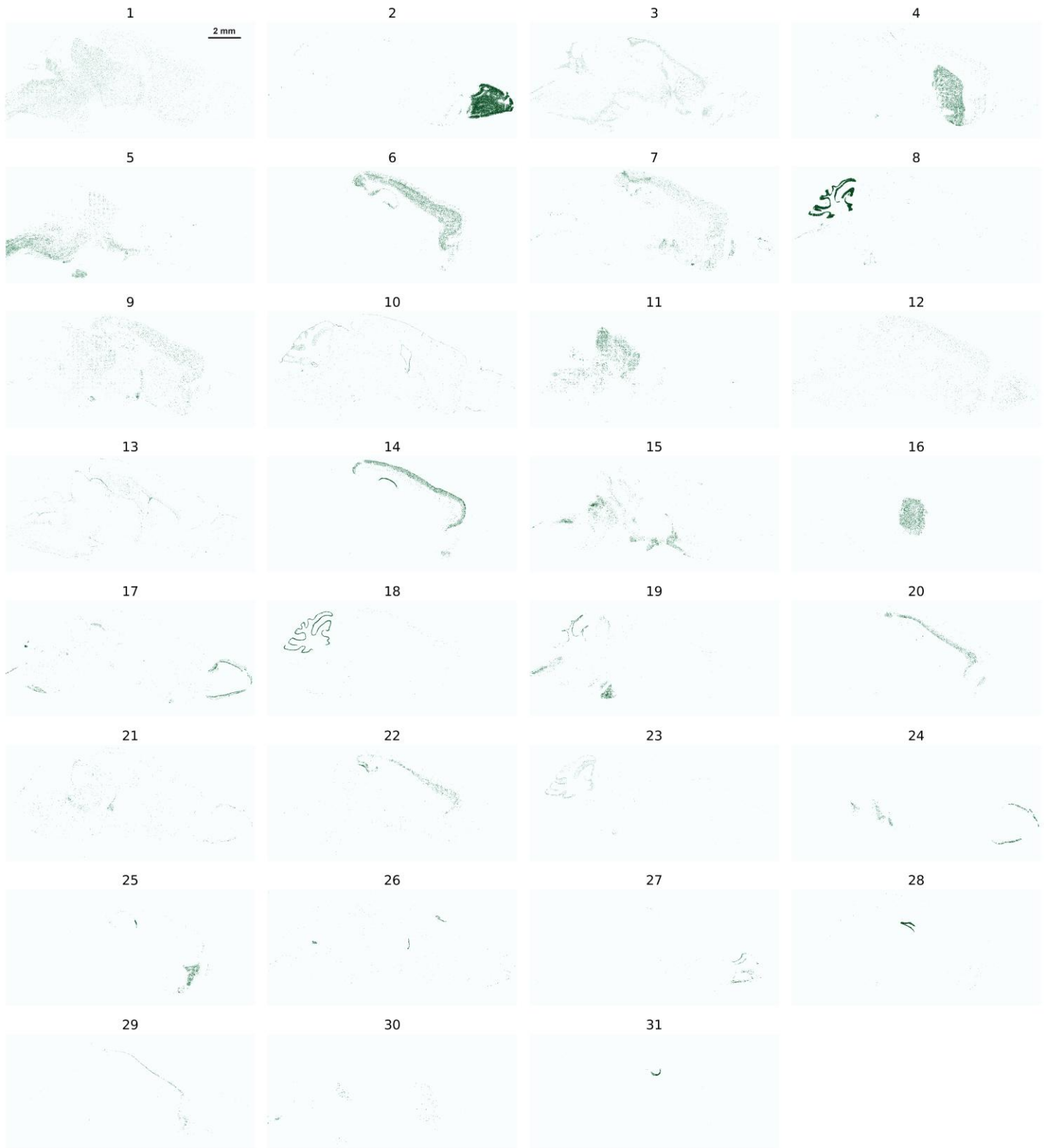

**Figure S17. Tissue projections of 31 RNA and protein clusters.** The 31 clusters generated by RNA + protein targets in Fig. 5b(i) were projected on the tissue. $n = 113,547$ cells.

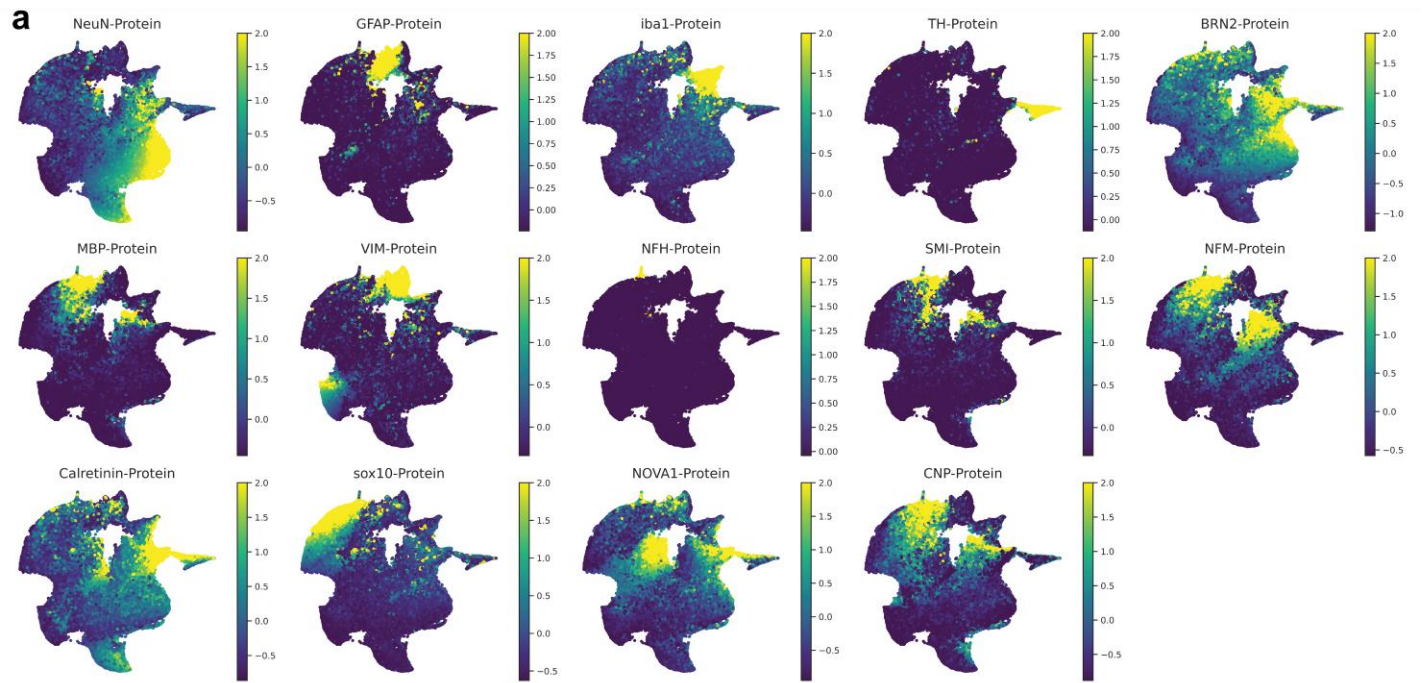

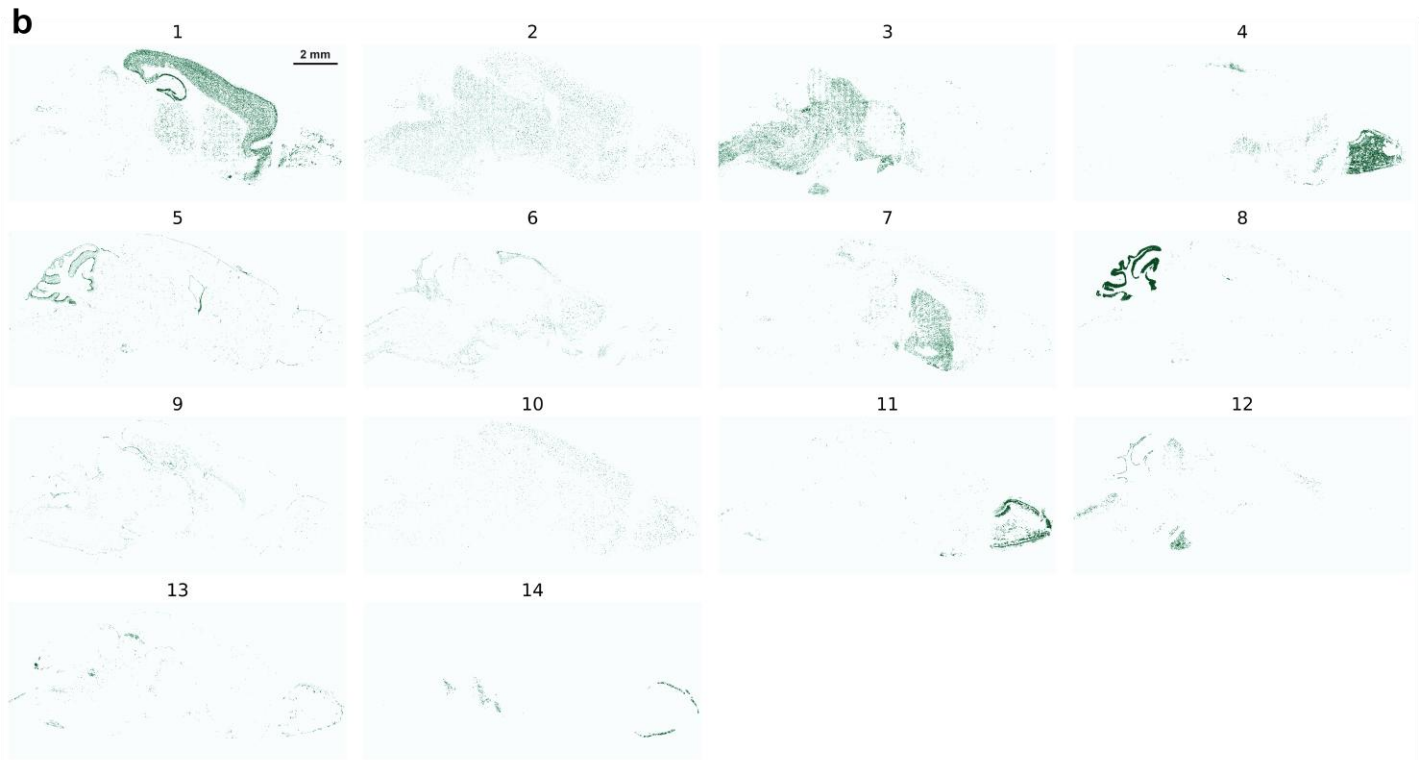

**Figure S18. Protein expression on UMAP and tissue projections of 14 protein-based clusters. (a)** The expression of the selected 14 protein targets on the same UMAP in Fig. S13d. $n = 113,547$ cells. **(b)** The 14 clusters generated using protein targets in Fig. S13c were projected on the tissue. $n = 113,547$ cells.

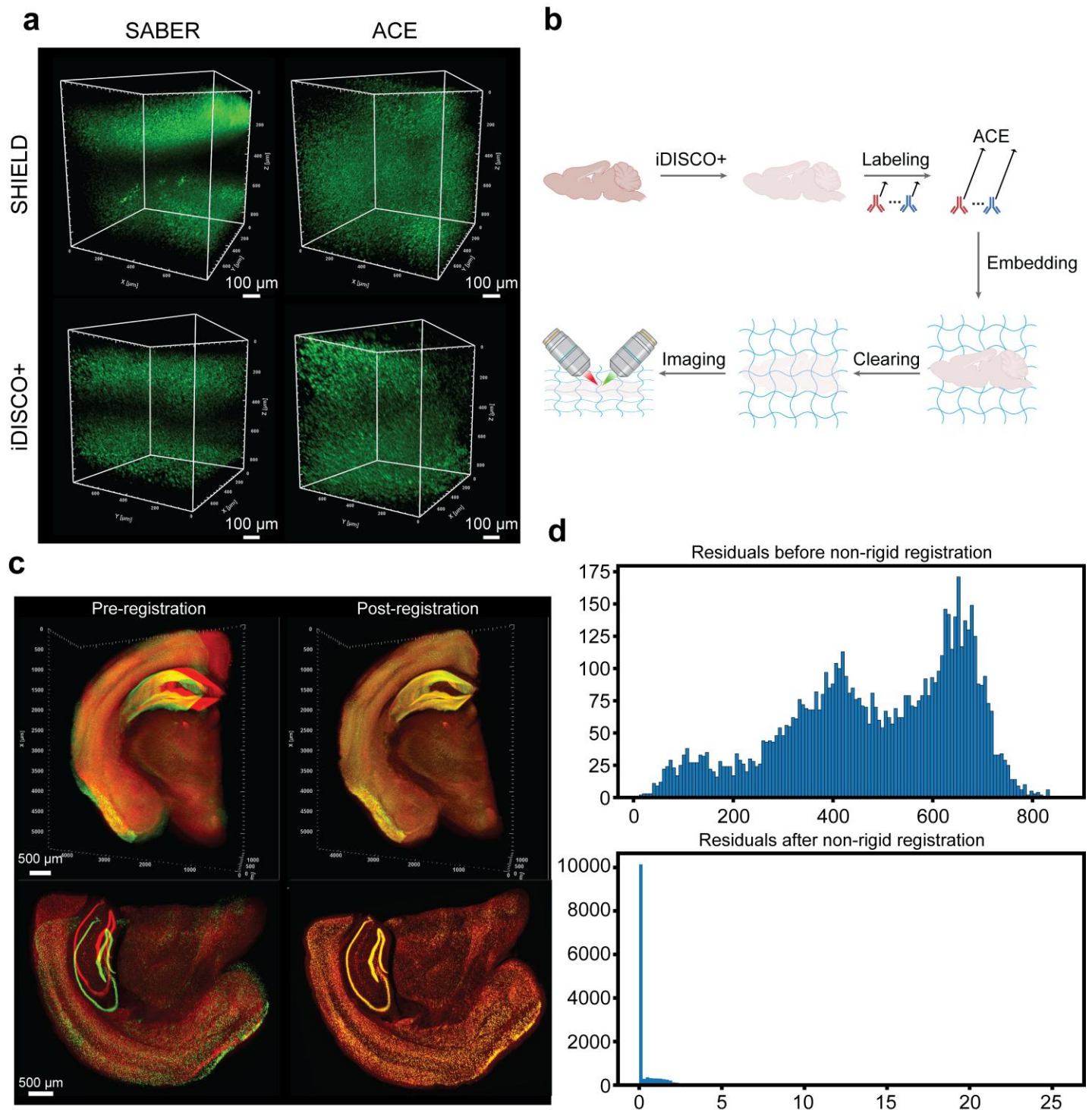

**Figure S19. Thick-tissue preparation and registration of imaging rounds.** (a) SABER vs ACE in 1-mm mouse brain sections using different tissue processing methods. (b) Schematic for ACE amplification and imaging of 200 μm sagittal section of mouse brain. (c) Before and after alignment for different rounds of imaging using NeuN as blobs for point cloud registration in a 1 mm tissue section. (d) Distribution of registration residuals between matched cell landmarks after initial coarse registration and 40 rounds of non-rigid registration in a 1 mm tissue section. Each residual represents the Euclidean distance between one transformed fixed-image landmark and its corresponding moving-image landmark. A total of 6,837 matched landmarks were analyzed. The initial and final median residuals were 500.90 μm and 3.99 μm, respectively. This point-cloud matching workflow was applied to both 200 μm and 1 mm tissue co-registration.

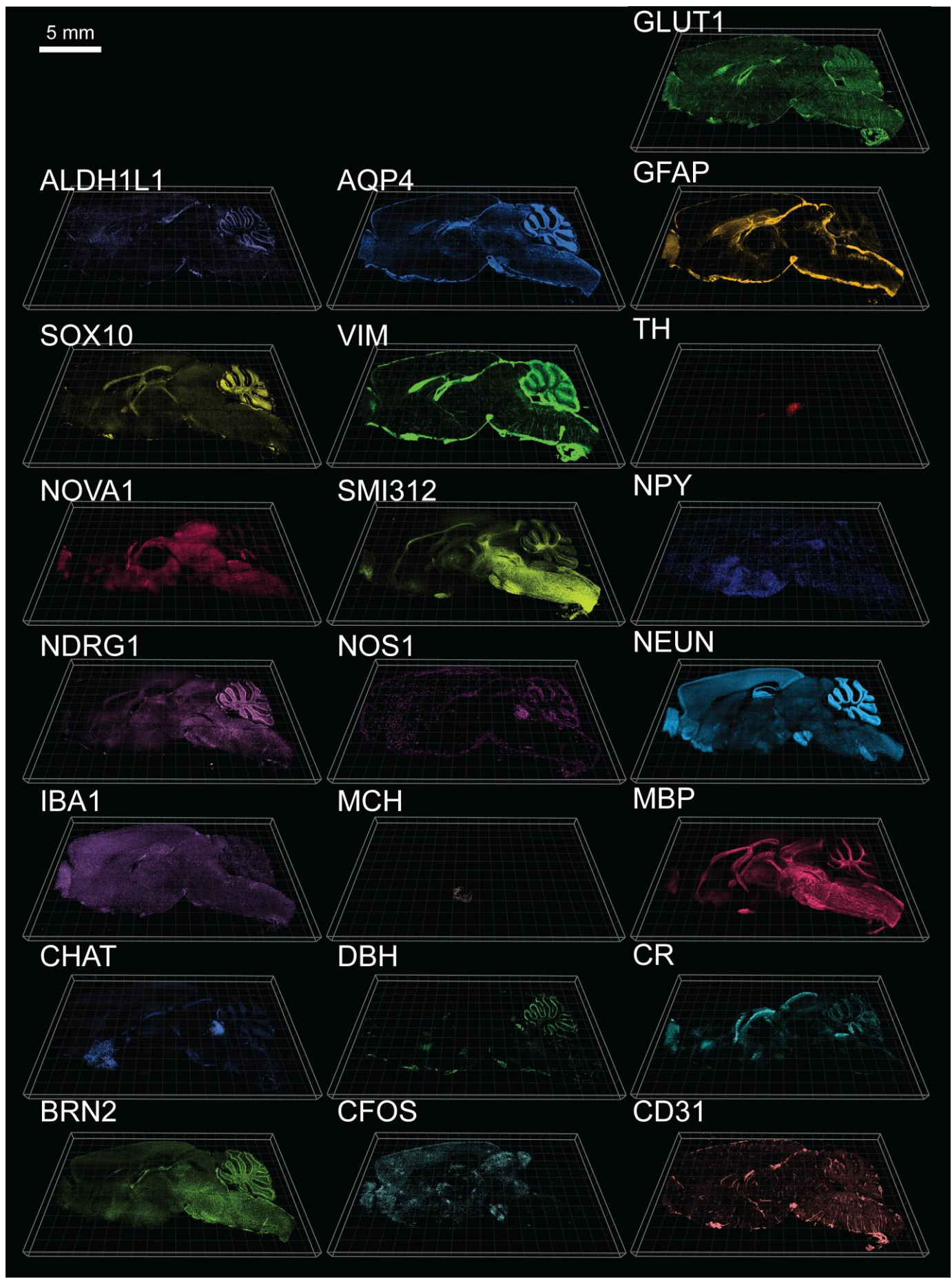

Figure S20. Zoomed-out views of 22 targets in the 200 μm sagittal section.

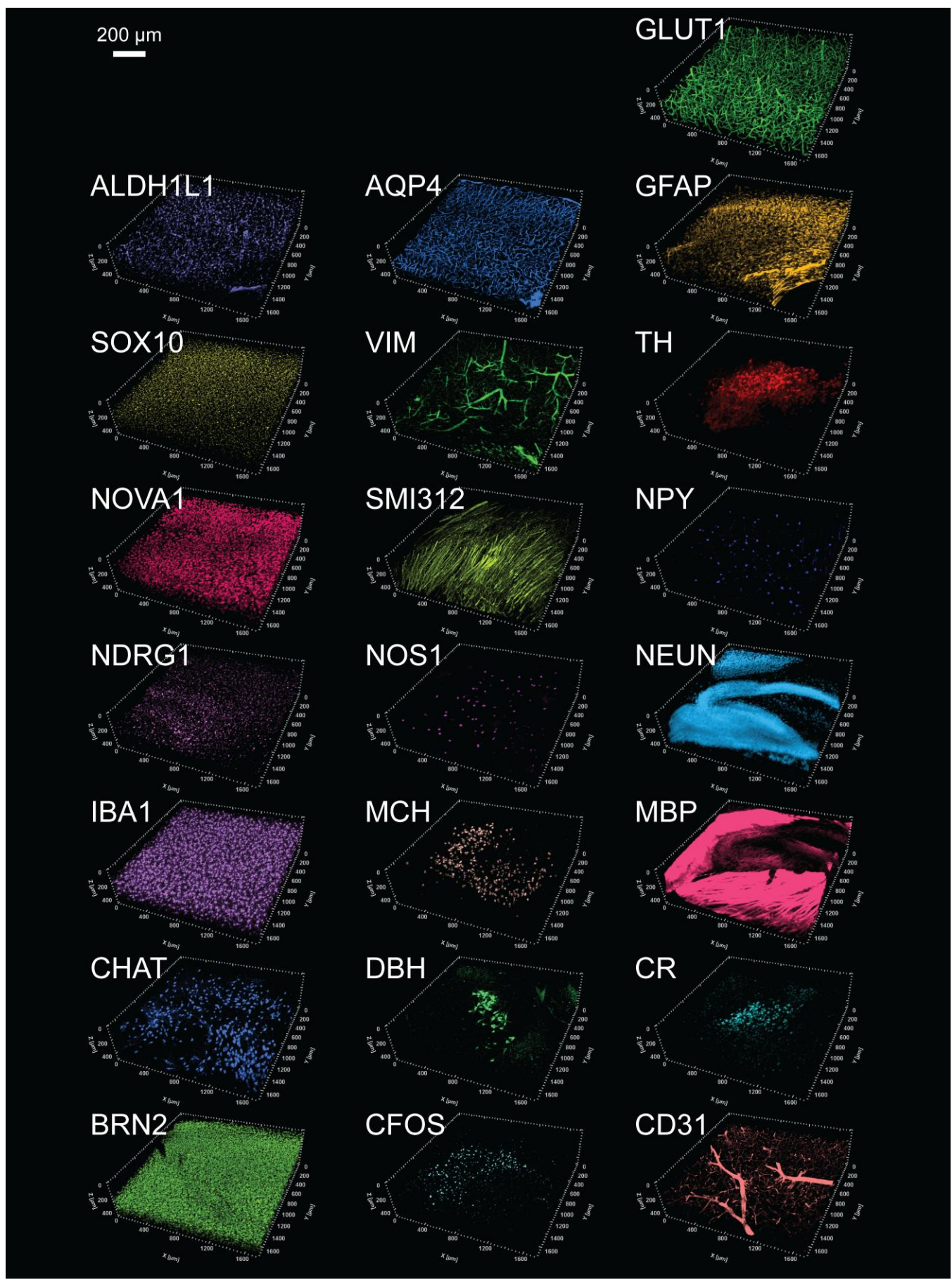

**Figure S21. Zoomed-in views of 22 targets in the 200 µm sagittal section.** Zoomed-in view of the 22 targets in the 200 µm sagittal section, with representative regions shown for each marker (regions vary).

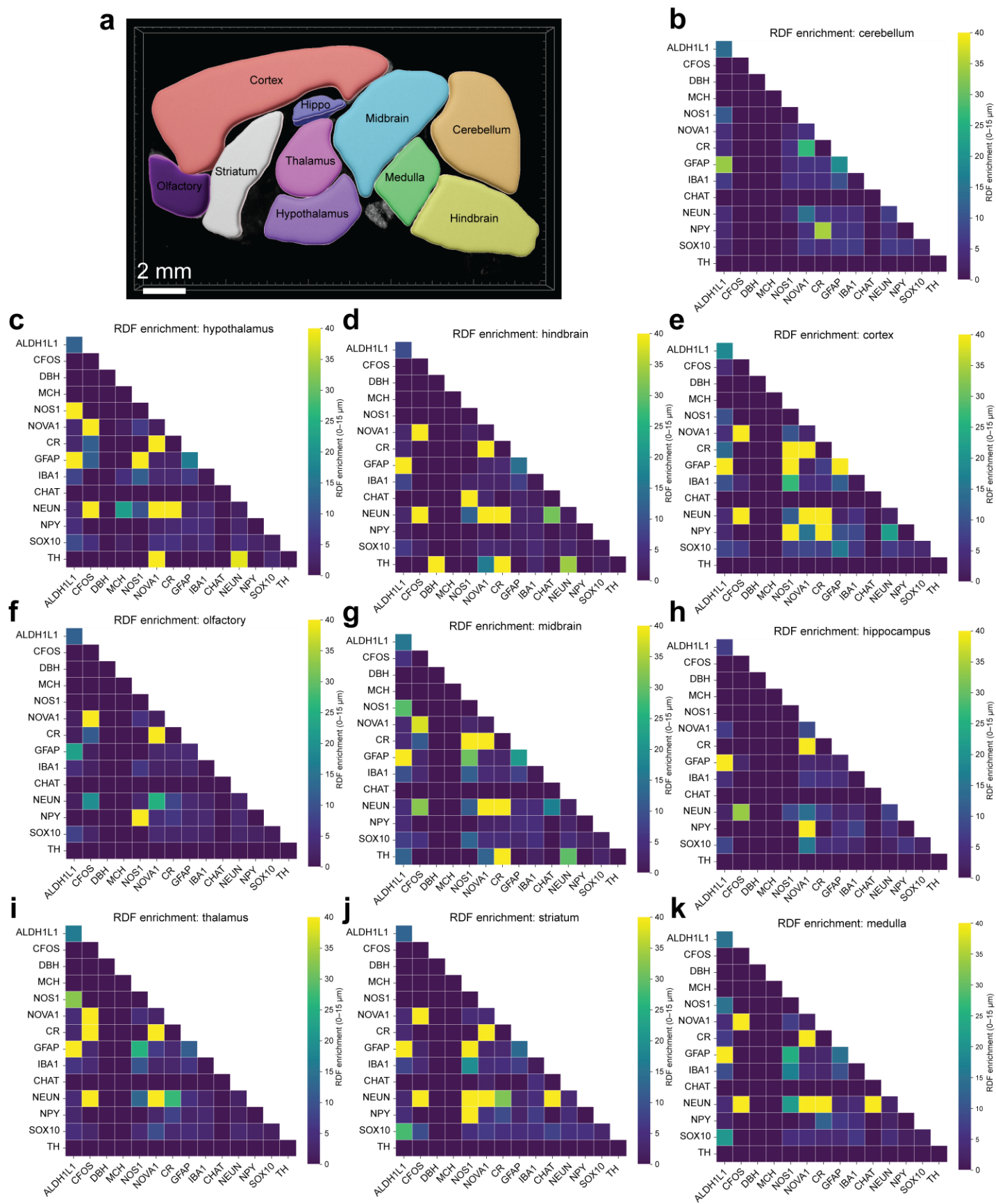

**Figure S22. Subregional annotation and cell neighborhoods in the 200 μm sagittal section.** (a) The subregional annotation of the 200 μm sagittal section. (b-k) Cell neighborhood analysis of the 14 cell-type markers in different subregions of the 200 μm sagittal section. $n = 2,542,660$ cells.

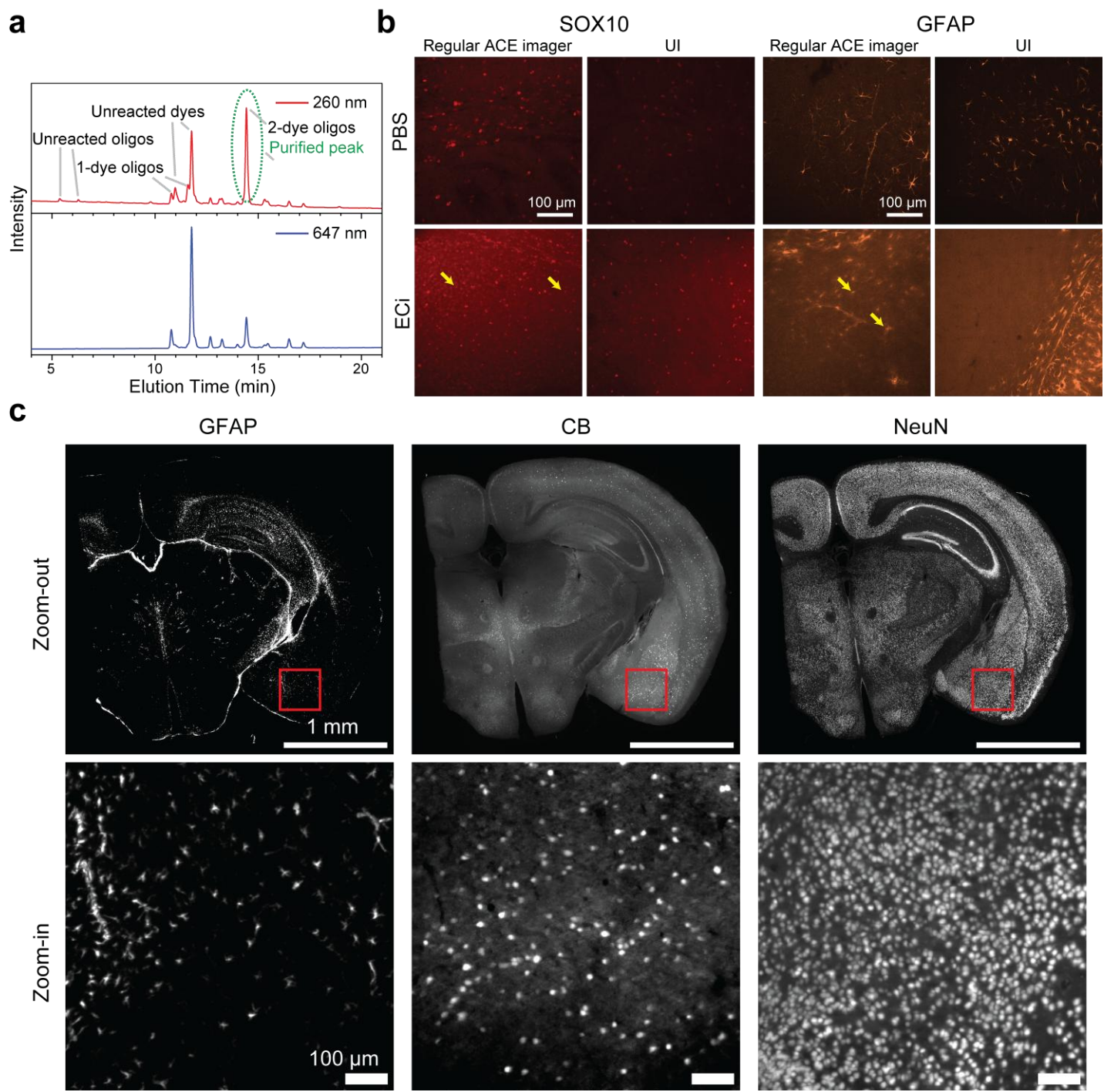

**Figure S23. Universal imagers and sequential imaging in refractive-index-matched tissue.** (a) HPLC was used to purify the DNA oligos conjugated with two dyes. (b) Comparison of regular ACE imagers and universal imagers (UI) for SOX10 and GFAP detection in 1-mm iDISCO+ mouse brain tissue before and after ECi refractive-index matching. Conventional ACE imagers produced target-specific signal in PBS but showed reduced compatibility after refractive index matching. For SOX10, the regular ACE imager showed increased nuclear background after ECi refractive-index matching (yellow arrows), whereas the universal imager maintained cleaner SOX10 staining. For GFAP, the regular ACE imager lost the expected astrocytic morphology after ECi processing, including fine protruding endfeet (yellow arrows), whereas the universal imager preserved GFAP-positive astrocyte processes. (c) Three consecutive rounds of targets imaged in the same fluorescence channel using the stripping-oligo strategy. Shown are maximum-intensity projections of 10 z-slices from the 1-mm section, with no residual signal detected between rounds.

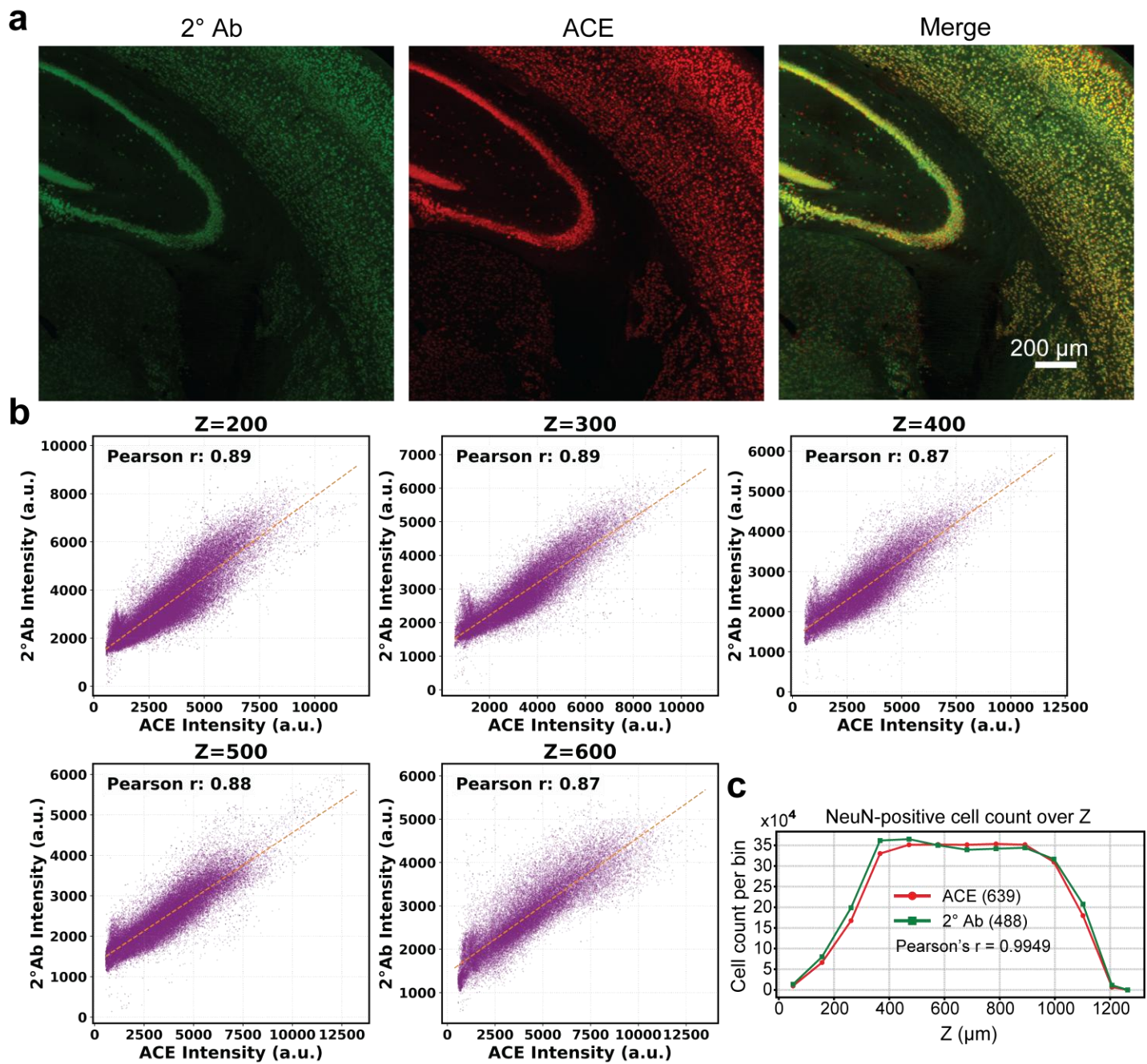

**Figure S24. Volumetric comparison of ACE and secondary-antibody NeuN signals.** (a) Spatial localization and merged-channel views of NeuN staining in a 1 mm mouse brain tissue sub-volume centered at $z = 300$ slices (out of 725 slices). Images show maximum-intensity projections (MIPs) over seven slices. (b) Scatter plots comparing pixel-neighborhood mean intensities of NeuN staining between the ACE channel (639 nm, x-axis) and the non-rigidly registered secondary antibody channel (488 nm, y-axis) at $Z = 200$, $Z = 300$, $Z = 400$, $Z = 500$, and $Z = 600$ slices. Intensities were sampled around high-confidence 3D volume-filtered landmarks within a 21-slice MIP window using a dilated neighborhood radius ($r = 3$ pixels). Dashed orange lines indicate ordinary least-squares linear fits. Inset values indicate the calculated Pearson correlation coefficients ($r$), showing stable, high-fidelity signal correlation ($r = 0.87$ to $0.89$) across the five depth blocks, indicating that the ACE intensity profiles accurately reflect target antibody distributions. (c) NeuN-positive cell counts per 60-slice $z$-bin ($105 \mu\text{m}$) plotted against tissue depth for ACE detection (639-nm channel; red circles) and secondary-antibody detection (488-nm channel; green squares) in the same 1-mm-thick tissue sample. Cell-count profiles were strongly correlated across depth bins (Pearson's $r = 0.9949$, $P < 0.0001$, two-tailed test).

**Figure S25. Comparison of ACE, HCR, and immuno-SABER for NeuN in 1 mm tissue.**

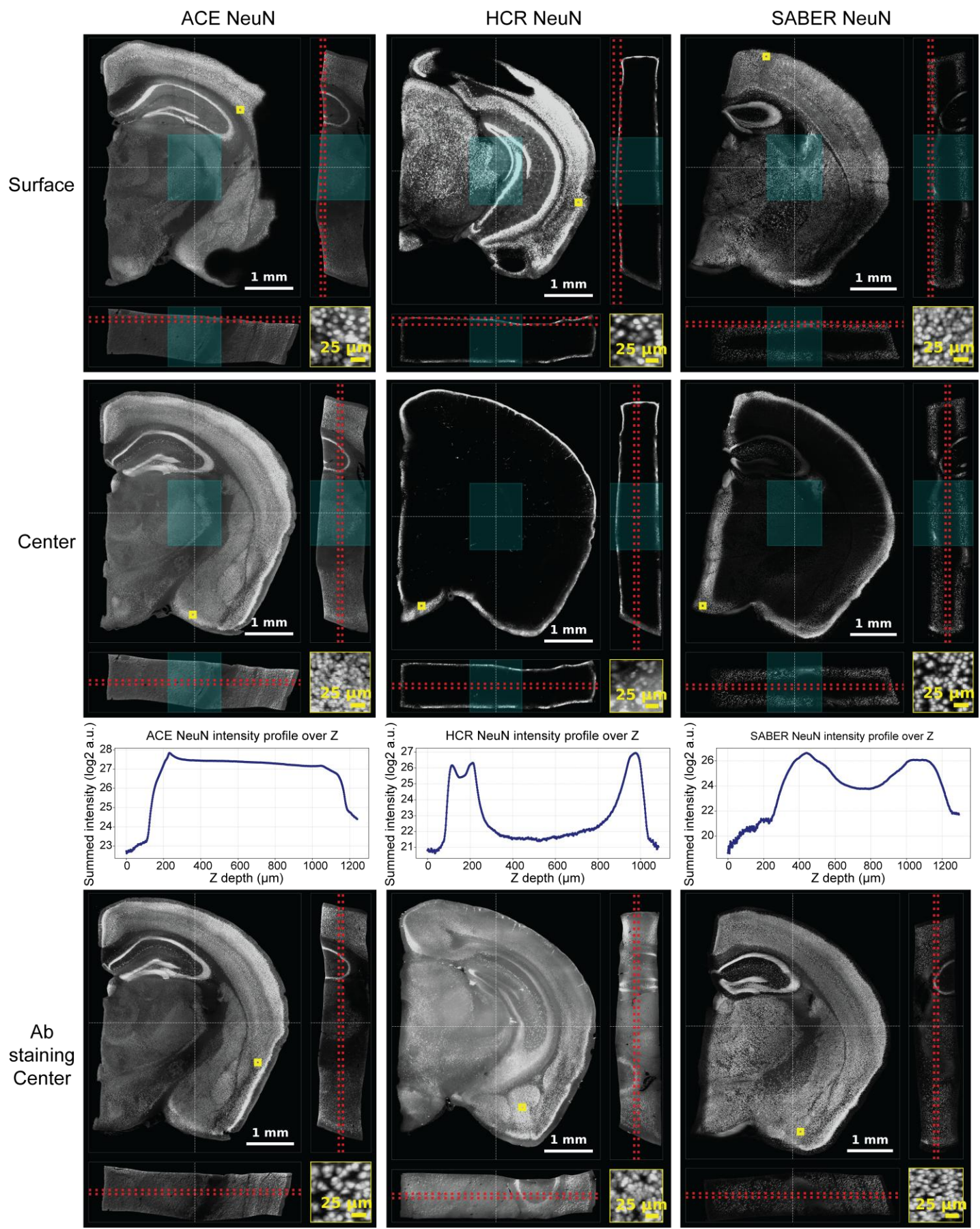

**Figure S26. Comparison of ACE, HCR, and immuno-SABER for CD31 in 1 mm tissue.**

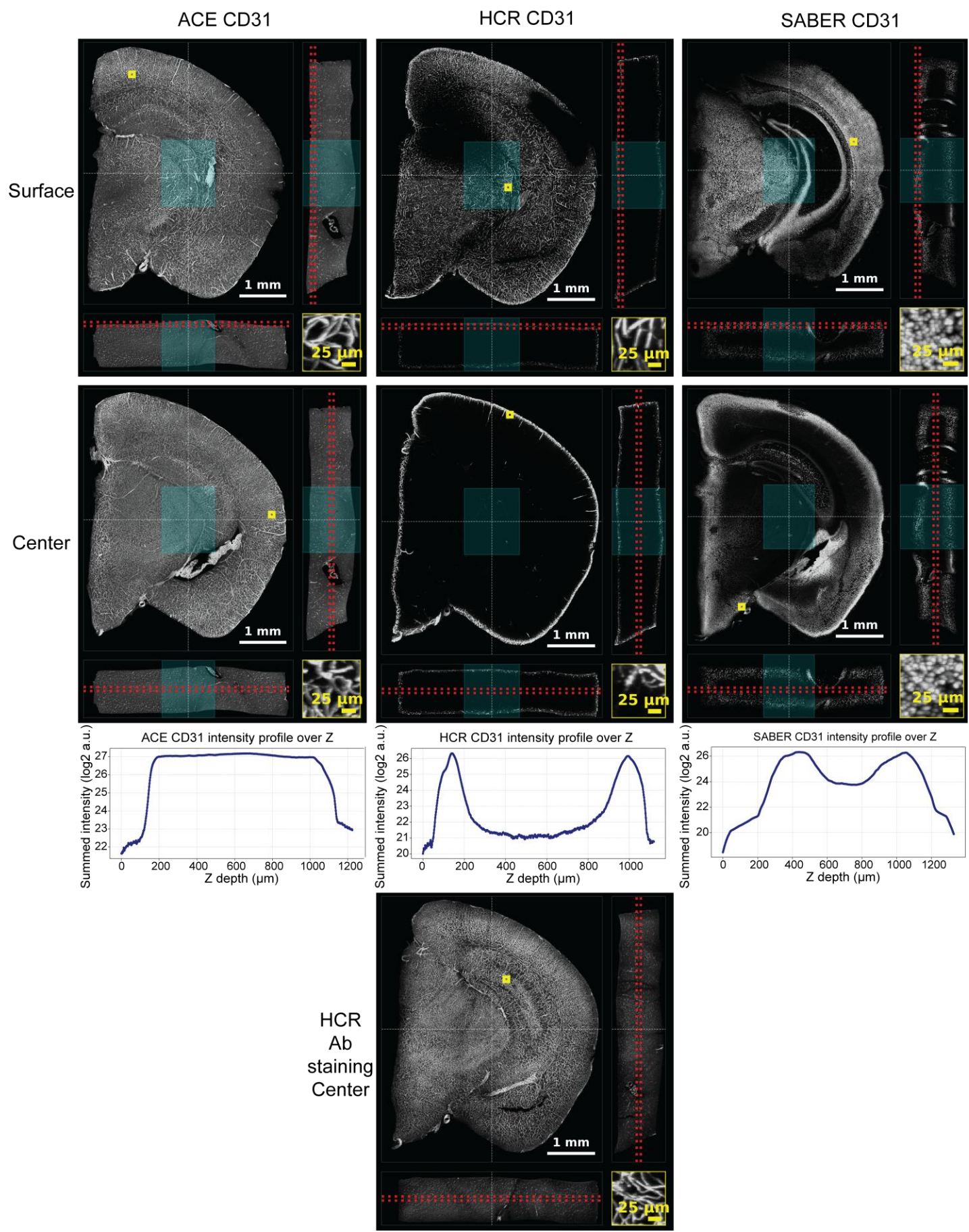

**Figure S27. Comparison of ACE, HCR, and immuno-SABER for GFAP in 1 mm tissue.**

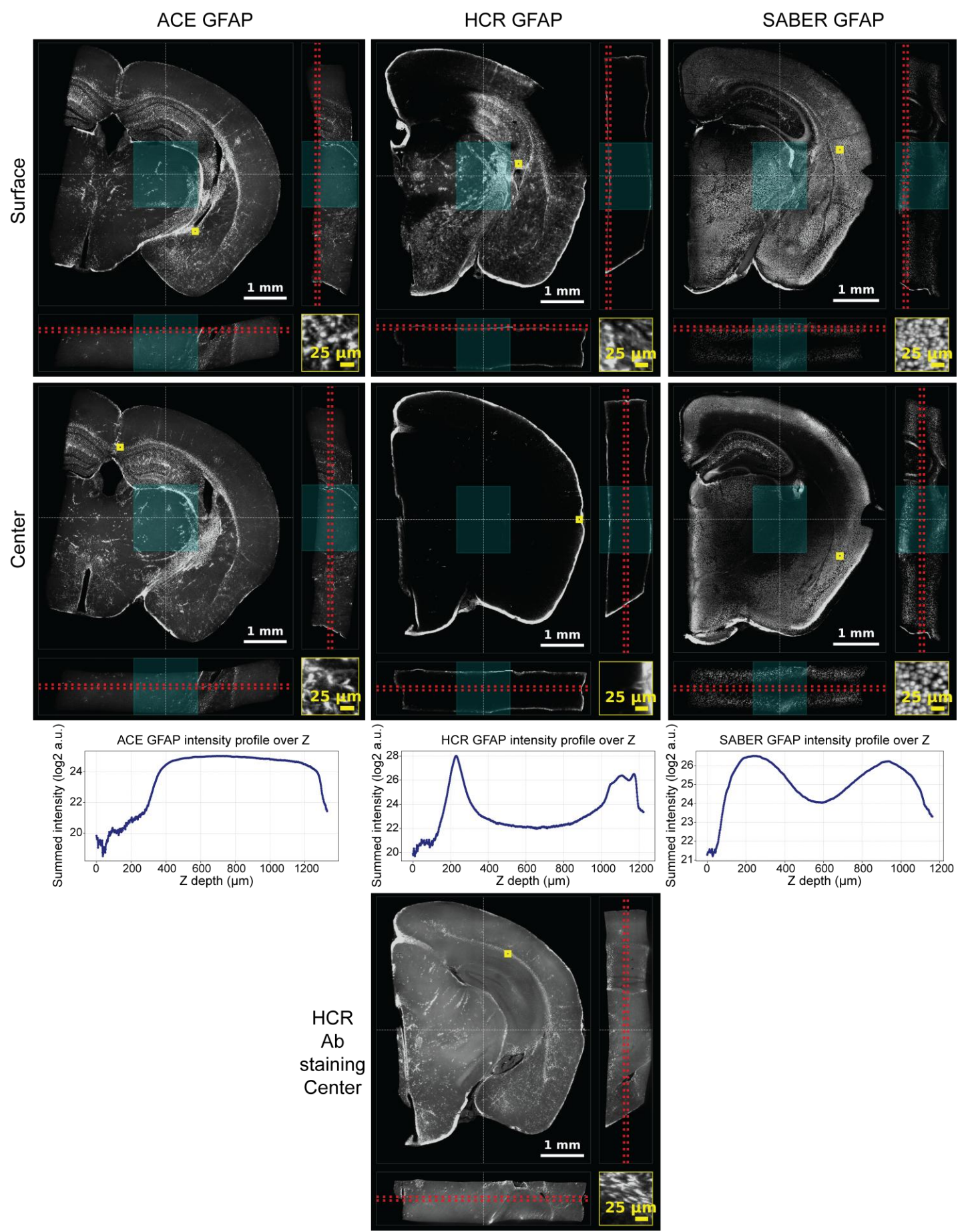

**Figure S28. Comparison of ACE, HCR, and immuno-SABER for OLIG2 in 1 mm tissue.**

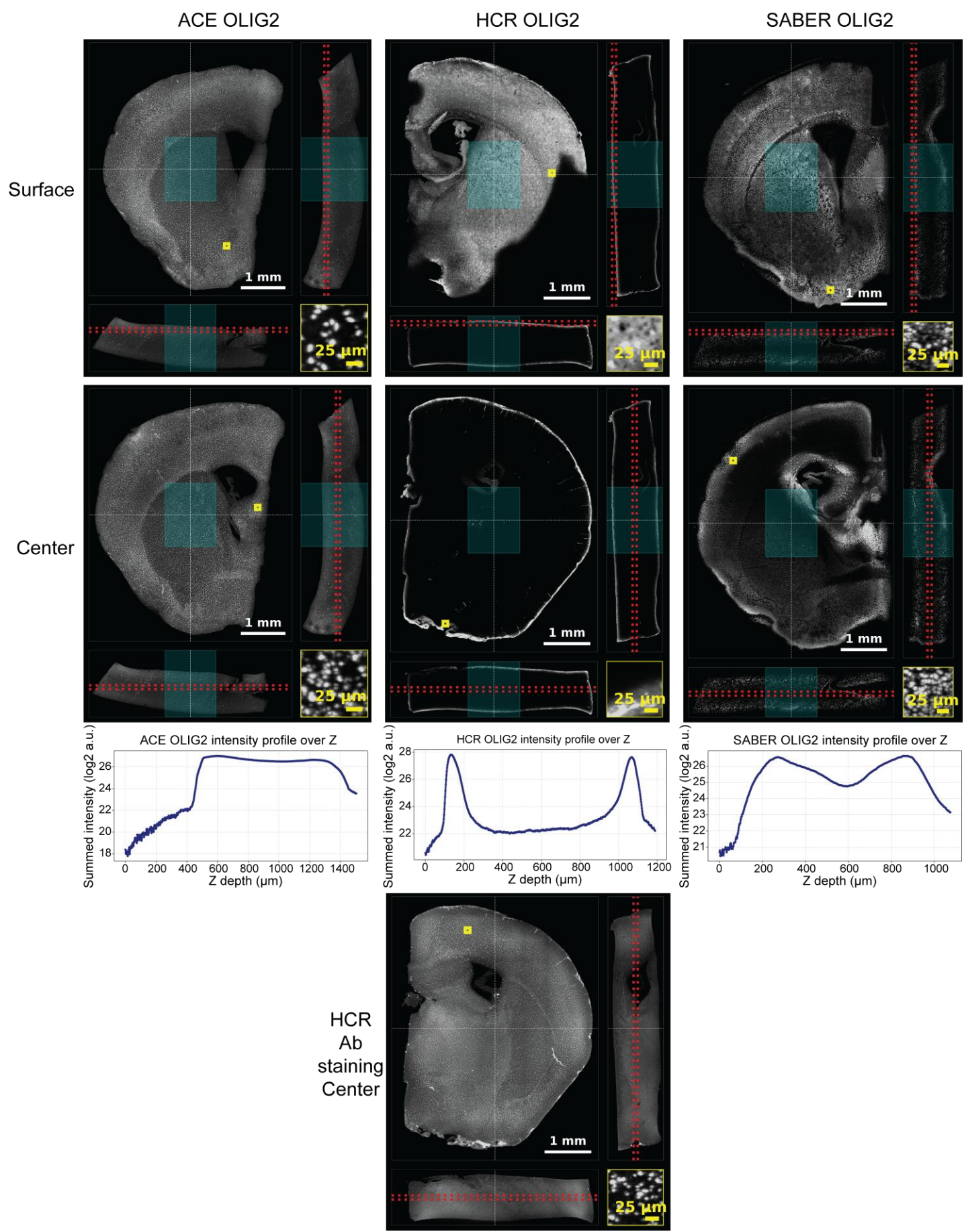

**Figures S25-S28. Shared legend for ACE, HCR, and immuno-SABER comparisons.** Comparison of ACE, HCR, and immuno-SABER for four representative protein markers spanning different expression levels and subcellular compartments in 1 mm mouse brain sections: **Figure S25**, NeuN, an abundant nuclear neuronal marker; **Figure S26**, CD31, an endothelial membrane marker; **Figure S27**, GFAP, a cytoplasmic astrocyte marker; and **Figure S28**, OLIG2, a relatively lower-expression nuclear oligodendrocyte-lineage marker. Images show signal near the tissue surface and center. Red dashed lines in the x-y and y-z views indicate the regions used to generate the corresponding maximum-intensity projections. For HCR, we used a commercial HCR protein imaging kit (Molecular Instruments, HCR™ Gold 2° IF). Fluorophore-labeled tertiary-antibody staining was used to assess antibody-labeling distribution before amplification. ACE produced marker-specific signal with relatively uniform intensity across depth, whereas other methods showed surface-bright, center-dim signal despite relatively uniform antibody staining. Line plots show summed signal intensity along z-depth plotted as log<sub>2</sub> arbitrary units. The area of the center 25% of both lateral dimensions (X and Y), indicated by the cyan square in the picture, was used to evaluate the intensity variation of tissue center over Z. In these figures, display contrast was adjusted independently for each method or condition to visualize staining patterns.

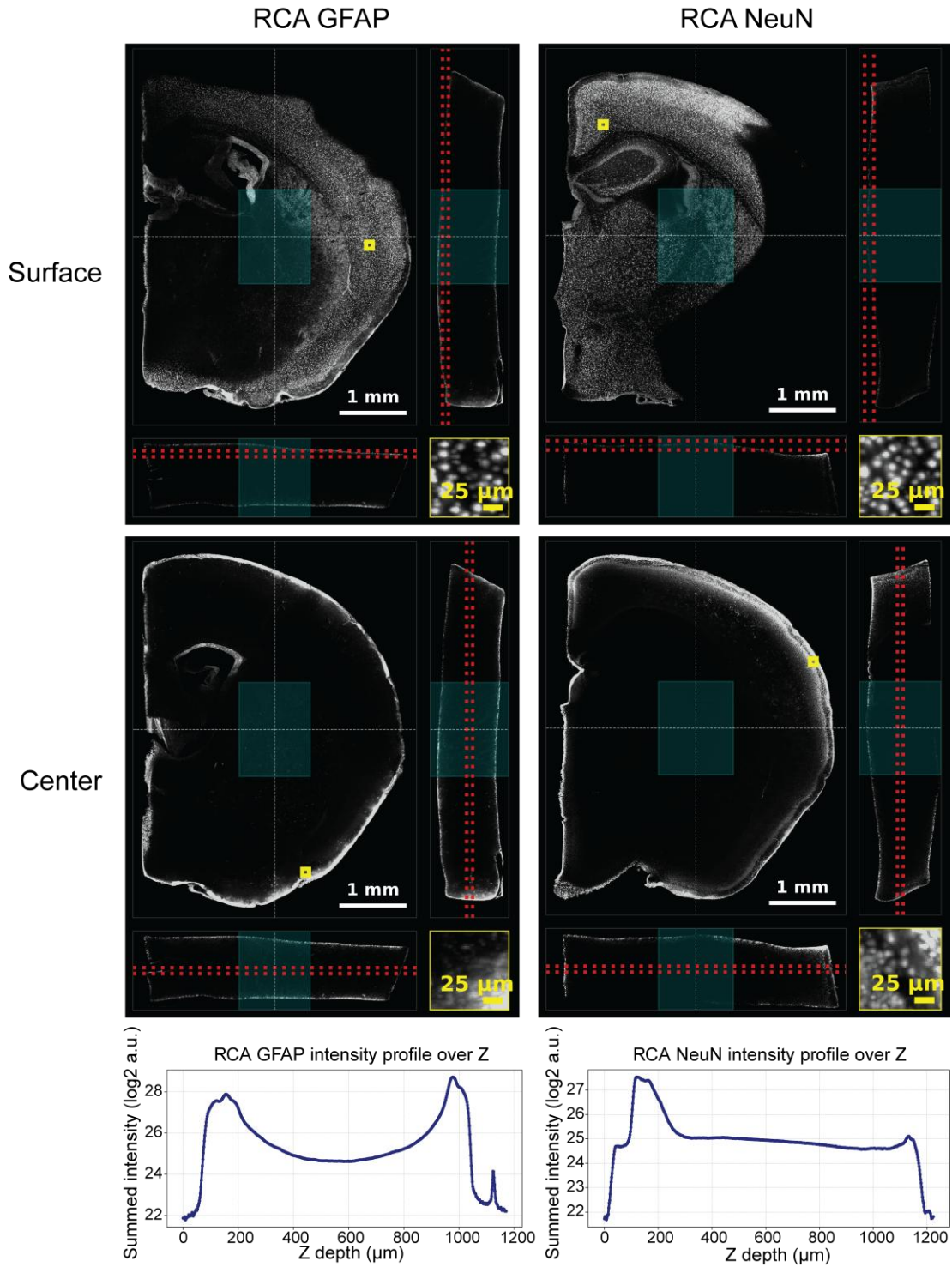

**Figure S29. RCA-based protein detection for GFAP and NeuN in 1 mm tissue.** RCA-based protein detection for GFAP and NeuN in 1-mm tissue. RCA produced nonspecific nuclear background and weak center signal under the tested conditions. Red dashed lines in the x-y and y-z views indicate the regions used to generate the corresponding maximum-intensity projections. Line plots show summed signal intensity along z-depth plotted as log2 arbitrary units. The area of the center 25% of both lateral dimensions (X and Y), indicated by the cyan square in the picture, was used to evaluate the intensity variation of tissue center over Z. In this figure, display contrast was adjusted independently for each condition to visualize staining patterns.

**Figure S30. Effects of preincubation and ssEXT concentration on signal uniformity.** (a) Fluorescent secondary-antibody staining of NeuN in 1-mm-thick tissue. (b) ACE NeuN imaging performed without preincubation of the reaction mixture in the tissue before thermal cycling. (c) ACE NeuN imaging performed with preincubation using ssEXT at 1 μM (undiluted), 100 nM (10× dilution), or 10 nM (100× dilution). Tissue-surface and central regions are shown as orthogonal views and magnified insets for each condition. Red dashed lines in the x-y and y-z views indicate the regions used to generate the corresponding maximum-intensity projections. The no-preincubation condition produced a surface-bright, center-dim signal profile, whereas the ssEXT dilution conditions showed high signal uniformity across the z-depth. (d) Depth-normalized mean intensity profile for the secondary-antibody staining shown in Fig. S30a. Mean intensity at each z-position was normalized to the mean intensity across all z-positions. The depth CV of the binned mean profile was 0.154. (e) Depth-normalized mean intensity profile for ACE performed without preincubation, corresponding to Fig. S30b, with depth CV = 0.309. (f) Depth-normalized mean intensity profiles for the ssEXT titration series in Fig. S30c and Fig. 7f. Depth profiles remained uniform with depth CVs of 0.105, 0.118 and 0.073 for 1 μM (regular ACE), 100 nM (10× dilution), and 10 nM ssEXT (100× dilution), respectively. In this figure, display contrast was adjusted independently for each method or condition to visualize staining patterns.

Figure S31. Zoomed-out views of 16 targets in the 1 mm coronal tissue block.

**Figure S32. Zoomed-in views of 16 targets in the 1 mm coronal tissue block.** Representative regions are shown for each marker (regions vary).

**Figure S33. Tissue boundaries and depth-resolved intensity profiles in the 1 mm block.** (a) Cross-sectional tissue area profile (black curve) computed across all Z-slices based on voxels exceeding intensity threshold 700. Vertical dashed blue lines indicate the established 10% cutoff threshold boundaries (start slice 73, 127.75 $\mu\text{m}$;

end slice 671, 1174.25 $\mu\text{m}$), corresponding to 10% of the maximum tissue area (7,985,645 voxels). Faint dotted horizontal line shows the absolute area cutoff value. **(b)** Representative binarized tissue mask visual slices at the boundary cutoff points: beginning start slice (73) and ending stop slice (671), mapped with a viridis colormap. Scale bar represents 1 mm. **(c)** Individual target depth-intensity distribution profiles and sampling variations. Multi-panel grid showing individual depth profiles for 14 spot-detection targets and 2 vessel-mask targets (CD31 and GLUT1). The primary left axis (blue line) tracks the relative mean intensity ($\pm 1$ STD shaded boundary, light blue) normalized against the target's baseline average across 20 $\mu\text{m}$ Z-bins. The secondary right axis (orange dashed line) maps the relative standard error of the mean (SEM), establishing high localized sampling precision within the active tissue window.

**Figure S34. Subregional annotation and cell neighborhoods in the 1 mm coronal block.** (a) The subregional annotation of the 1 mm coronal tissue block. (b) Cell neighborhood analysis of the 10 cell-type markers in different subregions of the 1 mm coronal tissue block. $n = 3,464,350$ cells.

**Figure S35. Nine-plex ACE protein imaging in human tonsil FFPE tissue.** (a) Whole-section views of 9-plex ACE protein imaging performed on a human tonsil FFPE tissue section. Individual channels show CD4, Ki67, CD11b, CK6, CD63, HLA-DR, CD31, CD19 and CD45. Scale bar, 5 mm. (b) Magnified views of the same markers showing marker-specific staining patterns and preserved tissue morphology. Scale bar, 1 mm. Tissue section thickness, 5 μm.

**ACE orthogonal sequence design**

The initial set of 50 ACE sequences was modified from our previous work <sup>1,2</sup>. To expand the number of sequences, we used Seqwalk <sup>14</sup> to identify sequences that are orthogonal to the initial set and generated 67 initiator-ssEXT pairs for 9 mers. Each ssEXT has two repeats of the reverse complement of the 9-mer initiators separated by a *T* base. Inverted *T* was used on the 3' end to prevent the extension from ssEXTs. All sequences were synthesized by IDT. ssEXTs were purified by HPLC. All DNA strands were dissolved in RNase-free water. See **Table S1** for specific sequences.

**RNA probe design**

RNA probes were designed using PaintSHOP with the hg38 newBalance probe set for human targets and mm39 newBalance for mouse targets <sup>15</sup>. Parameters were set to Max Off-Target Score= 200, Max K-mer Count =4, and Min Prob Value=0. The same ACE initiator was attached to the 3' end of all the probes targeting the same gene with *ATTT* as the spacer between the RNA-binding sequences and ACE initiators. After design, the sequences were filtered to retain those with melting temperature > 65 °C in 1× ThermoPol buffer. The sequences were also filtered based on off-target score and secondary structure probability. 17 to 65 probes were used for each gene. See **Table S2-S4** for specific sequences.

**In-tube ACE reaction**

9 nt initiators were synthesized by IDT. For in-tube ACE reactions, the reaction mixture contained 1× ThermoPol (NEB B9004S), 300 μM each of dATP, dTTP, and dCTP (NEB N0446S), 10 nM each initiator, 10 μM each ssEXT (IDT), 2 U/μL Bst large fragment DNA polymerase (NEB M0275M). For each initiator, 20 μL of ACE reaction mixture was added to a PCR tube. The tubes were placed on a thermal cycler. Amplification was achieved by 0-300 cycles of 22 °C 35 s, 58 °C 25 s. The distribution of the amplification was visualized by running the samples on denaturing urea-PAGE gels. For the one-ssEXT versus 50-ssEXTs test, each ssEXT was used at a final concentration of 500 nM.

**Crosstalk validation by next-generation sequencing**

To test the crosstalk among initiators with ssEXT, we first designed all the initiators with the same ligation adaptor (*CCTCTCTCTCCCCTCTCTCTCTTT*) on the 5' end. A 100 μL reaction containing 1× ThermoPol, 300 μM each of dATP, dTTP, and dCTP, 2 U/μL Bst large fragment DNA polymerase, all initiators with ligation adapter (1 nM each), and all ssEXTs (200 nM each) was mixed and split equally between two PCR tubes. The tubes were placed on a thermal cycler and subjected to 120 cycles of ACE amplification of 22 °C 35 s, 58 °C 25 s. After amplification, the reverse ligation primer (*GAGAGAGAGGGGAGAGAGAGG*) was added to the sample to a final concentration of 100 nM. The sample was subsequently purified using 1× volume Ampure XP beads (Beckman, a63881) and eluted in 20 μL ddH<sub>2</sub>O. A 23 μL ligation reaction with 10 μL extended product, 1 μL of 20 μM Illumina ligation adapter, 10 μL of 2× quick ligation buffer (NEB M2200L), 1 μL of Quick ligase (NEB M2200L), 1 μL of T4PNK (NEB M0236L) was used to ligate Illumina adapters on the samples. After ligation, the samples were purified using 1× volume Ampure XP beads and eluted into 20 μL of ddH<sub>2</sub>O. The ligated product was further PCR amplified using Illumina P5P7 primers with the following reaction: 10 μL of ligated product, 4 μL of 5× Q5 reaction buffer (NEB M0493L), 0.4 μL of dNTPs (10 mM each), 1 μL of Q5 high fidelity hot start DNA polymerase (NEB M0493L), 0.3 μL of Illumina P5 primer (10 μM), 0.3 μL of Illumina P7 primer (10 μM), and 4 μL of ddH<sub>2</sub>O. PCR was performed with the recommended protocol with 55 °C annealing temperature for 7 cycles. Amplified product was purified using 1× volume Ampure XP beads into 20 μL ddH<sub>2</sub>O. The amplified library was measured using Qubit dsDNA High Sensitivity kit (ThermoFisher Scientific Q33230). Library was diluted according to Illumina protocol. Sequencing was performed on an Illumina MiSeq using 300 cycles kit (Illumina MS-102-2002) or a 500 cycles kit (Illumina MS-102-2003). After sequencing, the generated FASTQ file was split based on initiators. For each initiator FASTQ file, extension specificity was calculated based on the frequency of each ACE sequence to the correct sequence for each extension event.

**DNA-antibody conjugation**

Antibodies were ordered from vendors in carrier-free formulation. For antibodies that were only available in unpurified whole serum form or formulated with stabilizers that interfere with conjugation, purification with protein A/G beads was applied to recover antibodies. Next, 50 μg of antibody was rinsed by two sequential centrifugation steps (6 min at 8,000 × g) in an Amicon Ultra filter (50 kDa) with 500 μL 1× PBS at 4 °C. Crosslinker solution was prepared by 1:10 dilution of NHS-Sulfo-DBCO stock (Vector Laboratories CCT-A124-25, 10 mg/mL in anhydrous DMSO or DMF, stored at -20 °C). Then, 1.8 μL of crosslinker stock solution and 3.6 μL of azide-modified DNA (1 mM stock concentration) were added into the Amicon filter tube with antibody. These amounts corresponded to an input DNA-to-antibody molar ratio of approximately 11:1. The solution was gently tapped to mix and stored at 4 °C overnight. The next day, the excess crosslinker and DNA were removed from the antibody solution by five sequential centrifugation steps (6 min at $8,000 \times g$) in the same Amicon filter tube. The final conjugated antibody was collected by inverting the Amicon filter and spinning 3 min at $3,000 \times g$. The successful conjugation was confirmed by SDS-PAGE, and the concentration of the product was measured using Qubit Protein Assay. The specificity of the conjugated antibody was validated via indirect immunofluorescence. To facilitate long-term storage, the DNA-conjugated antibody was supplemented with 10 mM EDTA, 0.25% BSA, 0.2% sodium azide, 0.2% Triton X-100, and 50% v/v glycerol. The conjugated antibody was preserved at $-20^\circ\text{C}$ until use.

##### **Gel electrophoresis**

**SDS-PAGE Antibody Gel.** Successful antibody conjugation was confirmed by gel electrophoresis on NuPAGE 4-12% Bis-Tris gels (Invitrogen #NP0321BOX). Samples were prepared by diluting 1 $\mu\text{L}$ of conjugated antibody into 9 $\mu\text{L}$ of $1\times$ NuPAGE LDS Sample Buffer (Invitrogen #NP0007) and then heated at $80^\circ\text{C}$ for 10 min. Meanwhile, the gels were loaded into the electrophoresis chamber and filled with $1\times$ MOPS SDS Running Buffer (Novex #NP0001). After heating, the samples were loaded into the gels along with 5 $\mu\text{L}$ of PageRuler Prestained Protein Ladder (Thermo #26616). Gels were run at 180 V for 50 min, after which they were removed from the chamber, stained with InstantBlue Coomassie Protein Stain (Abcam #ab119211) for 15 min, rinsed with tap water, and imaged on a Sapphire Gel Imager.

**PAGE DNA Gel.** The lengths of ex-situ generated concatemers were visualized by gel electrophoresis on 6% TBE Urea gels (Novex #EC6865BOX). Samples were prepared by diluting 10 $\mu\text{L}$ of extended concatemers with 10 $\mu\text{L}$ of denaturing buffer (Invitrogen #AM8547). The 50 bp DNA ladder was first diluted 1:10 in $\text{DIH}_2\text{O}$ and then 1:2 in denaturing buffer. The samples were then heated at $90^\circ\text{C}$ for 1 min and allowed to come to room temperature. Meanwhile, the gels were loaded into a heated gel electrophoresis flow chamber, filled with $1\times$ TBE Buffer (Invitrogen #15581044), and pre-run at $55^\circ\text{C}$ and 200 V for 10 min. After pre-running, the wells were cleared of stagnant urea by washing with a pipette dropper and then loaded with the samples and DNA ladder. Gels were run at $55^\circ\text{C}$ and 200 V for 30 min, after which they were removed from the chamber and stained with 1:10,000 Sybr-Gold (Invitrogen #S11494) in $1\times$ TBE for 15 min. After staining, gels were washed 2 x 5 min with $\text{H}_2\text{O}$ and imaged on an Azure Sapphire Biomolecular Imager.

##### **Cell culture**

HeLa cells and Py2T cells were cultured in a cell culture medium composed of DMEM High Glucose (Gibco #11965092) supplemented with 10% Fetal Bovine Serum (Gibco #16000044) and 1% Penicillin-Streptomycin (Gibco #15070063). First, 1 mL of frozen stock were thawed and seeded into a T75 cell culture flask in 12 mL of cell culture medium and incubated at $37^\circ\text{C}$. After cells were grown to $\sim 90\%$ confluency, the cells were washed with $1\times$ DPBS and incubated for 10 min with 3 mL 0.05% Trypsin-EDTA (Gibco #25300054) at $37^\circ\text{C}$. After incubation, the trypsin was neutralized with 7 mL of cell culture medium and all 10 mL of suspended cells were transferred to a 15 mL conical tube. The cells were centrifuged at $1,200 \times g$ for 5 min, after which the supernatant was removed, and the cell pellet was resuspended in 10 mL of cell culture medium. To continue culture of the cell line, 100 $\mu\text{L}$ of cell suspension was added to a new T75 flask with 12 mL of cell culture medium where it was incubated at $37^\circ\text{C}$ until $\sim 90\%$ confluency, or approximately 2-3 days.

##### **Cell sample preparation**

Cells were cultured in an ibidi 18-well glass-bottom chamber (ibidi 81817) to appropriate confluency. Cells were then washed twice with $1\times$ PBS to wash off the cell culture medium and fixed with 4% paraformaldehyde for 10 min. 0.1 M glycine was then used to quench the fixation. Cells were washed with 0.1% Triton and 4 U/mL RNase inhibitor (NEB, M0314L) in $1\times$ PBS three times.

For RNA imaging, cells were permeabilized with 0.5% Triton and 4 U/mL RNase inhibitor in $1\times$ PBS for 10 min then washed with 0.1% Tween 20 and 4 U/mL RNase inhibitor in $2\times$ SSC three times. RNA probes were diluted in $1\times$ FISH buffer containing 10% Dextran sulfate, $2\times$ SSC, 40% Formamide, and 0.1% Tween 20. The final concentration of the probe set targeting each gene was 200 nM. Samples were incubated at $60^\circ\text{C}$ for 3 min and incubated at $42^\circ\text{C}$ for at least 6 hours. Samples were quickly washed with 30% Formamide in $1\times$ PBS 2 times and incubated and washed with 30% Formamide in $1\times$ PBS for 3 min 5 times. Samples were then washed with 0.1% Triton and 4 U/mL RNase inhibitor in $1\times$ PBS three times. After washes, samples were then ready for ACE amplification.

For protein imaging, samples were blocked and permeabilized with 5% BSA, 4 mM EDTA, and 0.3% Triton in $1\times$ PBS for 1 hour. Antibodies were premixed at 5 $\mu\text{g}/\text{mL}$ in 3% BSA, 0.2% Triton X-100 with 4 mM EDTA, 0.02% dextran sulfate, and 500 nM antibody DNA blockers. The blocking buffer was aspirated before the antibody mix solution was added to the samples. Antibodies were incubated for at least 3 h at room temperature or $4^\circ\text{C}$ overnight. After antibody incubation, samples were incubated and washed with 0.1% Triton in 1× PBS five times for 3 min each. A post-fixation was subsequently performed using freshly prepared 5 mM BS(PEG)<sub>5</sub> for 30 min. 0.1 M glycine was then used to quench the fixation. After quenching, samples were washed with 0.1% Triton in 1× PBS three times. After washes, samples were ready for ACE amplification.

For protein and RNA imaging, cells were first stained with antibodies using the protein imaging protocol mentioned above. All buffers and solutions except for the post-fixation buffer were supplemented with RNase inhibitor with a final concentration of 4 U/mL. After post-fixation and washes, cells were further permeabilized with 0.5% Triton and 4 U/mL RNase inhibitor in 1× PBS for 10 min and washed with 0.1% Tween 20 in 2× SSC three times. RNA staining was performed using the RNA protocol mentioned above.

The ACE reaction mixture contained 1× ThermoPol (NEB B9004S), 300 μM each of dATP, dTTP, and dCTP (NEB N0446S), 1 μM each ssEXT (IDT), 2 U/μL Bst large fragment DNA polymerase (NEB M0275M or MCLab BPL-400). For each well of an 18-well ibidi chamber, 40 μL of ACE reaction mixture was added. The well was sealed with 60 μL mineral oil. The chamber was placed on a plate thermal cycler (such as Eppendorf nexus flat and ProFlex™ PCR System). Amplification was achieved by 120-300 cycles of 22 °C 35 s, 58 °C 25 s, with heated lid set to 60 °C. After ACE amplification, samples were washed with 50% Formamide five times for 3 min each. Imagers in 4% Dextran sulfate and 30% formamide in 1× PBS were added to the amplified sample for 1 hour. Cells were washed with 30% formamide five times for 3 min each, then stained with DAPI in 1× PBS for 5 min.

##### **FAM-dCTP test**

We cultured the HeLa cells in an 18-well ibidi chamber to reach ~70% confluency. To evaluate nonspecific extension of ACE reactions *in situ*, we stained the HeLa cells with 2 μg/mL initiator-conjugated anti-β-tubulin antibodies overnight at 4 °C. After post-fixation, ACE reaction was performed. We spiked in 30 μM FAM-labeled dCTP to label DNA generated by the ACE reaction. After reaction, samples were washed 4 times with 1× PBST and imaged.

##### **Loss-of-signal experiment**

HeLa cells were seeded at 30,000 cells per well in an Ibidi 18-well glass-bottom chamber (Ibidi, #81817) and cultured overnight. One well was used for the experiment. Cells were stained with Lamin A/C antibody. 100 cycles of ACE amplification were performed according to the protocol mentioned above. Samples were then washed three times for 5 min with 60% formamide in 1× PBS and incubated for 30–60 min at room temperature with 50 μL imaging solution containing 100 nM universal imager 647 adaptor and 120 nM Alexa Fluor 647 universal imager in 30% formamide and 4% dextran sulfate. Following three 5-min washes with 30% formamide in 1× PBS, cells were stained with DAPI in 1× PBS for 10 min, washed three times with 1× PBS, and imaged in oxygen-scavenging solution containing 5× protocatechuic acid, 2× Trolox, and 7.5 U/mL recombinant protocatechuate 3,4-dioxygenase in 1× PBS. After each imaging round, signal was removed by six 5-min washes with 60% formamide, followed by three washes with 1× PBS, before repeating the imager incubation and imaging cycle.

##### **GFP antibody dilution**

To fluorescently tag the tubulin, the cells were transfected with GFP-tubulin using the CellLight Reagent BacMam 2.0 system from ThermoFisher. In short, 5 μL of the BacMam 2.0 reagent was combined with 55 μL of culture medium in an ibidi 18-well cell culture plate when cells were at 50% confluence. These cells were then incubated at 37 °C for 16 hours. After incubation, they were fixed using 4% PFA for 20 min, then quenched for 10 min in a solution of 0.1 M glycine combined with 1× PBS. Subsequently, the HeLa cells were stained using a 1:1,000,000 dilution of the DNA-barcoded anti-GFP antibody (final concentration, 100 pg/mL) and subjected to ACE signal enhancement. The prepared sample was imaged using a Nikon Ti microscope integrated with the CSU-W1 spinning disk system.

##### **Flow cytometry**

All spins in these experiments were 500 × g for 1 min. HEK293T cells were transfected with GFP plasmid using Lipofectamine (ThermoFisher Scientific 18324012). Cells were trypsinized to a single-cell suspension the next day in a 1.5 mL tube, quickly spun, and washed with 1× PBS twice. Cells were then resuspended and fixed with 4% paraformaldehyde for 10 min, then spun and washed with 0.1 M glycine for 5 min. After quenching, cells were washed with 0.1% Triton in 1× PBS three times. Cells were then blocked and permeabilized with 5% BSA, 4 mM EDTA, and 0.3% Triton in 1× PBS for 1 hour. After spinning and aspirating the blocking solution, antibody mix containing 2 μg/mL anti-GFP antibody in 3% BSA, 0.2% Triton X-100 with 4 mM EDTA, 0.02% dextran sulfate, and 500 nM antibody DNA blockers was added. After antibody incubation, samples were spun and washed with 0.1% Triton in 1× PBS five times for

3 min each. Cells were split into two tubes. For tube 1, a secondary antibody mix containing 2 $\mu\text{g}/\text{mL}$ anti-rat Alexa-647 antibody in 3% BSA, 0.2% Triton X-100 was added for 2 hours and spun and washed with 0.1% Triton in 1× PBS five times for 3 min each. For tube 2, cells were post-fixed with freshly prepared 5 mM BS(PEG)<sub>5</sub> for 30 min. 0.1 M glycine was then used to quench the fixation. After quenching, post-fixed cells were washed with 0.1% Triton in 1× PBS three times. For each PCR tube, 100k post-fixed cells were added into 50 $\mu\text{L}$ of ACE mix containing 1× ThermoPol, 200 $\mu\text{M}$ each of dATP, dTTP, and dCTP, 1 $\mu\text{M}$ ssEXT, 2 U/$\mu\text{L}$ Bst large fragment DNA polymerase. Amplification was achieved by 130 or 260 cycles of 22 $^{\circ}\text{C}$ 35 s, 58 $^{\circ}\text{C}$ 25 s. After ACE, cells were spun and washed with 50% Formamide five times for 3 min each. 1 $\mu\text{M}$ Alexa-647 imager in 1× PBS was added to the amplified sample for 1 hour. Cells were spun and washed with 30% formamide five times for 3 min each then spun and washed with 1× PBS three times. Cells were subjected to flow cytometry to measure the signal from GFP and the signal from DNA imagers or secondary antibodies.

##### **In situ crosstalk experiments**

Based on the sequencing results, we first filtered the initiator/ssEXT library to retain sequences with >99% specificity, resulting in a final set of 58 sequences. From this set, we selected nine representative sequences: three with the lowest measured specificity, three with intermediate specificity, and three with close to 100% measured specificity. Each sequence was conjugated separately to 50 $\mu\text{L}$ of 1 mg/mL Lamin A/C antibodies. For each conjugate, HeLa cells were stained, subjected to 58-plex ACE amplification for 120 cycles, and first imaged with the cognate imager (100 nM 50 $\mu\text{L}$ imager buffer) for 1 hour to measure the on-target signal. After signal removal by 70% formamide washing, we re-imaged the same sample using a leave-one-imager-out pool containing all non-cognate imagers (100 nM each 50 $\mu\text{L}$) except the matching imager. The specific ssEXT was added as mock imager blocker at a concentration of 100 nM in the leave-one-imager-out pool. After 1 hour of incubation in imager buffer, cells were washed with 50% formamide for 6× 10 min. Intensity was then measured as the crosstalk signal.

##### **Py2T sample preparation**

For Py2T multiplex imaging, Py2T cell culture was split into two lines expressing different phenotypes, epithelial and mesenchymal. The epithelial cell line was cultured following the protocol above, and the mesenchymal cell line was cultured using this protocol but with a supplemented cell culture medium containing 4 ng/mL TGFβ (BioLegend #781802). Upon reaching confluency, each cell line was washed with 1× DPBS, treated with trypsin, centrifuged, resuspended, and passaged as outlined above. Cells were seeded onto a circular glass coverslip (bioptechs #30-1313-03192) for imaging in the FCS2 Microfluidic Chamber (Bioptechs #060319-2-03). First, one of the wells from a removable silicone 3-well chamber (Ibidi #80381) was cut off, and the resulting 2-well chamber was adhered onto the center of the glass slide. No additional surface functionalization is required to allow for sufficient cell adherence and growth. Cell lines were passaged following the above protocol until the pellet resuspension step, at which point the density of the cell suspension was determined. 20 $\mu\text{L}$ of suspension was added into 20 $\mu\text{L}$ of cell counting buffer, of which 10 $\mu\text{L}$ was added into a cell counting chip and read by the supplier cell counter. To deliver optimal density for cell segmentation, the cells were seeded at a count of 10K cells/well for epithelial cells and 15K cells/well for mesenchymal cells. 200 $\mu\text{L}$ of cell culture medium, containing 4 ng/mL TGFβ for the mesenchymal cells, was added to each well along with the calculated volume of cell suspension. The medium was then mixed via pipette aspiration, and the slide was covered and incubated at 37 $^{\circ}\text{C}$ overnight to allow cells to adhere for fixation. The rest of the protocol was described in the **cell sample preparation** section. After protein and RNA staining, the samples were ACE amplified for 180 cycles.

##### **Fresh-frozen brain preparation**

All procedures involving animals were in accordance with the US National Institutes of Health Guide for the Care and Use of Laboratory Animals. Both male and female wild-type mice (C57BL/6, 6-8 weeks, from JAX) were used in this study. Mice were transcardially perfused at room temperature using an ice-cold 4% paraformaldehyde in PBS solution. Subsequently, mouse brains were collected and immersed in 20 mL of the same fixative solution at 4 $^{\circ}\text{C}$ overnight. These fixed specimens were then placed in 100 mM glycine for 6 hours at 4 $^{\circ}\text{C}$, followed by immersion in PBS for 6-12 hours at the same temperature. Afterwards, the brains were submerged in a solution of 30% sucrose in 1× PBS at 4 $^{\circ}\text{C}$ either overnight or until they settled to the base of the container. They were then conditioned in OCT at 4 $^{\circ}\text{C}$ overnight before being flash-frozen using liquid nitrogen or dry ice. For prolonged preservation, the brains were kept at -80 $^{\circ}\text{C}$.

##### **Surface preparation and cryosection**

Briefly, round glass coverslips were cleaned with agitation by three 10-min ethanol washes, followed by three successive washes in Milli-Q water. Slides were then coated with 40% v/v of Poly-L-lysine solution (Sigma Aldrich, P8920-100ML) in Na Borate Buffer (Thermo Scientific, J62902.AK) at room temperature for 2 h on a shaker. The slides were then dried and briefly rinsed with Milli-Q water. The slides were transferred to 40% v/v of Poly-D-lysine solution (Millipore Sigma,

A-003-E) in Na Borate Buffer at room temperature overnight on a shaker. The slides were then fully dried and rinsed with Milli-Q water until visible residue was removed. Finally, the slides were dried and left in a desiccator until use. Before the brain was sectioned, poly-lysine-coated slides were equilibrated to -20 °C in the cryostat (Leica CM3050S). 10-20 µm sagittal or coronal mouse brain sections were sliced from a mouse brain block with the cryostat and mounted on the glass coverslips. The sections were quickly dried in a desiccator for 10 min and fixed by 4% PFA for 3 min, followed by 5 min incubation in 0.1 M glycine in 1× PBS. The tissues were then stored in 1× PBS with 1:500 dilution of RNase inhibitor (NEB, M0314L, 40 kU/mL) at 4 °C. To prevent RNA degradation, the tissues were used for RNA probe hybridization within 2 h after sectioning.

##### Thin section preparation

For RNA or protein staining, or <50-plex RNA + protein imaging, the staining protocol is similar to the cell sample preparation protocol mentioned above with 2× volume for each wash and protein or RNA staining. For ACE amplification, 90 µL of ACE mix containing 1× ThermoPol, 200 µM each of dATP, dTTP, and dCTP, 1 µM each ssEXT, 2 U/µL Bst large fragment DNA polymerase was added to samples. 300 µL of mineral oil was used to seal the reaction. Amplification was achieved by 200 or 300 cycles of 22 °C 1 min, 58 °C 40 s. The following imaging protocol is similar to the cell sample preparation protocol above.

For > 50-plex RNA+protein imaging on the same thin tissue, RNA and protein were stained, amplified and imaged separately. Tissue sections were first stained with RNA probes, ACE amplified, and imaged. After RNA imaging, tissue samples were digested with 100 µL RNase H + RNase I<sub>f</sub> reaction mix containing 1× ThermoPol, 2 mM DTT, 100 U/mL RNase H + RNase I<sub>f</sub> (NEB M0243S, NEB M0297L) for 2 hours at 37 °C. After aspirating the reaction mix, samples were washed 3 times with 0.1% Triton in 1× PBS at room temperature and then washed twice with 0.1% Triton in 1× PBS at 60 °C for 2 min each. After RNA digestion, tissue can be further stained, amplified, and imaged for protein targets using the protein imaging preparation protocol mentioned above.

##### Single-Cell Staining and Reproducibility Analysis

Two serial 12 µm coronal sections of PFA-perfused fresh frozen mouse brain were cut onto Superfrost Plus Gold Slides (Thermo #22-035813). Sections were thawed in a desiccator for 15 min and one of the wells from a removable silicone 3-well chamber (Ibidi #80381) was cut and adhered onto each slide. Samples were rinsed briefly with 1× PBS to remove storage residue, followed by fixation with 4% paraformaldehyde in 1× PBS for 20 min and quenching with 100 mM glycine in 1× PBS for 5 min. Samples were then blocked and permeabilized for 1 hour with 5% BSA and 0.3% Triton X-100 in 1× PBS. Antibody incubation buffer (3% BSA, 0.2% Triton, 4 mM EDTA, 0.02% dextran sulfate) was supplemented with 5 µg/mL of DNA-conjugated antibodies (NeuN, CST 36662SF; OLIG2, Biotechne AF2418) and 1 µM of each ssDNA blocker. After blocking and permeabilization, samples were incubated with antibodies overnight at 4 °C in a humidified chamber. Samples were then washed 3 × 10 min with 0.1% Triton in 1× PBS and post-fixed with 5 mM BS(PEG)5 in 1× PBS for 30 min, followed by quenching with 100 mM glycine in 1× PBS for 5 min. Samples were prepared for ACE amplification by washing 3 × 5 min with 60% formamide in 1× PBS and rinsed briefly 3× with 1× PBS. ACE reactions were run separately for each sample, as previously described with 120 cycles of 80 seconds at 40 °C and 60 seconds at 58 °C. After ACE, samples were washed 3 × 10 min with 60% formamide in 1× PBS and then briefly 3× with 1× PBS. Samples were then stained for 1 hour with 500 nM of each ACE imager in 30% formamide and 4% dextran sulfate in 1× PBS, followed by washing 3 × 5 min with 30% formamide and staining for 10 min with 1:5000 DAPI in 1× PBS. Finally, samples were rinsed briefly with 1× PBS and fixed with an imaging coverslip using silicone adhesive gel. Imaging was performed on a Nikon confocal microscope as previously described.

##### ACE with expansion microscopy

Mouse brain sections were stained with DNA-conjugated antibodies, followed by ACE reaction. After the reaction, unbound single-stranded extenders were removed using a 50% formamide in 1× PBS wash. The samples were then conditioned overnight at 37 °C with a mixture of 1 mg/mL Melphalan-X in MOPS buffer and 0.1 mg/mL AcryloylX, SE (Invitrogen A20770)<sup>16</sup>. Pre-expansion imaging was achieved by the addition of 1 µM imagers, which were later removed using a 50% formamide in 1× PBS wash. Subsequently, samples were rinsed with 1× PBS and subjected to a modified expansion microscopy procedure<sup>17</sup>. Briefly, samples were first immersed in monomer solution comprising 19% (wt/vol) sodium acrylate, 10% (wt/vol) acrylamide, 0.05% (wt/vol) N,N'-methylenebisacrylamide, 0.2% (wt/vol) APS, 0.2% (wt/vol) TEMED, and 0.01% (wt/vol) 4-HT for 20 min in an ice bath. This was followed by placing the samples between two #1 glass coverslips, with an additional coverslip on top. This assembly was then relocated to a moisture-controlled chamber and maintained at 37 °C for two hours. The expanded gel was separated from the cover glasses and submerged into a digestion buffer containing 50 mM Tris (pH 8.0), 1 mM EDTA, 0.5% Triton X-100, 0.8 M guanidine HCl, and 8

units/mL Proteinase K. After an overnight digestion, the gel underwent expansion in 0.5× PBS. For imaging of the enlarged tissue section, the gel was exposed to 1 μM imagers for an hour and rinsed 3 times with 1× PBS. Samples were imaged on a Zeiss confocal microscope.

##### **Confocal multiplexed imaging and fluidic system.**

Multiplexed Py2T confocal images were captured using a motorized Nikon Ti2 inverted microscope paired with a CSU-W1 Yokogawa spinning disk scan head with a 50 μm pinhole, an Andor Zyla 4.2 Plus sCMOS camera, a Plan Apo λ 60×/1.42 Oil DIC objective, and the Perfect Focus system v3. The Nikon Elements Acquisition Software AR 5.02 managed the microscope operations. The tissue section in FCS2 chamber was excited with 647 nm, 561 nm, 488 nm, and 405 nm lasers, utilizing a Semrock Di01-T405/488/568/647 emission filter turret. A JOBS sequence was created to facilitate overnight imaging with fluidic exchange at a 540 MHz readout rate, and a 12-bit gain of 4. At each tile, a 15 μm z-stack with 0.5 μm step-size was captured using identical settings. The image stack was automatically stitched and saved in ND2 format. Multiplexed tissue confocal images were captured using a similar setup. To increase the throughput, a Plan Apo λ 40×/1.3 Oil DIC objective was used, and we imaged each tile with a 28 μm z-stack of 7 μm step-size.

Fluid exchange was executed using a commercial automated sequential injection system (Fluigent, Aria). This system was connected to the FCS2 chamber using plastic microfluidic tubing. All washing procedures for the tissue section were conducted directly on the microscope. Initially, the tissue was washed with stripping solution (60% formamide in 1× PBS), followed by a 5-min rinse in 1× PBS, with the Aria delivering at a flow rate of 100 μL/min. To determine the appropriate focus plane, the tissue section was treated with 1 μM Round #1 imager solution and subsequently with a 4 μg/mL DAPI solution. After establishing the imaging and z-stack parameters, the Aria was activated to consecutively deliver the 1 μM imager solution, 4 μg/mL DAPI staining, 1× PBS, and a published oxygen scavenging system consisting of Trolox, PCA, and PCD<sup>18</sup>. The image capture process was initiated by transmitting a TTL signal to the microscope, which was managed by Nikon Elements Acquisition Software, following the final fluid delivery. After each imaging cycle, the Nikon Ti2 microscope dispatched a TTL signal to the Aria, prompting the subsequent buffer exchange round.

##### **Light-sheet microscopy**

Thick brain tissues were imaged using a commercial light-sheet microscope (LifeCanvas) optimized for high-throughput volumetric imaging of optically cleared specimens. The system includes a multi-immersion optical path compatible with both organic (e.g., ECI) and aqueous (e.g., 1× PBS) clearing protocols. Samples were mounted on an ASI motorized XY stage (200 mm range; 16TPI scan-optimized in the X-axis) and imaged using a 3.6× multi-immersion objective (NA 0.2, 18 mm working distance). Excitation was achieved using solid-state lasers at 405 nm (200 mW), 488 nm (150 mW), 561 nm (150 mW), and 639 nm (150 mW), with fluorescence collected through Chroma bandpass emission filters (ET445/58m, ET525/50m, ET600/50m, ET690/50m). Images were acquired with a 16-bit Hamamatsu ORCA Fusion sCMOS monochrome camera. Laser power was individually adjusted for each sample to optimize signal-to-noise performance. For stabilization during imaging, 200 μm-thick gel-embedded samples were mounted onto slides using 1% low-melting-point agarose (in 1× PBS). For 1 mm-thick tissue sections, a pre-index-matched agarose gel was applied on top of the tissue and immobilized using magnets to create a sandwich-like configuration between the slide and the gel. The slides remained stationary on the stage during image acquisition. Raw image stacks were stitched and stripe-corrected using the LifeCanvas image processing pipeline. The resulting TIFF images were then used for downstream image analysis.

##### **iDISCO+ tissue preparation**

Whole mouse brains were purchased from Jackson Laboratory, pre-fixed using standard PFA perfusion, followed by overnight incubation in 4% PFA. Samples were shipped in 1× PBS containing 0.02% NaN<sub>3</sub> on ice packs. Upon arrival, the published iDISCO+ protocol was adapted for tissue preparation<sup>19</sup>. Briefly, the brains were dehydrated and decolorized through sequential methanol/water washes (20%, 40%, 60%, 80%, and 100%), then incubated overnight in a 2:1 dichloromethane (DCM)/methanol solution at room temperature (RT) with shaking. For bleaching, samples were rinsed twice in 100% methanol (1 hour each) at RT and incubated overnight at 4 °C in 5% hydrogen peroxide (H<sub>2</sub>O<sub>2</sub>) in methanol to clear blood vessels. Rehydration was performed through sequential methanol washes (80%, 60%, 40%, and 20%) before transitioning to 1× PBS, with each step lasting 1 hour at RT. All procedures were conducted in a 15 mL Falcon tube.

##### **Universal imager preparation and hybridization**

Target-specific adapter oligonucleotides (100–350 nM each) and their corresponding universal imager oligonucleotides were mixed in 1× PBS containing 4% (w/v) dextran sulfate and 30% (v/v) formamide. Each universal imager was added at a concentration 20% higher than that of its corresponding adapter. The mixture was heated to 80 °C and cooled to 25 °C in 1 °C increments, with a 5-s hold at each temperature, to anneal the adapters and universal imagers. After cooling,

universal-imager blocking oligonucleotide was added at 30% of the adapter concentration, and the mixture was incubated at room temperature for 5 min to reduce nonspecific universal-imager binding. The resulting hybridization mixture was then applied to samples.

##### **1 mm tissue preparation**

The iDISCO+ tissue was sectioned into 1 mm slices using a vibratome (Leica VT1000S). The iDISCO+ tissue was blocked in 5% BSA, 0.3% Triton X-100, 10 µg/mL heparin, 100 µg/mL salmon sperm DNA (Thermo Fisher AM9680), and 5 µM of each antibody blocker overnight at room temperature. The DNA blockers consisted of short single-stranded DNA strands complementary to the DNA barcodes on the antibodies. The next day, each antibody was pre-hybridized with its corresponding blocker on ice for 30 min, using 7 µg antibody mixed with 1 µL of 2 mM antibody-blocker stock. Antibody mixes were then prepared by combining and concentrating all antibodies to 5 µg/mL each in 3% BSA, 0.2% Triton X-100, 4 mM EDTA, 0.02% dextran sulfate, and 5 µM antibody DNA blockers. Amicon washes were repeated four times to remove glycerol from the antibody stock solution. Prepared antibody solutions were stored at 4 °C until use.

After blocking, tissues were incubated with the antibody solution for 2 days at 37 °C or 4 days at room temperature on a nutator under controlled conditions. Following staining, tissues were washed 6 × 30 min in 1× PBS containing 0.3% Triton X-100, followed by 2 days of washing on a nutator. After washing, tissues were post-fixed with 5 mM BS(PEG)<sub>5</sub> (prepared from a 250 mM stock in DMSO and diluted in 1× PBS immediately before use) for 2 h. The fixation solution was replenished and incubated overnight. The samples were then quenched with 0.1 M glycine in 1× PBS for 1 h.

Next, tissues were washed 6 × 15 min in 30% formamide in 1× PBS, followed by three 15-min rinses in 1× PBS. Samples were then incubated with ACE solution under overnight agitation, with the ACE solution freshly replaced immediately prior to initiating the reaction. Tissues were transferred to an ibidi 2-well chamber and submerged in 1-1.5 mL of freshly prepared ACE solution. The ACE reaction was performed using 200 to 300 thermal cycles, alternating between 40 °C (3-5 min) and 58 °C (3-5 min). After the reaction, tissues were rinsed with 1× PBS and incubated at 37 °C in 60% formamide in 1× PBS, followed by additional rinsing in 1× PBS to completely remove formamide.

New imager hybridization and stripping of the previous imager were carried out in a single one-pot incubation. Briefly, 300 nM universal DNA imager was combined with 5 µM of each stripping oligo in imager incubation buffer (1× PBS, 30% formamide, 4% dextran sulfate). Samples were incubated at 55 °C with shaking for 2 h, followed by an additional 2 h at 37 °C. After incubation, tissues were washed eight times with 30% formamide in 1× PBS at 37 °C for 15 min per wash. Tissues were then cooled to 4 °C in 1× PBS and dehydrated in methanol three times (5–10 min each) at 4 °C. For index matching, samples were incubated in ethyl cinnamate (ECi) on a rotator or nutator for 30–45 min with three ECi replenishments. A ~20% reduction in linear dimensions was observed for 1 mm tissue samples following ECi-based index matching. After imaging for 20 min (MegaSPIM with 3.6× objective, half-coronal section), tissues were briefly rinsed in methanol, incubated in methanol for 15 min, and rehydrated in 1× PBS for 10–30 min with two PBS exchanges. Imager stripping and hybridization were then repeated as described above.

##### **200 µm tissue preparation**

The iDISCO+ tissue was sectioned into 200 µm slices using a vibratome (Leica VT1000S). Blocking was performed in 5% BSA and 0.3% Triton X-100 for 2 hours at room temperature. Simultaneously, the antibody solution was prepared by premixing antibodies at a final concentration of 10 µg/mL in 3% BSA, 0.2% Triton X-100, 4 mM EDTA, 0.02% dextran sulfate, and 500 nM antibody-specific DNA blockers. The antibody solution was stored at 4 °C until use. Following the blocking step, the tissue was incubated with the antibody solution for 24 hours at 37 °C. After incubation, the tissue underwent six sequential 15 min washes in 1× PBS containing 0.3% Triton X-100. Post-washing, the tissue was post-fixed with 5 mM BS(PEG)<sub>5</sub> for 2 hours. The fixation was followed by quenching with 0.1 M glycine in 1× PBS for at least 30 min. Next, the tissue was washed six times with 30% formamide in 1× PBS for 30 min per wash and subsequently rinsed with 1× PBS. The sample was then incubated overnight with the ACE solution under agitation. The ACE solution was replenished, and the reaction was carried out for 260 cycles in a 2-well ibidi glass-bottom chamber using a thermal cycler. Meanwhile, the Melphalan-X solution was prepared by incubating 50 µL of Acryloyl-X (10 mg/mL) and 200 µL of Melphalan (2.5 mg/mL) with overnight shaking. The tissue was then incubated overnight at 37 °C in a mixture containing 250 µL Melphalan-X, 25 µL 1× PBS, 10 µL MOPS, 215 µL H<sub>2</sub>O, and 5 µL Acryloyl-X (10 mg/mL). Following incubation, the brain samples were washed six times with 1× PBS (10 min per wash) and incubated on ice with a gelling solution composed of 95.3 µL 40% acrylamide, 100 µL 2% bis-acrylamide, 100 µL 10× PBS, and 664.7 µL H<sub>2</sub>O for 30 min. During incubation, 10% APS and 10% TEMED were prepared separately and kept on ice. The tissue was then transferred onto glass slides, and excess solution was wicked away using a Kimwipe. A fresh gel mixture (100 µL of gelling solution, 2 µL

TEMED, and 2 μL APS) was applied over the sample, with ∼50 μL of the solution dropped onto the tissue, which was then sandwiched with a coverslip. The polymerization reaction was carried out in a humidified chamber at room temperature for at least 30 min. After polymerization, the gel-embedded tissue was trimmed to the appropriate size on the glass slides and rinsed with 1× PBS. Tissue digestion was carried out overnight at 37 $^{\circ}$C using proteinase K (1:50 dilution in 1× PBS containing 0.5% Triton X-100 and 2% SDS). The tissue was then rinsed with 1× PBS and subjected to a 2nd gelling step to reinforce the hydrogel structure. Following the second gelling, the tissue was washed with 1× PBS, resulting in a slight expansion (∼1.5×). Finally, the gel was stored at 4 $^{\circ}$C until further use.

The tissue gel was rinsed with 1× PBS and incubated for 2 hours in a 250 nM DNA universal imager solution (1× PBS, 30% formamide, 4% dextran sulfate). After incubation, POPO-1 solution (Invitrogen P3580, 1:5000 dilution in 1× PBS, 30% formamide) was applied to the tissue gel for 1 hour, followed by either an overnight wash in 1× PBS or six 30-min washes at room temperature. The tissue was then mounted on slides by embedding in 1% low-melting-point agarose (prepared in 1× PBS and stored at 40 $^{\circ}$C) for stabilization. Imaging was performed using a light-sheet microscope. To remove the imaging signal, the tissue was incubated overnight at 37 $^{\circ}$C with agitation in 70% formamide in 1× PBS. The sample was subsequently rinsed with 1× PBS, and the imager solution was reapplied following the same protocol described above.

##### **Immuno-SABER in 1 mm mouse brain section**

For Figure S19a data set, immuno-SABER imaging of a 1 mm SHIELD-processed mouse brain section was prepared following a published protocol<sup>1</sup> using commercial reagents from LifeCanvas. For Figure S19a and Figures S25–S28 data sets, 1 mm iDISCO+ tissues were prepared using an adapted version of the published iDISCO+ protocol<sup>19</sup>. Then, the tissues were blocked with 5% BSA and 0.3% Triton X-100 for 1 day at room temperature. Then DNA-barcoded antibody was added at 5 μg/mL in 3% BSA, 0.2% Triton X-100 with 4 mM EDTA, 0.02% dextran sulfate, 5 μM of each antibody blocker strand, and subsequently incubated for 4 days with shaking at room temperature. Post-staining, the tissues were washed with 1× PBS+0.3% Triton X-100 for 2 days. Following this, a post-fixation was done using 5 mM BS(PEG)<sub>5</sub> overnight, followed by a quenching process with 0.1 M glycine in 1× PBS for 1 h. The tissues were rinsed in 1× PBS and incubated 1 day with pre-synthesized SABER concatemer (600-650 nt long) at a concentration of 150 nM in the SABER concatemer hybridization buffer. After concatemer hybridization, the tissues were washed with 50% formamide in PBS for 30 min, repeated six times. They were then rinsed with 1× PBS and incubated with 1 μM DNA imager in 1× PBS overnight, rinsed again with 1× PBS for 30 min (repeated three times), and then made ready for imaging.

##### **HCR amplification in 1 mm mouse brain section**

Immuno-HCR imaging in a 1 mm iDISCO-treated mouse brain section was performed using HCR<sup>TM</sup> Gold 2<sup>o</sup>IF reagents from Molecular Instruments Inc., with minimal modifications for thick cleared tissue. Briefly, a 1 mm iDISCO tissue section was prepared using the published iDISCO+ protocol. The tissue was blocked with 5% BSA and 0.3% Triton X-100 in 1× PBS for 1 day at 37 $^{\circ}$C with shaking. Primary antibody was diluted in 3% BSA and 0.2% Triton X-100 in 1× PBS at 5 μg in 500 μL and incubated with the tissue for 2 days in a 37 $^{\circ}$C shaker. After primary antibody incubation, the tissue was washed in 1× PBS containing 0.3% Triton X-100 and 10 μg/mL heparin at 37 $^{\circ}$C, first for 30 min repeated six times followed by overnight washing, and then for 1 hour repeated three times followed by overnight washing.

The tissue was then incubated with HCR<sup>TM</sup> 2<sup>o</sup> Antibody Probe at 10 μg/mL in 1 mL antibody incubation buffer in a 37 $^{\circ}$C shaker. Following HCR<sup>TM</sup> 2<sup>o</sup> Antibody Probe incubation, the tissue was washed with 1× PBS containing 0.3% Triton X-100 for 30 min repeated six times, followed by overnight washing. The tissue was post-fixed using BS(PEG) for 2–5 hours followed by overnight incubation, quenched, and washed with 1× PBS.

For HCR amplification, HCR<sup>TM</sup> Gold Amplifier hairpins were prepared according to the Molecular Instruments HCR<sup>TM</sup> Gold 2<sup>o</sup>IF protocol and incubated with the tissue in HCR<sup>TM</sup> Gold Amplifier Buffer overnight at room temperature, protected from light. After amplification, excess amplifier was removed by washing, and the sample was prepared for imaging. For tertiary antibody labeling, the tissue was imaged and then incubated with tertiary antibody at 40 μg/mL in 1 mL buffer. The tissue was again washed with 1× PBS containing 0.3% Triton X-100 for 30 min repeated six times followed by overnight washing, post-fixed with BS(PEG) for 2–5 hours followed by overnight incubation, quenched, and washed with 1× PBS before further imaging.

##### **RCA amplification in 1 mm mouse brain section**

RCA amplification in a 1 mm iDISCO-treated mouse brain section was performed using DNA-barcoded primary antibody labeling followed by padlock-based rolling circle amplification. Briefly, a 1 mm iDISCO tissue section was prepared using the published iDISCO+ protocol. The tissue was blocked with 5% BSA and 0.3% Triton X-100 in 1× PBS for 1 day at 37 °C with shaking. Primary antibodies were conjugated to an azide-modified DNA barcode sequence, 5'-Azide-TTTTTTTTTTTTTTTTAGACACCGTTCCTTGGACAGAGC-3', and diluted in 3% BSA and 0.2% Triton X-100 in 1× PBS. The DNA-barcoded primary antibody was incubated with the tissue in a 37 °C shaker for 2 days. After antibody incubation, the tissue was washed in 1× PBS containing 0.3% Triton X-100 and 10 µg/mL heparin at 37 °C, first for 30 min repeated six times followed by overnight washing, and then for 1 h repeated three times followed by overnight washing.

After antibody post-fixation, tissue slices were incubated with padlock oligos (5'-/5Phos/GAACGGTGTCTATTATATCCACCATTCCCACCATACCCACCATCTTATATGCTCTGTCCAAG-3', 300 nM) in the hybridization buffer at 37 °C with gentle shaking for 24 h. The base composition of the hybridization buffer included 2× SSC, 1% Triton X-100, 10% small dextran sulfate. After incubation, the samples were washed four times for 1 h each with PBST and were incubated in ligation mixture (0.25 U µl<sup>-1</sup> T4 DNA ligase, 1:100 BSA) at 4 °C for 5 h with gentle shaking and then incubated at 22 °C for 14 h. Then the sample was washed with PBST three times for 1 h each. After washing, the sample was incubated with RCA mixture (0.5 U µl<sup>-1</sup> Phi29 DNA polymerase, 1 mM dNTP, 1:100 BSA in 1× Phi29 buffer) at 4 °C for 24 h for equilibration before incubating at 30 °C for 8 h for amplification and washed with PBST three times for 1 h each. Then, the sample was incubated with 300 nM imager in 30% formamide, 1× PBS buffer for 8 h. After detection probe hybridization, the tissue was washed with 1× PBS and prepared for fluorescence imaging.

##### **Formalin-fixed, paraffin-embedded (FFPE) sample preparation**

Human specimens were obtained from the archives of the Department of Pathology at Beth Israel Deaconess Medical Center under a discarded and excess tissue protocol approved by the Institutional Review Board (IRB; Protocol 2017P000585). All relevant ethical regulations were followed. FFPE tissue blocks were sectioned at a thickness of 5 µm using a rotary microtome. Sections were floated on a 30 °C water bath, mounted onto positively charged glass slides, and baked at 60 °C for 2 h to promote tissue adhesion. Deparaffinization, rehydration, and heat-induced epitope retrieval were performed in citrate buffer using a PT Link system (Agilent). Following antigen retrieval, the sections were washed and stored in PBS at 4 °C for up to 1 week before staining; storage for longer than 1 week may reduce antigenicity. Immediately before staining, the sections were washed in PBS for 15 min, and the tissue area was either outlined with a hydrophobic barrier pen or enclosed within a removable incubation chamber (Ibidi, catalog no. 80381). Antibody staining, ACE amplification, imager-probe hybridization, and DAPI staining were subsequently performed using the corresponding procedures described for cultured-cell samples.

##### **Data analysis and statistics**

###### **Gel-image analysis and product-length quantification**

Gel images were converted to grayscale and analyzed independently. Lane centers were obtained from the physical lane map. Lane half-width was set to 18% of the mean lane spacing and constrained to 8–15 pixels. A one-dimensional migration profile was generated by averaging fluorescence intensity across the width of each lane. A migration-dependent spatial background was estimated from predefined lane-free corridors on each gel. For each image row and corridor, pixels exceeding the corridor median plus 3.5 times the median-absolute-deviation scale estimate were rejected. The retained row-wise medians were smoothed along the migration axis using a Gaussian kernel ($\sigma = 8$ pixels). Corridor-specific background curves were then linearly interpolated across the horizontal position of the gel to obtain a migration-dependent background curve for each lane. Residual low-size background was corrected using the same rule for every included lane. Two 12-pixel side windows, separated from the lane edge by 6 pixels, were summarized by their row-wise medians and smoothed using a Gaussian kernel ($\sigma = 10$ pixels). The lower of the two side curves was used as conservative bilateral background support, minimizing the influence of a bright neighboring lane or a one-sided image artifact. Only positive background exceeding the spatially interpolated curve was added. For each initiator–extender pair, the fluorescence-weighted mode, mean, median, 25th and 75th percentiles, and the fluorescence fractions within 100–199, 200–399, 400–599, 600–799 and 800–900 nt were calculated from the normalized distribution. The violin plot represents the distribution of the 58 per-sample median lengths, with individual samples shown as points and the median across samples shown as a horizontal line. In the per-sample box plot, boxes extend from the 25th to the 75th percentile, center lines denote medians and whiskers extend from the 5th to the 95th percentile. The box-plot axis was displayed on a log<sub>2</sub> scale from 100 to 800 nt, whereas the underlying distributions were quantified over the complete 100–900 nt interval.

###### **RNA intensity quantification**

We labeled the APC gene with two sets of probes in HeLa cells. The reference probe set contained 65 probes, and the ACE probe set contained 40 probes, each with the same ACE initiator. After ACE amplification and imager incubation, we imaged the cells subjected to different numbers of amplification cycles using 35 z-planes with a 0.3-$\mu\text{m}$ step size. We first created a maximum z projection of the images and subtracted the background intensity from the pixel intensity. We used puncta imaged from the reference probes as references and isolated the overlapped puncta from the ACE imager channel. We then quantified the maximum intensity of pixels in the puncta to compare signal amplification for different time points.

##### **Protein quantification**

We labeled the Lamin-B protein with 400 ng/mL ACE initiator-conjugated anti-Lamin-B antibodies in HeLa cells overnight at 4 °C. After ACE amplification and imager incubation or secondary Ab with the same fluorophore staining for 2 h, we imaged the cells subjected to different numbers of amplification cycles using 23 z-planes with a 0.7-$\mu\text{m}$ step size. We first created a maximum z projection of the images and subtracted the background intensity from the pixel intensity. We then used the DAPI channel to create masks for the cell nuclei and measured the mean intensity of the DNA imager signal or secondary Ab signal for each nucleus. We then quantified the distribution of intensity for different time points.

##### **RNA Puncta Co-localization Analysis**

Single-molecule RNA transcripts were quantified and analyzed on 2D Maximum Intensity Projections (MIP) generated from 3D ND2 image stacks for the Cy5, TRITC, and DAPI channels. To segment nuclei, DAPI MIP images were smoothed with a Gaussian filter ($\sigma=3.0$), segmented using Otsu's thresholding method, closed morphologically using a disk-shaped structuring element, and filtered to remove small noise objects. For RNA spot detection, a Difference of Gaussians (DoG) filter (with standard deviation multipliers $\sigma_1=1.0$ pixels and $\sigma_2=2.0$ pixels) was applied to the Cy5 and TRITC MIP images to enhance puncta signals. To ensure robust thresholding in the presence of hot pixels or saturated regions, a trimmed standard deviation was computed by clipping values above the 99.9th percentile. Puncta were identified as local intensity maxima in the DoG-filtered image. For ACE-amplified samples, a raw intensity cutoff of $\geq 2250$ was applied to both channels, while non-amplified smFISH samples were filtered with a raw intensity cutoff of $\geq 600$. Co-localization between Cy5 and TRITC channels was computed using a standard greedy 1-to-1 matching algorithm. Puncta were classified as overlapping if their Euclidean distance in physical space was $\leq 0.35 \mu\text{m}$.

##### **SOX10 ACE vs secondary Ab**

To evaluate the quantitative agreement between the secondary antibody and the ACE imager signals for the target marker SOX10, signal correlation analysis was performed at single-cell resolution. Single-cell nuclear segmentation was performed on the DAPI channel by applying a Gaussian smoothing filter, binarizing via Otsu's thresholding, and resolving touching nuclei using a distance transform-based watershed segmentation. Segmented nuclear masks were size-filtered to exclude debris and multicellular aggregates. To identify SOX10-positive cells, local background subtraction was performed on both the secondary antibody and ACE imager channels using a morphological white top-hat filter. Individual cells were classified as SOX10-positive if their mean background-subtracted nuclear intensity exceeded the Otsu threshold calculated across the nuclear population in either channel. For the identified positive cell population, raw (unsubtracted) mean nuclear intensities were extracted from both channels. The signal co-localization and intensity scaling were quantified by calculating Pearson's linear correlation coefficient. All image processing, segmentation, and statistical analyses were performed in Python using the nd2, scikit-image, and scipy packages.

##### **EGFR KO analysis**

To quantify cellular EGFR expression, single-cell boundaries were reconstructed using DAPI-labeled nuclear channels as seeds. Briefly, nuclei were segmented using Otsu's thresholding, filtered to remove small noise artifacts (area < 100 pixels), and separated using a distance-transform-seeded watershed algorithm. Approximate cell boundaries (encompassing both cytoplasm and plasma membrane) were defined by dilating the segmented nuclear masks by 15 pixels, with adjacent cell boundaries partitioned via a flat-image watershed constraint to prevent spatial overlap. To perform local background subtraction, a non-cell background mask was generated by inverting a 35-pixel dilation of the nuclear masks. For each field of view and each fluorescence channel, 10,000 pixels were randomly sampled within this non-cell area to calculate the local median background intensity and background noise (standard deviation). The channel- and image-specific median background was then subtracted from the mean raw intensity of each segmented cell. Target specificity was evaluated as the ratio of average background-subtracted cell intensities in wild-type (WT) cells to those in knockout (KO) cells.

##### **Loss of signal analysis**

Image registration, segmentation, and intensity profiling were performed using a custom Python pipeline (based on scikit-image and scipy). To correct for inter-cycle stage drift, translation shifts were calculated relative to Round 1 using a hybrid DAPI alignment strategy: a coarse grid search was performed on 4-fold downsampled, Gaussian-filtered ($\sigma=2$) DAPI

images to resolve large shifts, followed by high-resolution phase cross-correlation (upsampling factor of 10) on Gaussian-filtered ($\sigma=1$) DAPI images to refine the sub-pixel alignment. To profile independent nuclear properties, DAPI images were segmented using local background subtraction, Gaussian smoothing ($\sigma=2$), Otsu thresholding, and distance-transform-seeded watershed segmentation. For cell-by-cell tracking across the 20 cycles, reference nuclei from Round 1 were filtered to exclude boundary cells (within 30 pixels of the image borders). The reference segmentation mask was then dynamically translated to each subsequent round using the calculated DAPI registration shifts via nearest-neighbor interpolation (order=0) to preserve label identities. Mean and integrated Lamin A/C channel intensities were extracted from the registered images, and log2-transformed intensities and intensity ratios normalized to Round 1 were calculated.

##### **Inter-run analysis**

Cell nuclei were segmented from the DAPI channel of Nikon ND2 raw images by applying Gaussian smoothing ($\sigma=1.5$), local adaptive thresholding (51-pixel block size), and distance-transform-guided watershed segmentation to separate touching cells. For each detected cell, mean fluorescence intensities were extracted from the nuclear mask and a corresponding cytoplasmic mask (defined by a 5-pixel dilation of the nuclear boundary, excluding the nucleus). Cells were classified as positive for NeuN ($\geq 1100$, nuclear), and OLIG2 ($\geq 1200$, nuclear) based on signal-to-background profile distributions. Staining reproducibility was evaluated across independent runs by comparing median intensities of positive cells.

##### **Anti-GFP Image Analysis and Puncta Quantification**

To quantify the anti-GFP antibody signal, multi-channel images (DAPI, GFP, and 640 nm antibody channels) were processed using a custom Python pipeline based on scikit-image. Nuclei were segmented from the DAPI channel. Cell boundaries were defined by expanding the nuclear masks outward up to 27.6 $\mu\text{m}$ (300 pixels) using a Voronoi-like label expansion algorithm. The transfected cell population was isolated by classifying cells based on their mean GFP (488 nm) intensity, including only cells exceeding a threshold of 950 a.u. (GFP-High) and excluding non-transfected ($<850$ a.u.) or intermediate cells. To isolate localized puncta from diffuse background staining, a white top-hat filter (disk structuring element, radius = 6 pixels) was applied to the 640 nm antibody channel. To ensure an unbiased comparison between positive and negative control (NC) images, a positive-calibrated thresholding strategy was employed: three positive images were randomly selected per assay, the peak intensities of the top 20 candidate puncta (above a noise floor of 30 a.u.) were measured, and the assay-specific threshold was set at 50% of the overall median of these pooled peaks. These fixed calibrated thresholds were applied uniformly to all positive and negative control images in their respective groups. The puncta intensity per cell was calculated as the sum of pixel intensities in the white top-hat filtered 640 nm channel within the segmented cell boundary that overlapped with the detected puncta mask. For statistical analysis, the mean puncta intensity was averaged across all transfected cells within each image to treat the image as the independent unit of biological replication ($n=8-16$ images per group).

##### ***In situ crosstalk experiments***

To quantify Lamin antibody signal-to-noise ratio (SNR) across barcodes, Maximum Intensity Projections (MIP) along the Z-axis were calculated for both the specific imager and nonspecific all-imagers conditions. Spatial drift between imaging sessions was corrected via DAPI-to-DAPI alignment using Phase Cross-Correlation, applying a well-average translation shift fallback for failed field-of-view registrations. Cell nuclei were segmented on the reference DAPI channel using Otsu thresholding followed by distance-transform-seeded watershed segmentation. To subtract background noise, Well 10 (Negative Control, no antibody) nuclear masks were used to estimate the average background pixel intensities for both imaging conditions; these background levels were subtracted pixel-wise from the masks of Wells 1 to 9, clipping negative values at zero. Finally, the single-cell Sum SNR was calculated for each segmented nucleus as the ratio of the integrated background-subtracted specific imager intensity (signal sum) to the integrated background-subtracted all-imagers intensity (noise sum).

##### **Py2T cell analysis**

After each round of imaging, fields of view were first stitched using Nikon software. We then created maximum z-projections for the stitched images and aligned different rounds of images using the DAPI channel. After image alignment, we performed cell segmentation using Python package `napari_segment_blobs_and_things_with_membranes`<sup>20</sup> and `pyclesperanto_prototype`. PolyA signal and DAPI signal were used for the segmentation. We subtracted the background using images taken from the sample where no imagers were added. For RNA targets, we thresholded the images to remove the signal from nonspecific probe binding. For most RNA targets except for Krt8 and Fn1 where puncta are too dense to resolve, we quantified the number of puncta for each segmented cell. For protein targets and RNA targets Krt8 and Fn1, we quantified the intensity of all pixels in each cell. We then size-normalized the targets to reduce the effect of cell size.

We then scaled the expression level for each target and performed Leiden clustering using the Python package scanpy<sup>21</sup>. We performed heatmap analysis of clusters using the seaborn package<sup>22</sup> and UMAP analysis using the umap package<sup>23</sup>. Pseudotime analysis was performed using the built-in diffusion pseudotime analysis of the scanpy package. We used the matplotlib package<sup>24</sup> to plot most of the results.

##### Mouse brain thin section data analysis

The mouse brain thin section cell segmentation and background subtraction were similar to the Py2T cell pipeline. We used median blur and Voronoi analysis to reduce nonspecific signals for RNA targets. For all RNA targets and protein targets, we quantified the intensity of all pixels in each cell and performed size normalization and scaling. The clustering and embedding were similar to the Py2T cell pipeline. To perform neighborhood analysis, we used the squidpy package<sup>25</sup>. To plot the chord diagrams showing the spatial proximity, we used the pyCirclize package<sup>26</sup>.

##### Thick tissue data analysis and alignment

We imaged a thick section of the mouse brain over multiple sequential imaging rounds, labeling Histone H3 (for 1mm coronal section) or POPO-1 (for 200 μm sagittal section) in every round to serve as a fiducial channel. All Histone H3 or POPO-1 channels from round 2 onward were mapped to the corresponding channel in round 1. Specifically, we first cropped the 3D image to remove white space. We then wrote the images to chunked Zarr arrays ($64 \times 64 \times 64$ voxels) for parallel I/O. Blob-like Histone H3/POPO-1 signals were detected in the pre-processed volumes using the Bigstream Blob detection function<sup>16</sup>, and their centroids were exported as point clouds. Coregistration was performed in 3-D with the Phathom package<sup>27</sup>: (i) geometric features were computed for the fixed (round 1) and moving point clouds; (ii) putative correspondences were identified by correlation-based neighborhood search and filtered by distance and coherence; (iii) an affine transform was estimated, followed by iterative non-rigid refinement with B-spline regularization over 40 rounds using $\sim 100$ nearest-neighbor matches. The final coordinate interpolator was applied to resample every channel of each round into the reference volume, producing fully coregistered datasets. Alignment accuracy was verified by visual overlay in ImageJ and by quantification of residual distance of matched point pairs of successive rounds.

##### 3D image processing

To facilitate efficient data processing and 3D rendering, images acquired from 200 μm sagittal sections were downsampled twofold. Background signals were removed using the rolling ball subtraction algorithm in ImageJ prior to format conversion. TIFF images from both 200 μm and 1 mm tissue sections were then converted to Imaris-compatible format (.ims) and visualized using Imaris version 10.2.0. Anatomical subregions were manually defined using the “Surface” tool, guided by reference atlases from the Allen Brain Atlas<sup>28</sup>. Cell coordinates were measured using the “Spot” module in Imaris or with our customized blob-detection algorithms in Python. Spatial coordinates extracted from either method were used for downstream neighborhood analysis. For cells initially detected using the Python-based blob-detection pipeline, coordinates were subsequently confirmed and refined in Imaris using the “Spot” module to improve centroid accuracy and to calculate distances from each spot to anatomical subregions defined by the “Surface” tool. For vasculature analysis, surfaces representing large vessels and capillaries were generated using the “Surface” module based on CD31, VIM, and GLUT1 immunostaining. Overlapping volumes between vascular structures were quantified using the “Overlapped Volume” function in Imaris. 3D visualizations and videos were generated using the “Animation” and “Snapshot” modules.

##### Neighborhood analysis

We performed a 3D radial distribution function (RDF) analysis to quantify spatial enrichment between cell types based on immunofluorescence-stained cell coordinates. For each marker pair, we calculated the number of type B cells within concentric spherical shells surrounding type A cells using a cKDTree. For the 1 mm section, we set the radius to be 10 μm with bin size = 3. For the 200 μm section, we set radius to be 15 μm with bin size = 5 to account for expansion of the hydrogel during imaging. These counts were normalized by expected values assuming a random distribution, yielding an RDF curve $g(r)$. The maximum $g(r)$ value was used as the enrichment score. Pairs with fewer than 15 proximity events were excluded. The results were visualized as a heatmap of marker–marker enrichment scores.

##### Z bin quantification

To quantitatively evaluate the uniformity of signal amplification across the Z-depth of cleared tissue volumes, relative fluorescence staining intensities were profiled across a continuous 1 mm thickness window spanning from slice 73 to slice 671 (corresponding to a calibrated physical depth of 127.75 μm to 1174.25 μm). Profile extraction was executed for all 16 evaluated targets, comprising 14 discrete spot-detection channels (including NeuN) and 2 volumetric vessel masks (CD31 and GLUT1). The active tissue volume was discretized along the Z-axis into sequential, non-overlapping 20 μm Z-bins. For each target, localized descriptive statistics—including the mean intensity ($\mu_{\text{bin}}$), standard deviation ($\text{STD}_{\text{bin}}$), and standard error of the mean ($SEM_{bin}$)—were calculated within individual spatial bins. To control for varying baseline expression and absolute intensity scales across heterogeneous targets and channels, raw intensity values within each individual target profile were normalized against the global average baseline.

##### **Volumetric Signal Accuracy and Intensity Correlation Analysis**

To assess the quantitative accuracy and spatial preservation of target protein intensities following signal amplification steps, we evaluated the voxel-level signal correlation between ACE and the secondary Ab. Analysis was performed across the 3D volume at five representative optical depths centered at $Z = 200, 300, 400, 500$, and $600$ slices. At each depth center, a local Maximum Intensity Projection (MIP) was generated using a 21-slice window ($\pm 10$ slices) to capture overlapping biological features while minimizing depth-dependent registration noise. Spatial anchor locations were defined using 3D structural landmarks (blobs) extracted via a parallelized difference-of-Gaussians (DoG) detector on the reference volume, filtered with an intensity cutoff threshold of 500 to ensure high-confidence spot selection. To mitigate pixel-level signal fluctuations and account for sub-voxel registration tolerances, the final local intensity was computed as the mathematical mean within a dilated 2D spatial footprint radius of 3 pixels ($7 \times 7$-pixel neighborhood grid) around each blob center. The linear correlation and intensity accuracy across depths were quantified using the Pearson correlation coefficient ($r$), paired with ordinary least-squares (OLS) linear regression to map trend lines across the individual imaging windows.

##### **Depth-Resolved Intensity Profile Analysis**

To quantify signal distribution and depth penetration across the $Z$-axis, 3D Zarr-formatted image volumes were loaded and processed using Dask. For each target, a centered regional subvolume (ROI) corresponding to 25% of the total field of view in both $X$ and $Y$ (25% center-crop in $X \times 25\%$ center-crop in $Y$) was extracted. Raw fluorescence intensities were summed across both spatial dimensions ($X$ and $Y$) at each discrete $Z$ plane. To account for wide dynamic ranges, $\log_2$-transformed profiles ($\log_2[\text{Intensity}(Z) + \epsilon]$ with $\epsilon = 10^{-6}$) were computed. Depth values in micrometers were calculated using a voxel size of $1.75 \mu\text{m}$ along the $Z$-axis.

##### **Volumetric Cell Detection and Spatial Depth Analysis**

To evaluate the uniformity of spot counts across tissue depth, coordinates were binned along the $Z$-axis into discrete 60-slice intervals ($\Delta Z = 105 \mu\text{m}$ given a $1.75 \mu\text{m}/Z$-voxel step size) over a total depth range of 725 slices ($1,268.75 \mu\text{m}$). For each bin $i$ bounded by $[Z_{start}, Z_{end}]$, the absolute number of detected spots $N(Z_i)$ was computed as the sum of all spots falling within the specified $Z$-slice boundaries. The bin center coordinate in physical distance ($\mu\text{m}$) was defined as $Z_{center} = ((Z_{start} + Z_{end} - 1) / 2) * s_z$, where $s_z = 1.75 \mu\text{m}$ denotes the axial voxel spacing. Statistical similarity between channel spatial distributions was quantified using two-tailed Pearson's correlation coefficient ($r$).
